# In sickness and in health: RNA-Seq finds viruses associated with mutualist quality in the Amazonian plant-ant *Allomerus octoarticulatus*

**DOI:** 10.1101/2024.03.12.583798

**Authors:** María C. Tocora, Christopher Reid, Jesse Huisken, Haoran Xue, Megan E. Frederickson

## Abstract

Ant-plant symbioses are classic examples of mutualism in which ant “bodyguards” defend myrmecophytic plants against enemies in exchange for nest sites and often food. We used RNA-Seq to profile the transcriptomes of *Allomerus octoarticulatus* ant workers, which aggressively defend the Amazonian plant *Cordia nodosa* against herbivores, but to varying degrees. Field behavioral assays with herbivores in the Peruvian Amazon showed striking variation among colonies in the relative zeal with which *A. octoarticulatus* workers defend their host plant. Highly effective and ineffective bodyguards differed in their gene expression profiles, which revealed viral infections significantly associated with ant bodyguarding behavior. Transcripts from eight new positive-sense single-stranded RNA viruses were differentially expressed between colonies with high- or low-quality bodyguards. Colonies of high- and low-quality bodyguards were infected by distinct viruses, including viruses clustering phylogenetically with viruses known to cause aggression or reduced locomotion, respectively, in bees. Gene expression, including of immunity-related genes, also differed between broodcare workers and bodyguard ants, suggesting bodyguarding is a distinct worker task. Ant colony health and viral infections may influence ant cooperation with plants in ant-plant mutualisms.

**Significance statement:** Many animals have evolved to cooperate with plants in plant-animal mutualisms, including in pollination, seed dispersal, and defensive ant-plant interactions. Although the ecology of these interactions is well studied, little is known about the molecular mechanisms involved in animal cooperation with plants. Studying an Amazonian ant-plant symbiosis, we combined field assays of ant behavior with RNA-Seq to profile the transcriptomes of ants that vary in their quality as defensive plant “bodyguards.” Bodyguard ants that effectively defended plants against herbivores expressed different genes than both ineffective bodyguards and broodcare workers. Surprisingly, many genes differentially expressed between high- and low-quality bodyguards were from viruses, and not ants. Thus, we linked viral infections to mutualist quality in this ant-plant symbiosis.

## Introduction

Most plants and animals are involved in at least one mutualism; they depend on other species for nutrition, protection, or dispersal (Bronstein, 2015). Yet how mutualism evolves remains poorly understood, especially in insects, as we know little about the genes that encode mutualistic behaviors, or how these behaviors are regulated at the molecular level and evolve in insect genomes. In insects and many other animals, cooperation is typically a behavioral trait, and many studies have explored the molecular basis of social behavior (i.e., within-species cooperation) in insects such as ants (Kohlmeier et al., 2019; Manfredini et al., 2013) and bees (Jones et al., 2020). In contrast, very little is known about the molecular mechanisms that give rise to cooperative animal behaviors directed at other species in mutualisms. Here, we take a transcriptomic approach to understand the defensive “bodyguard” behavior of symbiotic ants that defend their myrmecophytic hosts against herbivores.

Many theoretical models explore mutualism evolution, often with the goal of explaining how mutualism originates and persists despite the potential for underlying fitness conflicts between partners (Sachs & Simms, 2006). Nonetheless, in plants and microbes, where research has made great strides towards unraveling the “molecular dialogue” between partners, the evolution of genes known to encode mutualistic traits does not always match theoretical predictions (e.g., Epstein et al., 2023). For example, theory predicts that fitness conflicts select for “cheaters” that drive the breakdown or dissolution of mutualisms, but cheating is unlikely to be an important source of variation in mutualist quality in many systems (Frederickson, 2013). Instead, symbionts often vary in effectiveness because they are mismatched to their host’s genotype (Batstone et al., 2020) or environment (Kiers et al., 2007). Failure to cooperate might also occur because of stress or illness. Disease might trigger a decrease in mutualistic behavior, making symbiotic partners “defective, but not defectors” (Friesen, 2012; Riehl & Frederickson, 2016). Although many studies have explored how pathogens modulate host behavior to increase parasite fitness and transmission (de Bekker, et al., 2018), how parasites or pathogens impact the beneficial behaviors of mutualists has received little attention outside of pollination mutualisms (Gillespie & Adler, 2013).

Approximately 4,000 species of angiosperms interact mutualistically with ants (Weber & Keeler, 2013). These associations are classic examples of defensive mutualisms, in which plants provide food or housing to “bodyguard” ants that defend them against herbivores or other enemies. Macroevolutionary analyses suggest that ant-plant mutualisms spurred plant diversification (Weber & Agrawal, 2014). However, despite ample ecological study (Mayer et al., 2014), few studies have applied ‘omics approaches to ant-plant interactions. On the plant side, metabolomic and transcriptomic comparisons have assessed the effects of ant colonization on *Tococa* ant-plants (Müller et al., 2022). On the ant side, Malé et al. (2017) suggested that the cGMP-PKG signaling pathway modulates ant protection of plants, while Rubin and Moreau (2016) found higher rates of molecular evolution genome-wide in mutualistic acacia-ants compared to their non-mutualistic close relatives, but there has been little additional research.

Here, we studied the mutualism between the ant species *Allomerus octoarticulatus* and the Amazonian myrmecophyte *Cordia nodosa* (Fig. 1A). *Cordia nodosa* has hollow stem swellings called domatia where ants nest, with a single *A. octoarticulatus* colony nesting across many domatia on a single *C. nodosa* tree. We combined bioassays in the field with transcriptomics to unravel the molecular mechanisms underlying mutualistic outcomes. We first characterized bodyguard behavior in ant colonies in the Peruvian Amazon by placing herbivores (grasshoppers) on plants and monitoring worker recruitment (Supplementary Video 1 and Fig. 1). Most measured variables, including herbivore size, age, and species (Supplementary Fig. 1), did not significantly affect worker recruitment, but ant activity in these bioassays did significantly predict herbivory on young *C. nodosa* leaves. We then identified transcriptional signatures of mutualist quality by analyzing differential gene expression between high- and low-quality ant bodyguards. We also used RNA-Seq on broodcare workers and bodyguards to identify task-specific transcripts that vary between ants working inside and outside the nest. Our results show that highly effective ant bodyguards express different genes than both ineffective ant bodyguards in other colonies and broodcare workers from the same nest. We also found underlying viral infections and variation in the expression of immunity-related genes that correlated with bodyguarding activity, suggesting that ant pathogens can affect not only the degree of cooperation exhibited by workers, but also the fitness of the host plants that they inhabit. Finally, we investigated the phylogenetic placement of eight new positive-sense single-stranded RNA viruses that were differentially associated with illness-related behavioral changes in bodyguard ants.

**Figure 1.**
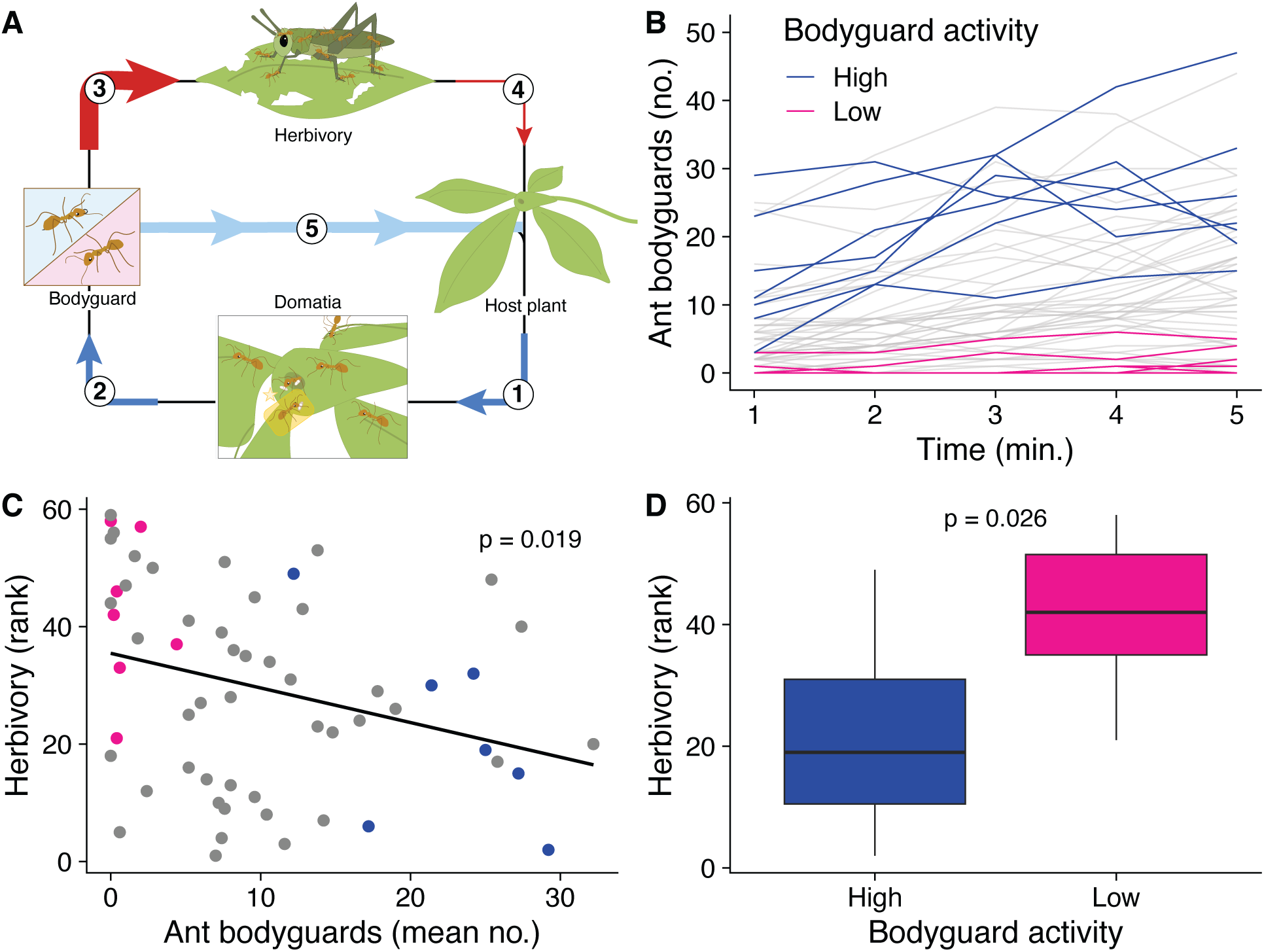
**A.** Summary of a typical ant-plant defensive mutualism. (1) Host plants produce specialized structures (domatia) that (2) attract ants that benefit from the food and shelter the plant provides. (3) As they forage for food, ant “bodyguards” defend the plant against attacking herbivores and other enemies. (4) This reduces the frequency and duration of herbivore visits to the host plant, and (5) ultimately benefits the plants by reducing tissue damage caused by herbivory. **B.** Spaghetti plot of behavioral trials across all 60 colonies. Blue and pink lines show colonies used for RNA-Seq. Grey lines indicate additional colonies that were measured but not used in transcriptomic analyses. **C.** Linear model of herbivory vs. bodyguard activity. Herbivory was measured by ranking plants based on estimated leaf area missing after three weeks in the field (here, higher numbers represent greater herbivory). **D.** Herbivory on plants occupied by the low-activity (pink) and high-activity (blue) bodyguards used for RNA-Seq.

## Results

### Ant colonies vary in their effectiveness as plant bodyguards

To characterize the bodyguarding behavior of *Allomerus octoarticulatus* colonies, we conducted behavioral bioassays on 60 naturally occurring colonies nesting in *C. nodosa* at the Los Amigos Biological Station (12°34’9” S, 70°6’0.40” W) in Madre de Dios, Peru from December 2018 to January 2019. Ant colonies varied considerably in bodyguarding behavior, from almost no ants attacking herbivores (even though all plants were occupied by *A. octoarticulatus* colonies) to sustained high recruitment to herbivores (Fig. 1, Supplementary Video 1). The ambient temperature, plant size (a proxy for ant colony size, Frederickson et al., 2012), and herbivore age (e.g., juvenile or adult), herbivore species, and herbivore size all had non-significant effects on bodyguarding behavior, but significantly fewer ants attacked herbivores in the afternoon than in the morning (Supplementary Fig. 1). The mean number of ants that attacked herbivores in bioassays significantly predicted herbivory over three weeks on the associated young leaves; plants with more active ant bodyguards experienced less herbivory (Fig. 1, linear regression, *R^2^* = 0.094, *p* = 0.019).

### Gene expression correlates with variation in ant bodyguarding behavior across colonies

We used RNA-Seq to assemble a *de novo* transcriptome for *A. octoarticulatus* (see Methods) and measured gene expression in ant bodyguards from 14 colonies, seven colonies each of high-quality bodyguards and low-quality bodyguards. For six colonies (Supplementary Table 1), we also generated RNA-Seq data from both ant bodyguards attacking herbivores on leaves and broodcare workers collected from inside the nearest domatium (i.e., ant nest).

We found 40 genes that were differentially expressed between high- and low-quality bodyguards, comprising 15 genes upregulated in low-quality bodyguards and 25 genes upregulated in high-quality bodyguards (Fig. 2). These genes have expression patterns correlated with better or worse ant defense of plants, meaning that their expression either generates or responds to changes in the mutualistic behavior of ants towards their host plants. Only one sample (s8), characterized as highly aggressive (average activity of 27.2 attacks/min), did not show the same expression pattern for most genes as the rest of the high-activity bodyguards (Fig. 2A). At the transcript level, we found 13 genes with differential transcript usage (DTU) between high- and low-quality bodyguards participating in different metabolic pathways (Supplementary Fig. 2, Supplementary Table 2). Manual annotation revealed only a few DEGs with functions studied in other insects, including cytoskeleton proteins and protease genes overexpressed in low-activity bodyguards (Fig. 2B and 2C, Supplementary Table 3). We used Gene Set Enrichment Analysis (GSEA) (FDR < 0.05) to test for gene sets that differed between high- and low-activity bodyguards, but we did not find any statistically significant gene sets (Supplementary Table 4). However, we would not necessarily expect GSEA to identify distinct expression profiles between high- and low-quality bodyguards if the behavioral differences among bodyguard ants result from other factors such as viral infections.

**Figure 2.**
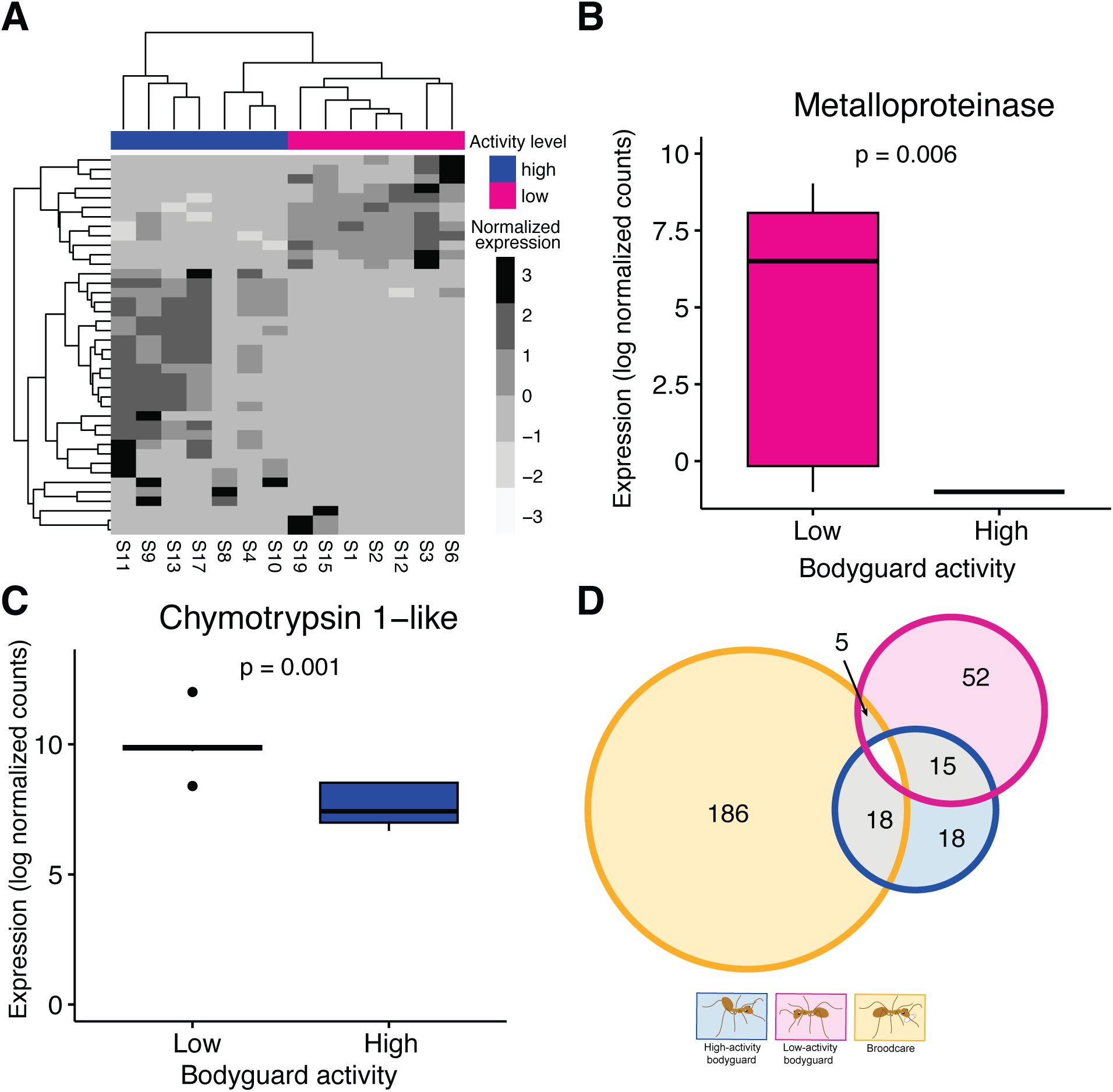
**A.** Heatmap showing 40 DEGs, with 15 genes upregulated in low-activity bodyguards and 25 genes upregulated in high-activity bodyguards. Comparison of **B.** Metalloproteinase and **C.** Chymotrypsin 1−like gene expression between low- and high-activity bodyguards. **D.** Venn diagram illustrating DEGs shared among workers doing different tasks. Most genes are uniquely highly expressed in broodcare workers.

### Viral infections are associated with mutualistic behavior

Eight of the 40 genes differentially expressed between high- and low-activity bodyguards were viral sequences, suggesting that viral infection may influence *A. octoarticulatus* behavior, or *vice versa*. These transcripts came from eight new positive-sense single-stranded RNA (+ssRNA) viruses, one matching homologous sequence from the family Picornaviridae, three matching Iflaviridae, and three others matching Dicistroviridae viruses (Supplementary Table 5). We also identified one virus in the family Narnaviridae (Supplementary Table 6).

To further examine the evolutionary history of these potential new Picorna-like and Narna viruses, we reconstructed a phylogeny comparing their RNA-dependent RNA polymerase (RdRp) sequences, a highly conserved viral domain, against RdRp regions of representative viruses within every known viral family (Fig. 3, Supplementary Tables 7 and 8). Phylogenetic analyses supported the placement of these viruses within the picorna- and narna-like clades. However, the viruses associated with *A. octoarticulatus* appear to be unique lineages that may represent new taxa. DGE analyses revealed different levels of viral expression between high- and low-activity bodyguards (Fig. 4A-H).

**Figure 3.**
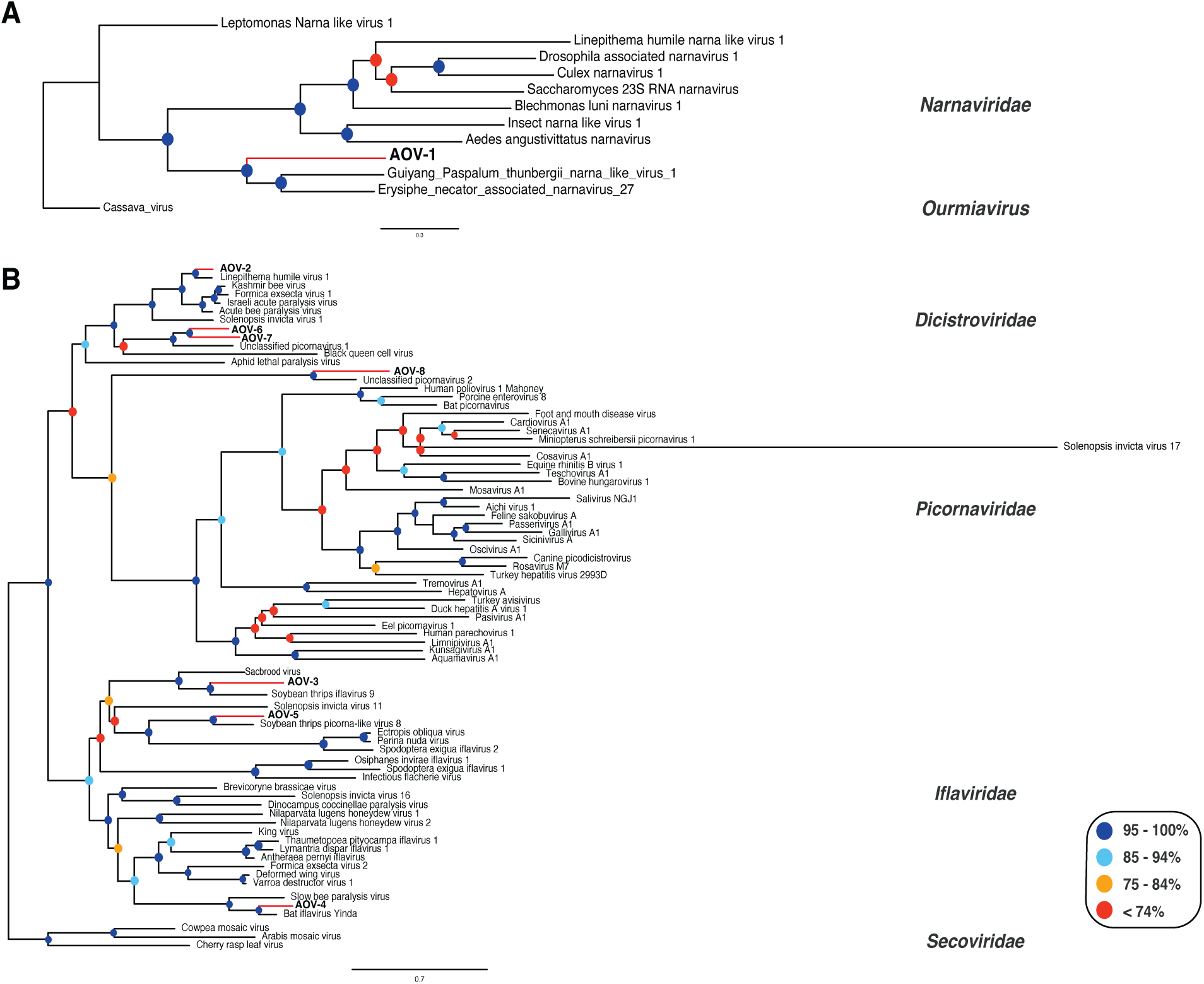
Phylogenetic reconstructions of new viruses associated with *A. octoarticulatus* based on comparisons of RNA-dependent RNA polymerase (RdRp) sequences of **A.** a narna-like virus (AOV-1, in bold) from *A. octoarticulatus* and representative species within the family Narnaviridae (order Wolframvirales), and **B.** other viruses (AOV-2 to AOV-8, in bold) from *A. octoarticulatus*, and representative species within the families Dicistroviridae, Picornaviridae, and Iflaviridae in the order Picornavirales. Three species of the family Secoviridae and an ourmiavirus are the outgroups of the trees in **B** and **A**, respectively. Ultrafast bootstrap (ufBS) support values are indicated by colored circles on respective nodes.

**Figure 4.**
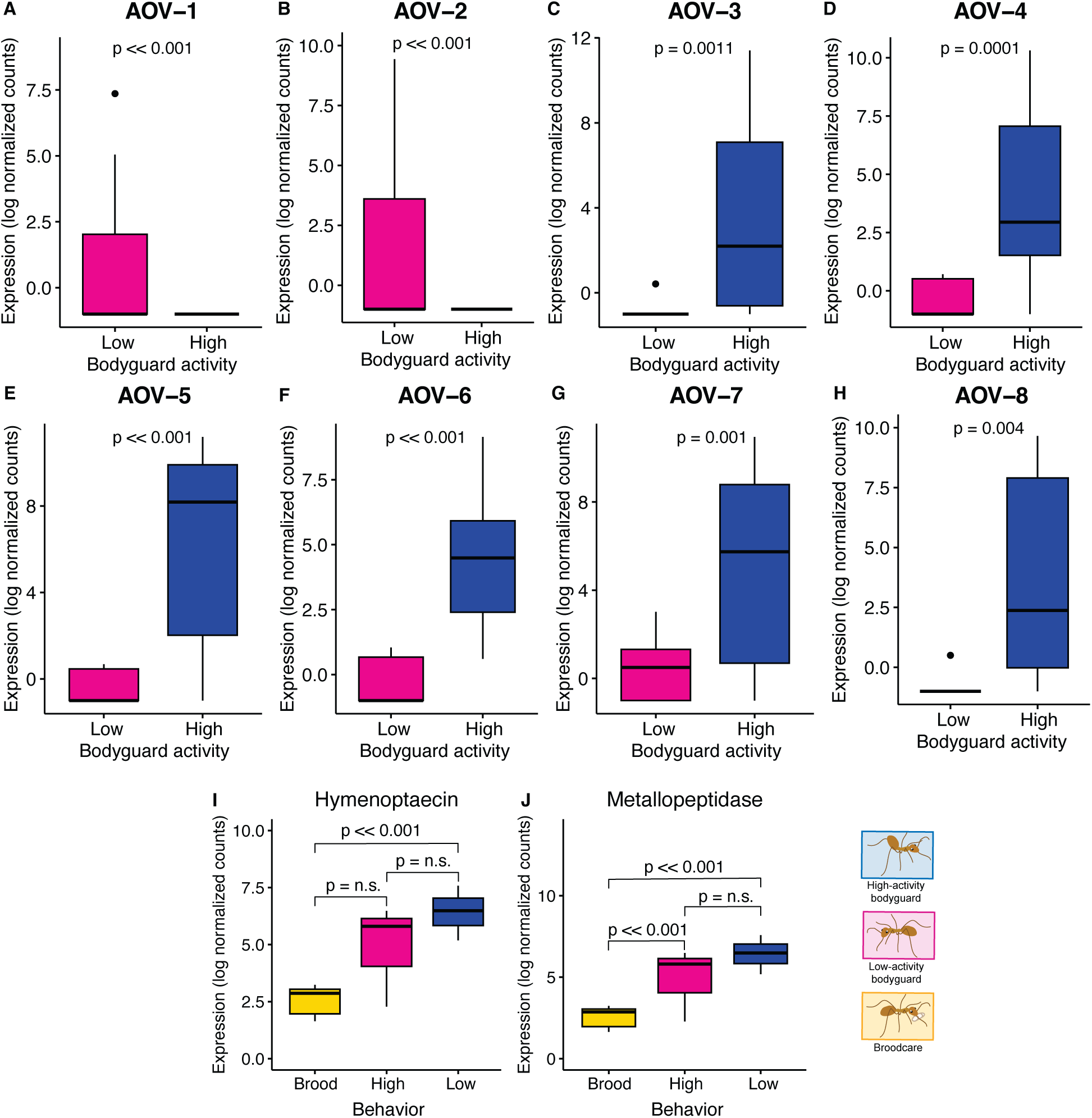
Differences in viral transcript expression between high- and low-activity bodyguards. **A**. The narnavirus: *A. octoarticulatus* virus 1 (AOV-1). **B.** A picorna-like virus highly expressed in low-activity bodyguards: *A. octoarticulatus* virus 2 (AOV-2). **C-H.** Picorna-like viruses highly expressed in high-activity bodyguards: *A. octoarticulatus* virus 3 (AOV-3), *A. octoarticulatus* virus 4 (AOV-4), *A. octoarticulatus* virus 5 (AOV-5), *A. octoarticulatus* virus 6 (AOV-6), *A. octoarticulatus* virus 7 (AOV-7), and *A. octoarticulatus* virus 8 (AOV-8). **I-J.** Expression of two immunity-related genes among broodcare workers, low-activity bodyguards, and high-activity bodyguards.

### Gene expression correlates with ant task within colonies

We found 155 DEGs between broodcare workers and high-activity bodyguards; 133 were upregulated in brood-care workers and 22 were upregulated in high-activity bodyguards (Fig. 5A, Supplementary Table 9). We observed similar patterns of gene expression when comparing broodcare workers and low-activity bodyguards, which identified 197 DEGs, 144 upregulated in broodcare workers and 53 genes upregulated in low-activity bodyguards (Fig. 5B, Supplementary Table 10). Seventy-eight genes were shared between these two comparisons (Supplementary Table 11). However, none of those genes were viral transcripts, unlike in the high-versus low-activity bodyguard comparison. At the transcript level, we obtained 28 genes with significant DTU between broodcare workers and high-activity bodyguards, and 40 DTU genes between broodcare workers and low-activity bodyguards (Supplementary Table 2). Finally, we compared the DEGs obtained in every comparison and found that most of the genes are uniquely highly expressed in broodcare workers in comparison to low- and high-activity bodyguards (Fig. 2D and Supplementary Fig. 3).

**Figure 5.**
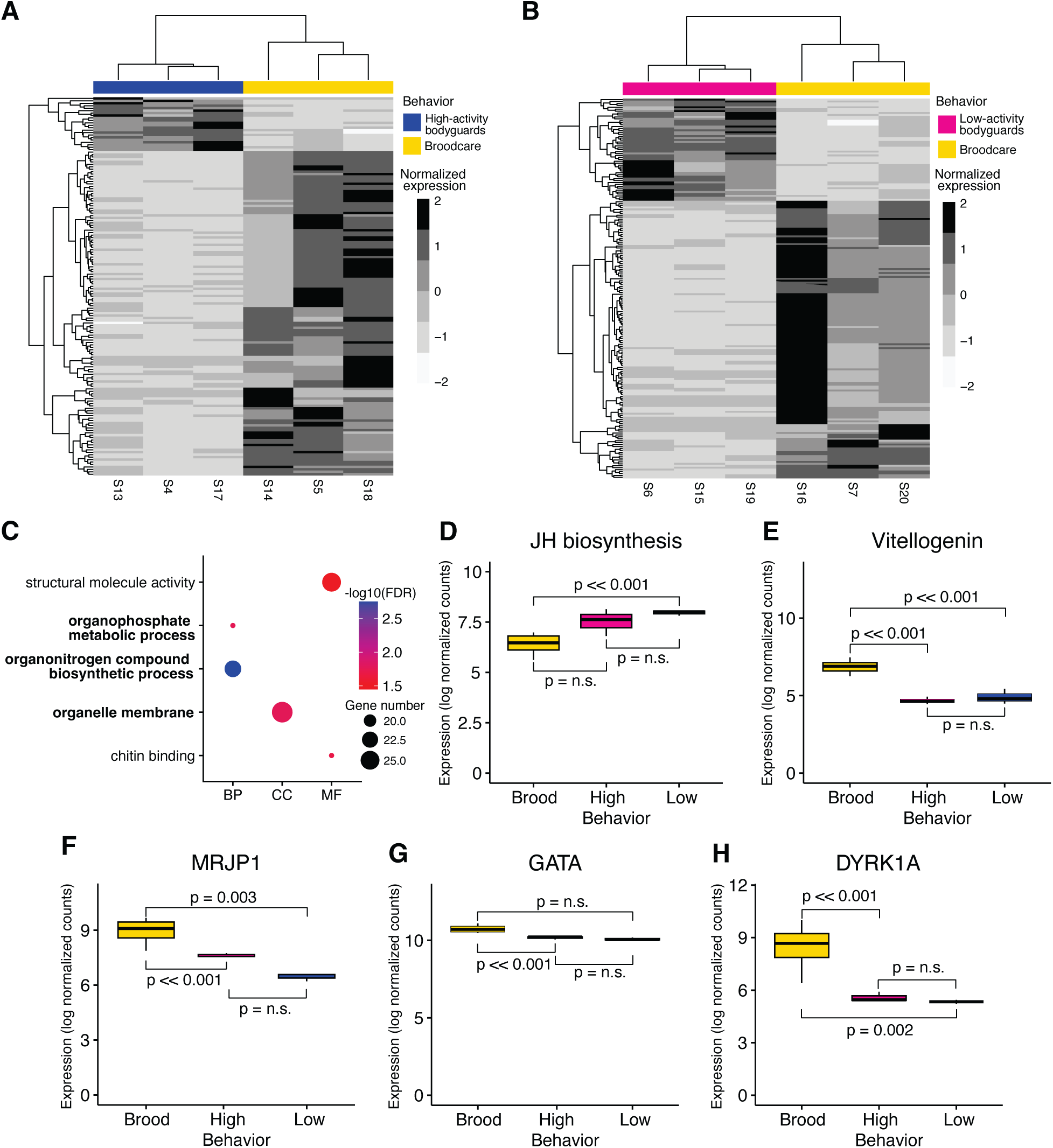
**A.** Heatmap showing 155 DEGs between broodcare workers (133 genes upregulated) and high-activity bodyguards (22 genes upregulated). **B.** Heatmap comparing 197 DEGs between broodcare workers (144 genes upregulated) and low-activity bodyguards (53 genes upregulated). **C.** Gene Set Enrichment Analysis (GSEA) results show three GO terms (in bold) overrepresented in bodyguards, and two in broodcare workers (FDR < 0.05). Ontologies: Biological Process (BP), Cellular Component (CC), and Molecular Function (MF). Gene number is number of genes in gene set. **D.** A subset of genes differentially expressed between workers performing different tasks, including juvenile hormone (JH) biosynthesis, vitellogenin, and major royal jelly protein 1-like (MRJP1). **E.** Serine-threonine protein kinase genes highly expressed in broodcare workers, including dual specificity tyrosine-phosphorylation-regulated kinase (DYRK1A) and GATA zinc finger domain-containing protein (GATA).

Again, we used GSEA (FDR < 0.05) to reveal molecular functions that differ between bodyguard and broodcare workers (Fig. 5C, and Supplementary Table 12). We found DEGs upregulated in broodcare workers significantly enriched for two gene ontology (GO) terms or gene functional categories involving chitin binding and structural molecule activity. Another three GO terms were enriched in bodyguards (Fig. 5C). We manually verified the GO annotation of every DEG (Supplementary Tables 9 and 10) and found genes associated with juvenile hormone (JH) (Fig. 5D), vitellogenin pathways (Fig. 5E), and major royal jelly protein 1-like (Fig. 5F) highly expressed in broodcare workers. Our results also revealed the overexpression of serine-threonine protein kinase genes (Fig. 5). Phylogenetic analysis of those kinase genes showed they cluster within the GATA zinc finger domain-containing protein (GATA) (Fig. 5G), and dual-specificity tyrosine-phosphorylation-regulated kinase (DYRK1A) (Fig. 5H) clades (Supplementary Fig. 3). Finally, we also identified genes known to be involved in insect immunity against viruses that were differentially expressed between broodcare workers and bodyguards (Fig. 4I-J).

## Discussion

Using field behavioral assays and RNA-Seq, we explored gene expression associated with bodyguarding behavior in *Allomerus octoarticulatus,* an obligately symbiotic ant species that defends its hosts against herbivores. We found differentially expressed genes that either generate or respond to variation in the quality of *A. octoarticulatus* colonies as plant mutualists. Many of the DEGs were viral transcripts, suggesting that viral infections explain why some *A. octoarticulatus* colonies are highly effective plant bodyguards, while others are slow to recruit to herbivores, allowing damage to accumulate on their host plants.

The high expression of viral transcripts suggests that colonies had active viral infections that modulated their bodyguarding behavior. Inactive bodyguards were infected by viruses phylogenetically related to viruses causing poor locomotion and paralysis in Hymenoptera. AOV-2, a picornavirus highly expressed in ineffective *A. octoarticulatus* bodyguards, is closely related to viruses in the acute bee paralysis (ABP) viral complex, namely acute bee paralysis virus (ABPV), Israeli acute paralysis virus (IAPV), and Kashmir bee virus (KBV). These viruses are well-known and widespread honeybee RNA viruses that induce paralysis and increase bee mortality (Amiri et al., 2019). We also identified AOV-1, a narna-like virus highly expressed in low-activity bodyguards that belongs to the same family as another ant-associated virus, *Linepithema humile* narna-like virus 1 (LhuNLV1), that has previously been described as influencing *L. humile* immunity (Lester et al., 2019). These viruses might reduce ant foraging, as previously observed in colonies of invasive fire ants infected with the dicistrovirus *Solenopsis invicta* virus 1 (SINV-1) (Hsu et al., 2018). In other words, ineffective bodyguards might be poor symbionts because they are sick.

However, we also found several viruses overexpressed in high-activity bodyguards (AOV-3, AOV-4, AOV-5, AOV-6, AOV-7, and AOV-8), and these viruses could potentially be inducing greater activity or aggression in their ant hosts to promote their own transmission. One of the best-known examples of parasite manipulation of host behavior occurs in (“zombie”) ants, in which the fungal parasite *Ophiocordyceps* manipulates the behavior of its ant hosts to promote its own onward transmission (de Bekker 2019). Similarly, Hsu et al. (2019) report aggressive behavior in the ant *Anoplolepis gracilipes* being shaped by viral disease transmission triggered by a dicistrovirus. In bees, iflaviruses such as Sacbrood virus and Deformed wing virus (DWV) increase aggressiveness in infected colonies of *Apis cerana* (Hasegawa et al., 2023) and *A. mellifera* (Fujiyuki et al., 2004; Terio et al., 2008, but see Rortais et al 2006). Thus, the viruses that were highly expressed in very active *A. octoarticulatus* bodyguards may also be manipulating the behavior of their ant hosts, with potential consequences for both their own transmission and for the degree of protection afforded by the ants. The new viruses we identified also increase the known viral diversity associated with ants (Baty et al., 2020).

Changes in aggressiveness might result from viral infections that trigger immune responses in ant workers. As part of ants’ humoral immunity, we found *hymenoptaecin* (*hym*) genes highly expressed in low-activity bodyguards. Previous studies described high expression of this infection-inducible antimicrobial peptide after ant workers are immune-challenged (Gupta et al., 2015; Yek et al., 2013; Negroni et al., 2020; Von et al., 2016). Expression of *hym* genes in ants has also been reported to be higher in foragers than in broodcare workers (Qui et al., 2017), as the former are exposed to more viruses from spending more time outside the nest where they may more regularly encounter viruses and their vectors. We also found higher expression of genes encoding matrix metalloproteinases (MMPs) in ants that are poor plant bodyguards. MMPs can be expressed in response to viral infections in Lepidoptera (Liu et al., 2022), and participate in insect innate immune regulation (Vilcinskas, 2022). Finally, *chymotrypsin-1-like*, a protease gene, was highly expressed in low-activity bodyguards. Proteolytic cascades are pathogen-inducible enzymes participating in the immediate response of the innate immune system in insects (Cerenius et al., 2010, Kanost & Jiang, 2015). These results further suggest that low-activity workers might be diseased.

Comparing between bodyguards and broodcare workers in the same colony, we observed patterns of task-specific gene expression that align with those previously reported in other social insects (Buttstedt et al., 2014; Hawkings et al., 2019; Malé et al., 2017; Das & Bekker, 2022; Qiu et al., 2022). We identified serine-threonine protein kinase (STK) genes over-expressed in broodcare workers. Alleman et al. (2019) report STK genes involved in learning and memory formation in ants. Previous research has also identified other kinase genes, especially cGMP-dependent protein kinase (PKG) genes, that mediate behavioral shifts in ants (Lucas et al., 2009; Ingram et al., 2011), including in *A. octoarticulatus* (Malé et al., 2017), although we did not find PKGs among the DEGs in this study. Overall, some genes highly expressed in bodyguards also follow previous reports of transcriptomic signatures of environmental stress in ants. For instance, we found the accumulation of digestive enzymes in bodyguards, as previously reported in foragers of other ant species exposed to temperature stress (Vatanparast et al., 2021).

Variation in gene expression has been associated with partner quality in pollination (Lüthi et al., 2022) and insect-microbe interactions (Koga et al., 2022;), but it has not yet been explored in defensive mutualisms. By combining field studies with transcriptomics, we report changes in gene expression associated with how ants cooperate with plants and find that viral infections may explain why some ants are better mutualists than others.

## Methods

### Field bioassays and sample collection

We measured bodyguard effectiveness in 60 naturally occurring colonies of *A. octoarticulatus* in the Peruvian Amazon. We collected live grasshoppers, almost entirely in the genus *Orphulella* (Supplementary Fig. 1), at our field site and attached these grasshoppers to *C. nodosa* leaves at a consistent distance to the domatium (i.e., nest) entrance using entomological pins (Supplementary Video 1). To measure ant recruitment, we counted the number of ants interacting with (i.e., stinging, biting, or walking on) the grasshopper each minute for five minutes and then averaged these counts to get a measure of bodyguarding activity (Fig. 1B, Supplementary Table 1). We also measured many factors that could potentially influence ant behavior in the bioassay, including time of day, ambient temperature, herbivore size, age, and species, and host plant size (i.e., number of domatia, a proxy for ant colony size, Frederickson et al. 2012), but found that they mostly had non-significant effects on ant behavior (Supplementary Fig. 1). Only time of day significantly predicted ant recruitment, with fewer ants recruiting to grasshoppers in the afternoon than the morning (Supplementary Fig. 1). We also measured herbivory on each plant in the field; three weeks before the bioassays, we marked up to three domatia per plant that had attached leaves with minimal herbivore damage, and then after conducting the bioassays three weeks later, we collected all the leaves from the marked domatia, grouped them by plant, and ranked the plants in order of herbivore damage, with a rank of 1 indicating the least herbivore damage and a rank of 60 indicating the most damage (Fig. 1C).

We collected bodyguards and broodcare workers from 14 colonies for transcriptomic analyses, evenly split between colonies with high- and low-activity bodyguards (Supplementary Table 1). Both high- and low-activity colonies were sampled in both the morning and the afternoon (Supplementary Fig. 1). Immediately after each bioassay, we collected 10-15 bodyguard workers on or near the grasshopper bait. We also collected up to 15 broodcare workers from six of these ant colonies by opening domatia with secateurs and extracting workers tending brood. We collected specimens in 1.5ml RNase-free tubes with 1ml of RNAlater Stabilization Solution (Thermo Fisher Scientific), transported them back to the University of Toronto (Toronto, Canada), and stored them at -80°C until RNA extractions. We obtained permits for field sampling and experiments from Peru’s Servicio Nacional Forestal y de Fauna Silvestre (SERFOR, authorization N**°** AUT-IFS-2018-061).

### RNA-Seq library construction and sequencing

We submitted samples for RNA-Seq to the Donnelly Sequencing Center at the University of Toronto (Toronto, Canada). The samples had a median RNA Integrity Number (RIN) of 8.3. The sequencing facility processed 500 ng per sample using the NEBNext Ultra II Directional RNA Library Prep Kit for Illumina, including PolyA selection using the NEBNext Poly(A) mRNA Magnetic Isolation. Fragmentation was 15 minutes at 94 °C and samples were amplified for 9 cycles. The facility ran 1 µL top stock of each purified final library on an Agilent Bioanalyzer dsDNA High Sensitivity chip. The libraries were quantified using the Quant-iT High-Sensitivity dsDNA Assay Kit and pooled at equimolar ratios after size adjustment. The final pool was run on an Agilent Bioanalyzer dsDNA High Sensitivity chip and quantified using a NEBNext Library Quant Kit for Illumina. The quantified pool was hybridized at a final concentration of 2.15 pM and sequenced with 2x50 bp reads on the Illumina NovaSeq 6000 platform using an SP flowcell, generating an average of 28.2 million read pairs per sample. Read quality scores were 36 and sequence length after trimming ranged from 30 to 51 bp (Supplementary Table 13).

### De novo *transcriptome assembly, validation, and completeness*

Because there was no reference genome or transcriptome for any *Allomerus* species before our study, we generated a *de novo* transcriptome assembly for *Allomerus octoarticulatus.* We removed low-quality sequences and adapters using Trimmomatic v0.39 (Bolger et al., 2014), and assessed the quality of the trimmed reads with FastQC v0.11.2 (Supplementary Table 13). Using the rnaSPAdes pipeline (Bushmanova et al., 2019) and CD-HIT (Li & Godzik, 2006; Huang et al., 2010), we assembled the high-quality RNA-Seq reads into a transcriptome with 67,613 unigenes and 93,122 transcripts. Then we assessed the quality of the transcriptome with rnaQUAST (Bushmanova, et al., 2019). The assembled transcripts had an average length of 1478 bp, and the N50 was 3232 bp (Supplementary Table 14). Assessed with BUSCO (Manni et al., 2021), the transcriptome had completeness scores of 95.62% (using the Hymenoptera database) and 99.34% (using the Insecta database). We deposited the *de novo* transcriptome assembly at GenBank (NCBI database) under the submission number (available upon acceptance: XXXXXXXXXX).

### Differential expression analysis

For transcript quantification, we implemented the Salmon v1.9.0 pipeline (Patro et al., 2017). We built a Salmon index based on a decoy-aware transcriptome to avoid false mapping (Srivastava et al., 2020). We used the genome of a close relative of *A. octoarticulatus, Wasmannia auropunctata* (GenBank: GCA_000956235.1) as the ‘decoy.’ Then we aligned the RNA-Seq reads to the Salmon index. In this step, we implemented the Gibbs sampling option (-- numGibbsSamples 20) and reduced autocorrelation in the Gibbs samples (thinningFactor 100). Finally, we obtained transcript-level counts for each sample with tximport (Soneson et al., 2015).

To identify DEGs, we used the DESeq2 workflow (Love et al., 2014) in v4.2.1 (R Core Team, 2022). We extracted the final list of significant genes with the *results* function and set a false discovery rate (FDR) < 0.01 and fold change (FD) > 1.5. We made heatmaps with the R library *pheatmap*.

For the phylogenetic tree of serine-threonine protein kinase genes, we obtained Formicidae orthologues of the GATA zinc finger domain-containing proteins (OrthoDB Group 32050at36668) and dual specificity tyrosine-phosphorylation-regulated kinases (OrthoDB Group 21499at36668). We also downloaded PKG orthologues for ants using OrthoDB (Group EOG8PP0WX) and aligned those genes to PKG-encoding genes characterized in *A. octoarticulatus*: *Aofor* (GenBank KX809576) and *Aopkg* (GenBank KX809574). We aligned sequences using MAFFT v7.505 (Katoh, et al., 2005) with the algorithm L-INS-I and the default scoring matrix (BLOSUM62) and gap penalty (1.53) values. We then generated a maximum likelihood tree based on amino-acid sequences with IQ-Tree v1.6.12 software (Nguyen et al., 2015) The amino acid substitution model implemented was JTT+F+G4 (Kalyaanamoorthy, et al., 2017), and we evaluated node support with 1000 bootstrap replicates (Hoang, et al., 2017).

### Differential transcript usage analysis

We implemented the Dirichlet-multinomial framework with the R library *DRIMSeq* (Nowicka & Robinson, 2016) following Love et al (2018). First, we scaled transcripts-per-million (TPM) abundance estimates per sample with the *scaledTPM* method to generate a final matrix with count-scale data. Along with that matrix, we imported to *DRIMSeq* a transcript-to-gene mappings file to create a *dmDSdata* object containing sample and count information. We filtered data to minimal expression and proportion levels of 10 and 0.1, respectively. We used the functions *dmPrecision* and *dmFit* for the precision and proportion estimations, and *dmTest* to test for DTU. Finally, we performed two-stage testing with the R library *stageR* (Van den Berge et al., 2017) by implementing screening and confirmation stages to identify the final list of genes participating in DTU. For the latter stage, we used a target overall false discovery rate (OFDR) level of 5% (alpha = 0.05).

### Functional annotation and enrichment analysis

For annotation of the *de novo* reference transcriptome, we used the blastx function in Blast v2.11.0 (Camacho et al., 2009). We used the *de novo* transcriptome as a query against a high-quality manually annotated and non-redundant protein sequence database -UniProtKB (UniProt Consortium, 2021), with an e-value cut-off of 1e-5. We then loaded this XML file into the OmicsBox program v2.1 (OmicsBox, 2019) to perform GO mapping and annotation (Gotz et al., 2018) against the Gene Ontology database implemented in the program. For GO assignment, we set the annotation cut-off to 55 and the e-value hit filter to 1.0E-6. To obtain a more complete functional annotation for the subset of DE genes obtained in the DEG and DTU analyses, we used the software package InterProScan (Jones et al., 2014) in addition to BLAST (Altschul et al., 1990) in OmicsBox. We implemented these annotation steps to allow for the sequences to be screened against more databases. We performed sequence annotation with the public EMBL-EBI InterPro web service by loading the fasta sequences and running the analysis under the default setting. We then ran a blast search against the non-redundant NCBI database with an e-value of 1.0E-5. We then used the same previously described approach for GO annotation and mapping of the DE genes. We also implemented Gene Set Enrichment Analysis (GSEA) in OmicsBox using an FDR value of 0.05. Finally, we retrieved Kyoto Encyclopedia of Genes and Genomes (KEGG) pathways (Kanehisa, 2000) by annotating and predicting orthologues with the tool EggNOG-Mapper (Huerta-Cepas et al., 2019) in OmicsBox.

### Virus characterization and phylogenetic analysis

We identified genes with GO terms corresponding to viral-related processes. The genes were manually annotated by aligning them against the NCBI non-redundant protein sequence database using blastx. We found eleven sequences of eight genes from picornavirales and Naranavirus that were homologous to the viral genes we identified. We further extracted potential open reading frames (ORF) and conserved domains by using those genes as queries on NCBI’s ORF Finder and Conserved Domain Database (CDD), respectively.

For phylogenetic reconstruction, we used the full-length amino acid sequences of the non-structural RNA-dependent RNA polymerase (RdRp), a highly conserved gene suitable for phylogenetic reconstructions of viruses (Venkataraman et al., 2018). We extracted RdRp sequences from genes identified as viral from CDD. Unclassified and classified representative amino acid sequences of species of each family were used in the phylogeny as suggested by Xavier et al. (2021). We also included viral sequences with the most similar matches based on the blastx search. We used noumavirus and three species in the family Secoviridae as outgroups for the phylogenetic placement of picorna-like viruses and narna-like viruses, respectively. We aligned viral protein sequences using MAFFT v7.505 (Katoh, et al., 2005). We generated a maximum likelihood phylogenetic tree based on amino acid sequences with the software program IQ-Tree v1.6.12 software (Nguyen et al., 2015), and the amino acid substitution model Blosum62+R5 (Henikoff and Henikoff, 1992). Finally, we evaluated the support for each node with 1000 bootstrap replicates.

## Acknowledgments

The authors thank the Centro de Investigación y Capacitación Río Los Amigos (CICRA) for their support of the field research, especially station manager Arianna Basto Eyzaguirre. MEF acknowledges funding from the Natural Sciences and Engineering Research Council of Canada (NSERC) Discovery Grant program, CR from a National Geographic Early Career Grant, a Sigma Xi Grant in Aid of Research, and a University of Toronto Centre for Global Change Science Graduate Scholarship, and MCT from a Vanier Canada Graduate Scholarship. We also thank Alberto Escudero for his help with fieldwork, Ina Anreiter for helping with RNA extractions, Kyle Turner and the Donnelly Sequencing Center for sequencing samples, Santiago Sanchez and Pierre-Jean Malé for bioinformatic support, Gianpiero Fiorentino and Patricio Picon for help with graphical design, and Naomi Pierce for feedback that improved the manuscript.

## Author contributions

MCT, CR, JH, and MEF designed and performed the research. MCT, CR, HX, and MEF analyzed the data. MCT wrote the article with input from all the authors.

## Declaration of interests

The authors declare no competing interests.

## Data availability

Lists of DEGs and DTUs are available in the supplementary information. Raw sequencing and transcriptome assembly data will be deposited in NCBI databases. Field data and code will be archived on Dryad and Zenodo, respectively, on acceptance, as well as in the GitHub repository: https://github.com/mariatocora/Transcriptomic-analysis-ant-plant.

## Notes

### Competing Interest Statement

The authors have declared no competing interest.

### Summary of Updates

Minor edits to figures and text

## References

1. A. Alleman, M. Stoldt, B. Feldmeyer, S. Foitzik, Tandem-running and scouting behaviour are characterized by up-regulation of learning and memory formation genes within the ant brain. Molecular Ecology 28, 2342–2359 (2019).

2. S. F. Altschul, W. Gish, W. Miller, E. W. Myers, D. J. Lipman, Basic local alignment search tool. Journal of Molecular Biology 215, 403–410 (1990).

3. E. Amiri, et al., Israeli Acute Paralysis Virus: Honey Bee Queen–Worker Interaction and Potential Virus Transmission Pathways. Insects 10, 9 (2019).

4. R. T. Batstone, L. T. Burghardt, K. D. Heath, Phenotypic and genomic signatures of interspecies cooperation and conflict in naturally occurring isolates of a model plant symbiont. Proc. R. Soc. B. 289, 20220477 (2022).

5. R. T. Batstone, A. M. O’Brien, T. L. Harrison, M. E. Frederickson, Experimental evolution makes microbes more cooperative with their local host genotype. Science 370, 476–478 (2020).

6. BioBam Bioinformatics, OmicsBox – Bioinformatics Made Easy, BioBam (2019).

7. A. M. Bolger, M. Lohse, B. Usadel, Trimmomatic: a flexible trimmer for Illumina sequence data. Bioinformatics 30, 2114–2120 (2014).

8. Bronstein, JL, Mutualism. (Oxford University Press, New York., 2015).

9. E. Bushmanova, D. Antipov, A. Lapidus, A. D. Prjibelski, rnaSPAdes: a de novo transcriptome assembler and its application to RNA-Seq data. GigaScience 8, giz100 (2019).

10. A. Buttstedt, R. F. A. Moritz, S. Erler, Origin and function of the major royal jelly proteins of the honeybee (*Apis mellifera*) as members of the *yellow* gene family: Honeybee’s major royals. Biol Rev 89, 255–269 (2014).

11. C. Camacho, et al., BLAST+: architecture and applications. BMC Bioinformatics 10, 421 (2009).

12. L. Cerenius, S. Kawabata, B. L. Lee, M. Nonaka, K. Söderhäll, Proteolytic cascades and their involvement in invertebrate immunity. Trends in Biochemical Sciences 35, 575–583 (2010).

13. B. Das, C. De Bekker, Time-course RNASeq of Camponotus floridanus forager and nurse ant brains indicate links between plasticity in the biological clock and behavioral division of labor. BMC Genomics 23, 57 (2022).

14. C. De Bekker, Ophiocordyceps–ant interactions as an integrative model to understand the molecular basis of parasitic behavioral manipulation. Current Opinion in Insect Science 33, 19– 24 (2019).

15. de Bekker, C, Will, I, Das, B, Adams, R.M.M, The ants (Hymenoptera: Formicidae) and their parasites: effects of parasitic manipulations and host responses on ant behavioral ecology. 28, 1– 24 (2018).

16. B. Epstein, et al., Combining GWAS and population genomic analyses to characterize coevolution in a legume-rhizobia symbiosis. Molecular Ecology 32, 3798–3811 (2023).

17. M. E. Frederickson, Rethinking Mutualism Stability: Cheaters and the Evolution of Sanctions. The Quarterly Review of Biology 88, 269–295 (2013).

18. M. E. Frederickson, et al., The Direct and Ecological Costs of an Ant-Plant Symbiosis. The American Naturalist 179, 768–778 (2012).

19. M. L. Friesen, Widespread fitness alignment in the legume–rhizobium symbiosis. New Phytologist 194, 1096–1111 (2012).

20. T. Fujiyuki, et al., Novel Insect Picorna-Like Virus Identified in the Brains of Aggressive Worker Honeybees. J Virol 78, 1093–1100 (2004).

21. S. D. Gillespie, L. S. Adler, Indirect effects on mutualisms: parasitism of bumble bees and pollination service to plants. Ecology 94, 454–464 (2013).

22. S. Gotz, et al., High-throughput functional annotation and data mining with the Blast2GO suite. Nucleic Acids Research 36, 3420–3435 (2008).

23. S. K. Gupta, et al., Scrutinizing the immune defence inventory of Camponotus floridanus applying total transcriptome sequencing. BMC Genomics 16, 540 (2015).

24. N. Hasegawa, et al., Evolutionarily diverse origins of deformed wing viruses in western honey bees. Proc. Natl. Acad. Sci. U.S.A. 120, e2301258120 (2023).

25. C. Hawkings, T. L. Calkins, P. V. Pietrantonio, C. Tamborindeguy, Caste-based differential transcriptional expression of hexamerins in response to a juvenile hormone analog in the red imported fire ant (Solenopsis invicta). PLoS ONE 14, e0216800 (2019).

26. S. Henikoff, J. G. Henikoff, Amino acid substitution matrices from protein blocks. Proc. Natl. Acad. Sci. U.S.A. 89, 10915–10919 (1992).

27. D. T. Hoang, O. Chernomor, A. Von Haeseler, B. Q. Minh, L. S. Vinh, UFBoot2: Improving the Ultrafast Bootstrap Approximation. Molecular Biology and Evolution 35, 518–522 (2018).

28. H.-W. Hsu, M.-C. Chiu, C.-C. Lee, C.-Y. Lee, C.-C. S. Yang, The Association between Virus Prevalence and Intercolonial Aggression Levels in the Yellow Crazy Ant, *Anoplolepis gracilipes* (Jerdon). Insects 10, 436 (2019).

29. H.-W. Hsu, M.-C. Chiu, D. Shoemaker, C.-C. S. Yang, Viral infections in fire ants lead to reduced foraging activity and dietary changes. Sci Rep 8, 13498 (2018).

30. Y. Huang, B. Niu, Y. Gao, L. Fu, W. Li, CD-HIT Suite: a web server for clustering and comparing biological sequences. Bioinformatics 26, 680–682 (2010).

31. J. Huerta-Cepas, et al., eggNOG 5.0: a hierarchical, functionally and phylogenetically annotated orthology resource based on 5090 organisms and 2502 viruses. Nucleic Acids Research 47, D309–D314 (2019).

32. K. K. Ingram, L. Kleeman, S. Peteru, Differential regulation of the foraging gene associated with task behaviors in harvester ants. BMC Ecol 11, 19 (2011).

33. James W. Baty, Mariana Bulgarella, Jana Dobelmann, Antoine Felden, Philip J. Lester, Viruses and their effects in ants (Hymenoptera: Formicidae) *Myrmecol News* 30, 213–228 (2020).

34. B. M. Jones, et al., Individual differences in honey bee behavior enabled by plasticity in brain gene regulatory networks. eLife 9, e62850 (2020).

35. P. Jones, et al., InterProScan 5: genome-scale protein function classification. Bioinformatics 30, 1236–1240 (2014).

36. S. Kalyaanamoorthy, B. Q. Minh, T. K. F. Wong, A. Von Haeseler, L. S. Jermiin, ModelFinder: fast model selection for accurate phylogenetic estimates. Nat Methods 14, 587–589 (2017).

37. M. Kanehisa, KEGG: Kyoto Encyclopedia of Genes and Genomes. Nucleic Acids Research 28, 27–30 (2000).

38. M. R. Kanost, H. Jiang, Clip-domain serine proteases as immune factors in insect hemolymph. Current Opinion in Insect Science 11, 47–55 (2015).

39. K. Katoh, MAFFT version 5: improvement in accuracy of multiple sequence alignment. Nucleic Acids Research 33, 511–518 (2005).

40. E. T. Kiers, M. G. Hutton, R. F. Denison, Human selection and the relaxation of legume defences against ineffective rhizobia. Proc. R. Soc. B. 274, 3119–3126 (2007).

41. R. Koga, et al., Single mutation makes Escherichia coli an insect mutualist. Nat Microbiol 7, 1141–1150 (2022).

42. P. Kohlmeier, A. R. Alleman, R. Libbrecht, S. Foitzik, B. Feldmeyer, Gene expression is more strongly associated with behavioural specialization than with age or fertility in ant workers. Molecular Ecology 28, 658–670 (2019).

43. P. J. Lester, K. H. Buick, J. W. Baty, A. Felden, J. Haywood, Different bacterial and viral pathogens trigger distinct immune responses in a globally invasive ant. Sci Rep 9, 5780 (2019).

44. W. Li, A. Godzik, Cd-hit: a fast program for clustering and comparing large sets of protein or nucleotide sequences. Bioinformatics 22, 1658–1659 (2006).

45. T. Liu, et al., The dual roles of three MMPs and TIMP in innate immunity and metamorphosis in the silkworm, *Bombyx mori*. The FEBS Journal 289, 2828–2846 (2022).

46. M. I. Love, W. Huber, S. Anders, Moderated estimation of fold change and dispersion for RNA-seq data with DESeq2. Genome Biol 15, 550 (2014).

47. M. I. Love, C. Soneson, R. Patro, Swimming downstream: statistical analysis of differential transcript usage following Salmon quantification. F1000Res 7, 952 (2018).

48. C. Lucas, M. B. Sokolowski, Molecular basis for changes in behavioral state in ant social behaviors. Proc. Natl. Acad. Sci. U.S.A. 106, 6351–6356 (2009).

49. M. N. Lüthi, A. E. Berardi, T. Mandel, L. B. Freitas, C. Kuhlemeier, Single gene mutation in a plant MYB transcription factor causes a major shift in pollinator preference. Current Biology 32, 5295–5308.e5 (2022).

50. P.-J. G. Malé, et al., An ant–plant mutualism through the lens of cGMP-dependent kinase genes. Proc. R. Soc. B. 284, 20170896 (2017).

51. F. Manfredini, et al., Sociogenomics of Cooperation and Conflict during Colony Founding in the Fire Ant Solenopsis invicta. PLoS Genet 9, e1003633 (2013).

52. M. Manni, M. R. Berkeley, M. Seppey, E. M. Zdobnov, BUSCO: Assessing Genomic Data Quality and Beyond. Current Protocols 1, e323 (2021).

53. V. E. Mayer, M. E. Frederickson, D. McKey, R. Blatrix, Current issues in the evolutionary ecology of ant–plant symbioses. New Phytologist 202, 749–764 (2014).

54. A. T. Müller, et al., Combined –omics framework reveals how ant symbionts benefit the Neotropical ant-plant Tococa quadrialata at different levels. iScience 25, 105261 (2022).

55. M. A. Negroni, F. H. I. D. Segers, F. Vogelweith, S. Foitzik, Immune challenge reduces gut microbial diversity and triggers fertility-dependent gene expression changes in a social insect. BMC Genomics 21, 816 (2020).

56. L.-T. Nguyen, H. A. Schmidt, A. Von Haeseler, B. Q. Minh, IQ-TREE: A Fast and Effective Stochastic Algorithm for Estimating Maximum-Likelihood Phylogenies. Molecular Biology and Evolution 32, 268–274 (2015).

57. M. Nowicka, M. D. Robinson, DRIMSeq: a Dirichlet-multinomial framework for multivariate count outcomes in genomics. F1000Res 5, 1356 (2016).

58. R. Patro, G. Duggal, M. I. Love, R. A. Irizarry, C. Kingsford, Salmon provides fast and bias-aware quantification of transcript expression. Nat Methods 14, 417–419 (2017).

59. B. Qiu, et al., Canalized gene expression during development mediates caste differentiation in ants. Nat Ecol Evol 6, 1753–1765 (2022).

60. R Core Team, R: A Language and Environment for Statistical Computing (2017).

61. C. Riehl, M. E. Frederickson, Cheating and punishment in cooperative animal societies. Phil. Trans. R. Soc. B 371, 20150090 (2016).

62. A. Rortais, et al., Deformed wing virus is not related to honey bees’ aggressiveness. Virol J 3, 61 (2006).

63. B. E. R. Rubin, C. S. Moreau, Comparative genomics reveals convergent rates of evolution in ant–plant mutualisms. Nat Commun 7, 12679 (2016).

64. J. Sachs, E. Simms, Pathways to mutualism breakdown. Trends in Ecology & Evolution 21, 585– 592 (2006).

65. C. Soneson, M. I. Love, M. D. Robinson, Differential analyses for RNA-seq: transcript-level estimates improve gene-level inferences. F1000Res 4, 1521 (2015).

66. A. Srivastava, et al., Alignment and mapping methodology influence transcript abundance estimation. Genome Biol 21, 239 (2020).

67. V. Terio, et al., Detection of a honeybee iflavirus with intermediate characteristics between kakugo virus and deformed wing virus. New Microbiol 31, 439–444 (2008).

68. The UniProt Consortium, et al., UniProt: the universal protein knowledgebase in 2021. Nucleic Acids Research 49, D480–D489 (2021).

69. K. Van Den Berge, C. Soneson, M. D. Robinson, L. Clement, stageR: a general stage-wise method for controlling the gene-level false discovery rate in differential expression and differential transcript usage. Genome Biol 18, 151 (2017).

70. M. Vatanparast, Y. Park, Comparative RNA-Seq Analyses of Solenopsis japonica (Hymenoptera: Formicidae) Reveal Gene in Response to Cold Stress. Genes 12, 1610 (2021).

71. S. Venkataraman, B. Prasad, R. Selvarajan, RNA Dependent RNA Polymerases: Insights from Structure, Function and Evolution. Viruses 10, 76 (2018).

72. A. Vilcinskas, Matrix metalloproteinases and their inhibitors – pleiotropic functions in insect immunity and metamorphosis. The FEBS Journal 289, 2805–2808 (2022).

73. K. Von Wyschetzki, H. Lowack, J. Heinze, Transcriptomic response to injury sheds light on the physiological costs of reproduction in ant queens. Mol Ecol 25, 1972–1985 (2016).

74. M. G. Weber, A. A. Agrawal, Defense mutualisms enhance plant diversification. Proc. Natl. Acad. Sci. U.S.A. 111, 16442–16447 (2014).

75. M. G. Weber, K. H. Keeler, The phylogenetic distribution of extrafloral nectaries in plants. Annals of Botany 111, 1251–1261 (2013).

76. C. A. D. Xavier, M. L. Allen, A. E. Whitfield, Ever-increasing viral diversity associated with the red imported fire ant Solenopsis invicta (Formicidae: Hymenoptera). Virol J 18, 5 (2021).

77. S. H. Yek, J. J. Boomsma, M. Schiøtt, Differential gene expression in *Acromyrmex* leaf-cutting ants after challenges with two fungal pathogens. Mol Ecol 22, 2173–2187 (2013)

