## Supplemental Information for "In sickness and in health: RNA-Seq finds viruses associated with mutualist quality in the Amazonian plant-ant *Allomerus octoarticulatus*"

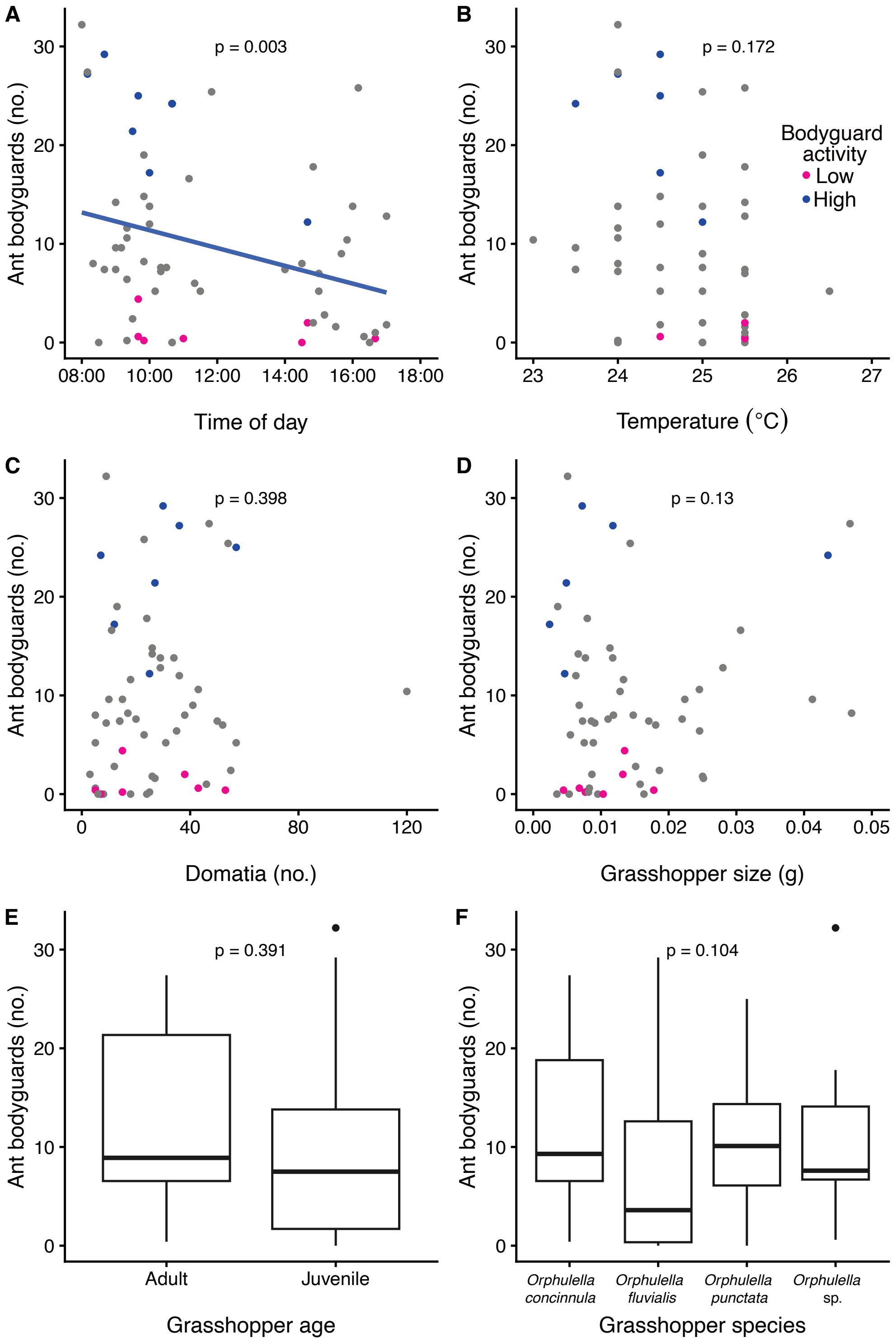


**Fig. S1.** Effects of environmental variables on bodyguard activity. **A.** Time of day. **B.** Temperature. **C.** Number of domatia on the host plant. **D.** Grasshopper size, **E.** grasshopper age, and **F.** grasshopper species used in the bioassays.

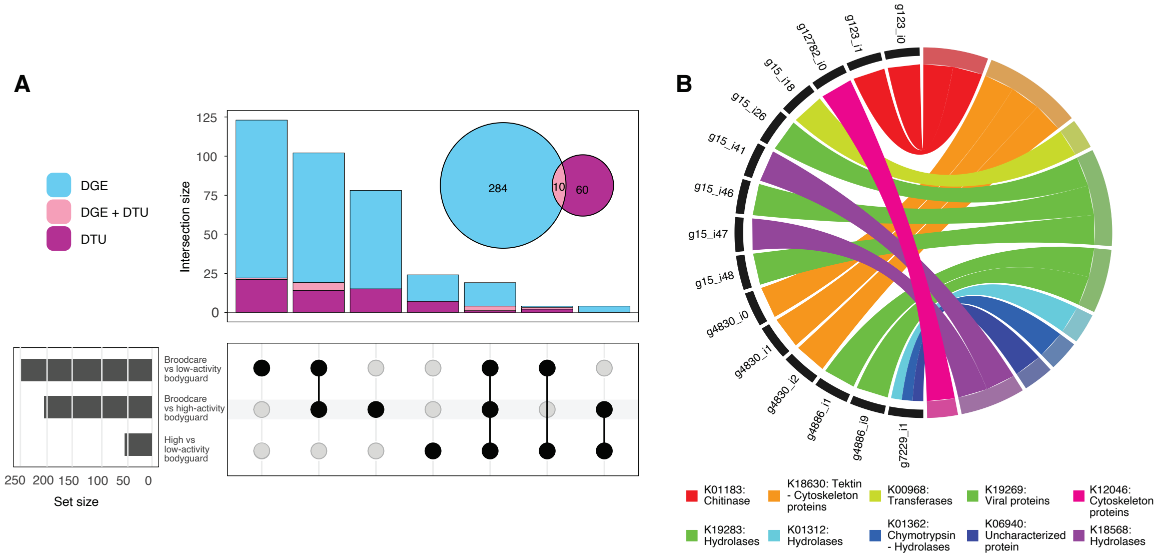


**Fig. S2. A.** UpSet plot of differentially expressed genes (DEGs) and genes with differential transcript usage (DTU) across comparisons. **B.** Chord plot of the top 15 Kyoto Encyclopedia of Genes and Genomes (KEGG) pathways overrepresented in some of the genes identified within the bodyguards' DTU comparison.

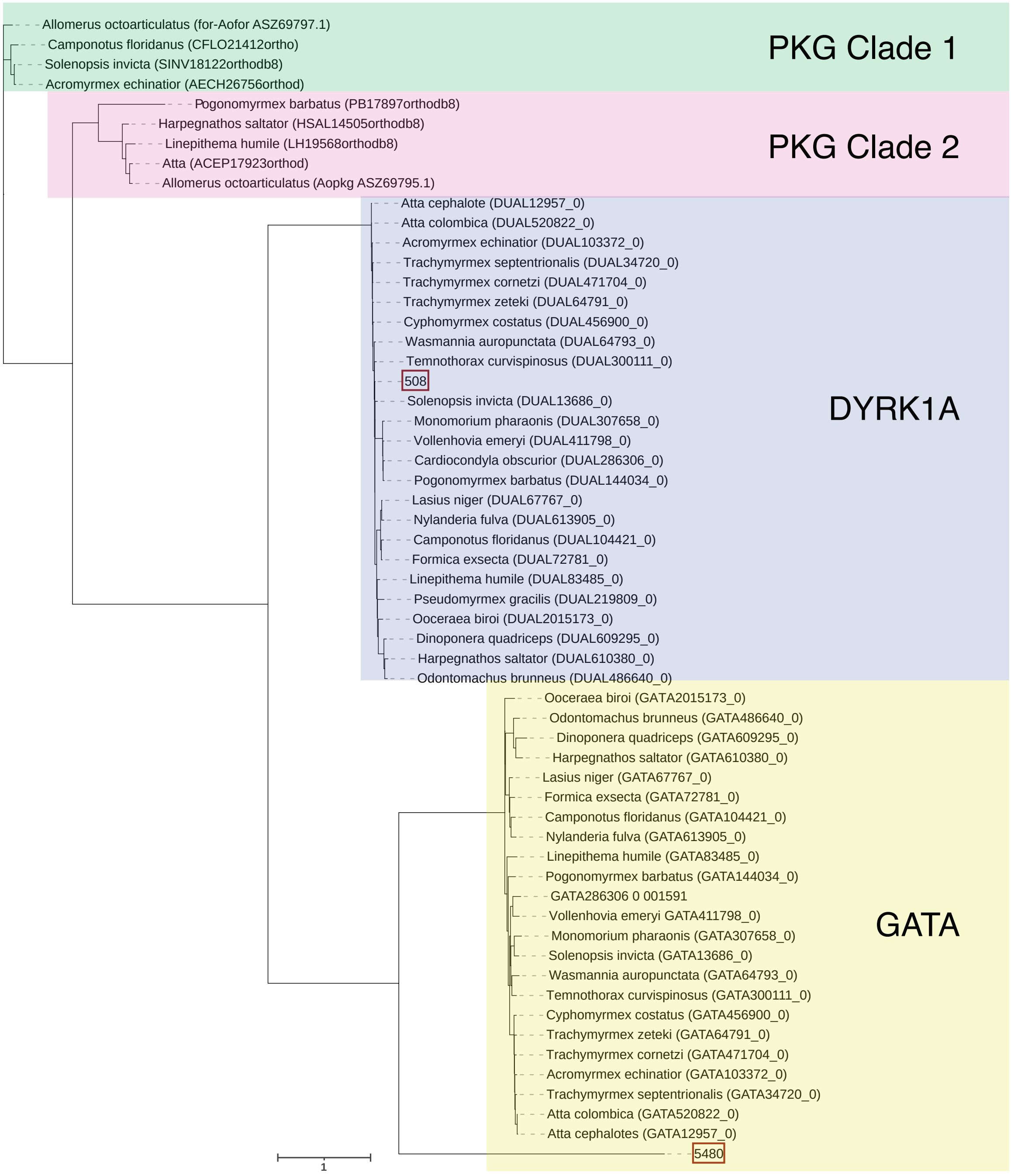


**Fig. S3.** Phylogenetic reconstruction using Maximum Likelihood (ML) best-tree and ultrafast bootstrap searches in IQ-TREE v1.6.12 of serine-threonine protein kinase genes in *A. octoarticulatus*. Genes *Aofor* (*A. octoarticulatus* orthologue of *for*) and *Aopkg* (a cGMP-dependent protein kinase, or PKG) in the *A. octoarticulatus* genome are reported in Malé et al. (2017). Genes 508 and 5480 are reported in this study. Putative orthologues for Formicidae of PKG (OrthoDB Group EOG8PP0WX), dual-specificity tyrosine-phosphorylation-regulated kinase (DYRK1A, OrthoDB Group 21499at36668), and GATA zinc finger domain-containing protein (GATA, OrthoDB Group 32050at36668) were used as outgroups.

**Table S1.** Summary of colonies and samples used for RNA-Seq.

| Colony | Bodyguarding (attackers/min) | Bodyguarding category | Sequenced bodyguards | Sequenced broodcare workers |
|---|---|---|---|---|
| 103 | 0.0 | Low | X | |
| 199 | 0.2 | Low | X | X |
| 28 | 0.4 | Low | X | |
| 207 | 0.4 | Low | X | X |
| 187 | 0.6 | Low | X | |
| 18 | 2.0 | Low | X | |
| 109 | 4.4 | Low | X | X |
| 107 | 12.2 | High | X | X |
| 145 | 17.2 | High | X | |
| 136 | 21.4 | High | X | |
| 201 | 24.2 | High | X | X |
| 193 | 25.0 | High | X | X |
| 125 | 27.2 | High | X | |
| 127 | 29.2 | High | X | |

**Table S2.** Differential transcript usage (DTU) analysis.

| | FDR adjusted p-value | Comparison |
|---|---|---|
| **g15** | 0.00404479904227522 | High- vs. low-activity bodyguards |
| **g3338** | 1.3144741514288E-07 | High- vs. low-activity bodyguards |
| **g4350** | 0.0420323815683882 | High- vs. low-activity bodyguards |
| **g4830** | 9.13292992992384E-05 | High- vs. low-activity bodyguards |
| **g4886** | 0.0257263776655302 | High- vs. low-activity bodyguards |
| **g5024** | 0.0420323815683882 | High- vs. low-activity bodyguards |
| **g5260** | 0.00159593749898846 | High- vs. low-activity bodyguards |
| **g5456** | 1.41264464927761E-15 | High- vs. low-activity bodyguards |
| **g5511** | 1.53032494831718E-05 | High- vs. low-activity bodyguards |
| **g5823** | 0.0417092460771174 | High- vs. low-activity bodyguards |
| **g7229** | 0.0376953607955536 | High- vs. low-activity bodyguards |
| **g8013** | 0.00158939564255058 | High- vs. low-activity bodyguards |
| **g8428** | 0.0257263776655302 | High- vs. low-activity bodyguards |
| **g20** | 0.000143753513213661 | Broodcare vs. high-activity bodyguards |
| **g60** | 0.0155144335000123 | Broodcare vs. high-activity bodyguards |
| **g245** | 0.0102312050442697 | Broodcare vs. high-activity bodyguards |
| **g869** | 0.0188461881991599 | Broodcare vs. high-activity bodyguards |
| **g1110** | 0.0107801812225088 | Broodcare vs. high-activity bodyguards |
| **g1150** | 0.00887124075418714 | Broodcare vs. high-activity bodyguards |
| **g1253** | 0.000277569840539996 | Broodcare vs. high-activity bodyguards |
| **g1435** | 8.736774017248E-06 | Broodcare vs. high-activity bodyguards |
| **g1669** | 0.0451067899349283 | Broodcare vs. high-activity bodyguards |
| **g1692** | 0.0360669895934468 | Broodcare vs. high-activity bodyguards |

| | | |
|---|---|---|
| **g1900** | 0.0185237334145286 | Broodcare vs. high-activity bodyguards |
| **g2176** | 0.000562596851561618 | Broodcare vs. high-activity bodyguards |
| **g2348** | 4.03101653825291E-07 | Broodcare vs. high-activity bodyguards |
| **g2640** | 0.0112376048340013 | Broodcare vs. high-activity bodyguards |
| **g3043** | 0.00179852432199161 | Broodcare vs. high-activity bodyguards |
| **g3204** | 0.00304000334997474 | Broodcare vs. high-activity bodyguards |
| **g3574** | 0.00132460859540181 | Broodcare vs. high-activity bodyguards |
| **g3623** | 0.0360669895934468 | Broodcare vs. high-activity bodyguards |
| **g5823** | 2.70963967682295E-21 | Broodcare vs. high-activity bodyguards |
| **g6067** | 3.82375013622042E-06 | Broodcare vs. high-activity bodyguards |
| **g8120** | 1.78439500060368E-06 | Broodcare vs. high-activity bodyguards |
| **g8364** | 0.0184801123132133 | Broodcare vs. high-activity bodyguards |
| **g8464** | 0.0161001488627781 | Broodcare vs. high-activity bodyguards |
| **g8495** | 0.00311734933303589 | Broodcare vs. high-activity bodyguards |
| **g8727** | 0.00473866525864315 | Broodcare vs. high-activity bodyguards |
| **g12754** | 0.00412130289122431 | Broodcare vs. high-activity bodyguards |
| **g20031** | 0.00887124075418714 | Broodcare vs. high-activity bodyguards |
| **g35676** | 0.00262336911741959 | Broodcare vs. high-activity bodyguards |
| **g12** | 3.27423316420866E-07 | Broodcare vs. low-activity bodyguards |
| **g20** | 0.0023817042605916 | Broodcare vs. low-activity bodyguards |
| **g226** | 0.030886106570575 | Broodcare vs. low-activity bodyguards |
| **g368** | 0.030886106570575 | Broodcare vs. low-activity bodyguards |
| **g508** | 0.0367757174480848 | Broodcare vs. low-activity bodyguards |
| **g822** | 5.45194492893232E-11 | Broodcare vs. low-activity bodyguards |
| **g960** | 0.0445906117101265 | Broodcare vs. low-activity bodyguards |
| **g1110** | 0.0417916184340779 | Broodcare vs. low-activity bodyguards |

| | | |
|---|---|---|
| **g1150** | 0.0301174189531593 | Broodcare vs. low-activity bodyguards |
| **g1211** | 0.0187828836696935 | Broodcare vs. low-activity bodyguards |
| **g1253** | 0.024714478429615 | Broodcare vs. low-activity bodyguards |
| **g1306** | 0.00120272870862363 | Broodcare vs. low-activity bodyguards |
| **g1388** | 0.0023817042605916 | Broodcare vs. low-activity bodyguards |
| **g1672** | 0.00120272870862363 | Broodcare vs. low-activity bodyguards |
| **g2002** | 0.0417916184340779 | Broodcare vs. low-activity bodyguards |
| **g2039** | 0.00052271922046481 | Broodcare vs. low-activity bodyguards |
| **g2157** | 0.0334247410498996 | Broodcare vs. low-activity bodyguards |
| **g2236** | 0.0334247410498996 | Broodcare vs. low-activity bodyguards |
| **g2274** | 0.000311335244419448 | Broodcare vs. low-activity bodyguards |
| **g2477** | 0.0127373980443393 | Broodcare vs. low-activity bodyguards |
| **g2658** | 0.0445906117101265 | Broodcare vs. low-activity bodyguards |
| **g2793** | 0.0023817042605916 | Broodcare vs. low-activity bodyguards |
| **g2940** | 0.0120642156753268 | Broodcare vs. low-activity bodyguards |
| **g3043** | 1.42447792968756E-07 | Broodcare vs. low-activity bodyguards |
| **g3049** | 0.00206351640464708 | Broodcare vs. low-activity bodyguards |
| **g3338** | 7.99215549413164E-05 | Broodcare vs. low-activity bodyguards |
| **g3589** | 0.00197948268536575 | Broodcare vs. low-activity bodyguards |
| **g4043** | 0.0023817042605916 | Broodcare vs. low-activity bodyguards |
| **g5024** | 0.0334247410498996 | Broodcare vs. low-activity bodyguards |
| **g5511** | 1.19657102786904E-14 | Broodcare vs. low-activity bodyguards |
| **g5823** | 1.53676132611733E-20 | Broodcare vs. low-activity bodyguards |
| **g6180** | 1.55420339951527E-05 | Broodcare vs. low-activity bodyguards |
| **g6497** | 0.0215219986222353 | Broodcare vs. low-activity bodyguards |
| **g6663** | 0.0229440431605382 | Broodcare vs. low-activity bodyguards |

| | | |
|---|---|---|
| **g7229** | 0.0370677243333985 | Broodcare vs. low-activity bodyguards |
| **g9002** | 0.00186431814797682 | Broodcare vs. low-activity bodyguards |
| **g10445** | 0.0266948820734706 | Broodcare vs. low-activity bodyguards |
| **g10518** | 2.88853602567832E-05 | Broodcare vs. low-activity bodyguards |
| **g13462** | 0.0334247410498996 | Broodcare vs. low-activity bodyguards |
| **g17778** | 0.00935002392587385 | Broodcare vs. low-activity bodyguards |

**Table S3.** Analysis and annotation of genes differentially expressed between high- and low-quality ant bodyguards.

| | baseMean | log2FoldChange | lfcSE | stat | pvalue | padj | Annotation (Blast and InterProScan) |
|---|---|---|---|---|---|---|---|
| **g9305** | 321.205 | 3.465 | 0.352 | 9.851 | 6.79E-23 | 1.19E-18 | Cuticle protein 64-like |
| **g6591** | 2783.654 | 5.303 | 0.558 | 9.505 | 1.99E-21 | 1.75E-17 | Cuticle protein 64-like isoform X2 |
| **g13731** | 574.859 | 4.934 | 0.591 | 8.355 | 6.55E-17 | 3.84E-13 | Tetra-peptide repeat homeobox protein 1 |
| **g6982** | 56.626 | -24.601 | 2.987 | -8.236 | 1.79E-16 | 7.85E-13 | Chain C, Structural polyprotein, VP2 [Israeli acute paralysis virus] |
| **g13589** | 1047.438 | 5.300 | 0.673 | 7.870 | 3.54E-15 | 1.25E-11 | Chondroitin sulfate glucuronyltransferase |
| **g6663** | 89.062 | 2.353 | 0.307 | 7.657 | 1.91E-14 | 5.59E-11 | Spore wall protein 1 isoform X1 |
| **g9141** | 14.031 | -22.768 | 2.989 | -7.618 | 2.57E-14 | 6.45E-11 | RNA-dependent RNA polymerase |
| **g7216** | 2675.897 | 3.138 | 0.485 | 6.471 | 9.73E-11 | 2.14E-07 | Cuticle protein 67-like |
| **g5151** | 7332.599 | 4.535 | 0.707 | 6.415 | 1.41E-10 | 2.76E-07 | Spidroin-1-like isoform X2 |
| **g9792** | 282.087 | 1.960 | 0.316 | 6.199 | 5.67E-10 | 9.96E-07 | Z512B protein |
| **g1894** | 383.029 | 10.711 | 1.750 | 6.122 | 9.24E-10 | 1.48E-06 | Polyprotein [Soybean thrips picorna-like virus 8] |
| **g7219** | 38.359 | 4.265 | 0.718 | 5.942 | 2.82E-09 | 4.12E-06 | Alpha/beta-gliadin A-IV-like |
| **g123** | 79.374 | 1.353 | 0.237 | 5.714 | 1.10E-08 | 1.49E-05 | CHIT1 protein |
| **g13743** | 128.322 | 3.247 | 0.579 | 5.609 | 2.03E-08 | 2.55E-05 | A-kinase anchor protein 14-like |
| **g2855** | 53.752 | 7.846 | 1.458 | 5.380 | 7.43E-08 | 8.65E-05 | RNA dependent RNA polymerase [Picornaviridae sp.] |
| **g20498** | 24.103 | 1.746 | 0.325 | 5.370 | 7.88E-08 | 8.65E-05 | structural polyprotein [Dicistroviridae sp.] |
| **g5994** | 178.421 | 9.583 | 1.823 | 5.257 | 1.46E-07 | 0.000151077 | polyprotein [Bat iflavirus] |
| **g12874** | 5110.841 | -0.919 | 0.177 | -5.195 | 2.04E-07 | 0.000199383 | Probable salivary secreted peptide |
| **g5511** | 159.380 | 2.305 | 0.446 | 5.164 | 2.42E-07 | 0.000224145 | Late embryogenesis abundant protein, group 3-like |
| **g8173** | 28.280 | 1.394 | 0.271 | 5.153 | 2.56E-07 | 0.000224773 | Putative mediator of RNA polymerase II transcription subunit 12 isoform X1 |
| **g5389** | 225.193 | 1.713 | 0.342 | 5.015 | 5.31E-07 | 0.000444691 | Mediator of RNA polymerase II transcription subunit 15-like |

| | | | | | | | |
|---|---|---|---|---|---|---|---|
| **g22293** | 11.976 | 3.133 | 0.645 | 4.858 | 1.18E-06 | 0.000945349 | uncharacterized LOC105451335 (LOC105451335), mRNA |
| **g7229** | 771.068 | -2.481 | 0.514 | -4.830 | 1.36E-06 | 0.00104256 | Chymotrypsin-1-like (LOC105448752), mRNA |
| **g3684** | 248.476 | 8.105 | 1.688 | 4.802 | 1.57E-06 | 0.001144054 | RNA dependent RNA polymerase [Picornaviridae sp.] |
| **g2275** | 236.501 | 11.799 | 2.461 | 4.795 | 1.63E-06 | 0.001144054 | Polyprotein [Iflaviridae sp.] |
| **g8308** | 41.005 | -1.967 | 0.411 | -4.779 | 1.76E-06 | 0.001147026 | Protein G12 |
| **g3245** | 127.371 | 0.698 | 0.149 | 4.686 | 2.79E-06 | 0.001749387 | Protein FAM136A-like (LOC105460419), partial mRNA |
| **g24155** | 9.941 | -2.541 | 0.556 | -4.574 | 4.78E-06 | 0.002895148 | Pheromone-binding protein Gp-9-like (LOC105449955), mRNA |
| **g16177** | 19.090 | -1.344 | 0.295 | -4.560 | 5.11E-06 | 0.002959751 | Histone H1-III-like (LOC105453141), mRNA |
| **g6884** | 16.358 | -2.949 | 0.647 | -4.556 | 5.22E-06 | 0.002959751 | Uncharacterized protein LOC105203903 |
| **g12782** | 361.702 | -0.792 | 0.177 | -4.465 | 8.00E-06 | 0.004259031 | Programmed cell death protein 2-like (LOC105448647), mRNA |
| **g8064** | 82.518 | 1.164 | 0.262 | 4.449 | 8.64E-06 | 0.004339047 | uncharacterized LOC105456783 (LOC105456783), mRNA |
| **g17038** | 734.581 | -2.886 | 0.652 | -4.426 | 9.59E-06 | 0.004590805 | Protein G12 |
| **g6082** | 43.325 | 1.189 | 0.269 | 4.425 | 9.66E-06 | 0.004590805 | Cell wall-associated hydrolase |
| **g3814** | 112.429 | 9.755 | 2.214 | 4.406 | 1.05E-05 | 0.004863869 | RNA dependent RNA polymerase [Picornaviridae sp.] |
| **g6602** | 10704.514 | -0.850 | 0.194 | -4.380 | 1.19E-05 | 0.005226643 | Probable salivary secreted peptide |
| **g11332** | 698.214 | -0.586 | 0.134 | -4.374 | 1.22E-05 | 0.005240447 | Lipase 3 |
| **g6124** | 1401.779 | -0.680 | 0.157 | -4.333 | 1.47E-05 | 0.006025702 | 15-hydroxyprostaglandin dehydrogenase [NAD(+)]-like |
| **g20928** | 46.856 | 7.340 | 1.697 | 4.325 | 1.53E-05 | 0.006102568 | Structural polyprotein |
| **g16223** | 82.331 | -9.924 | 2.300 | -4.314 | 1.60E-05 | 0.006252276 | Astacin-like (LOC105453010), mRNA |

**Table S4.** Gene Set Enrichment Analysis (GSEA) comparing high- and low-activity ant bodyguards.

| GO-ID | GO Name | GO Type | Size | ES | NES | Nominal p-val | FDR(q-val) | FWER p-val |
|---|---|---|---|---|---|---|---|---|
| **GO:0007017** | microtubule-based process | BP | 1 | -1.000 | -1.323 | 0.000 | 0.463 | 0.746 |
| **GO:0099512** | supramolecular fiber | CC | 1 | -1.000 | -1.320 | 0.000 | 0.363 | 0.758 |
| **GO:0005856** | cytoskeleton | CC | 1 | -1.000 | -1.320 | 0.000 | 0.295 | 0.767 |
| **GO:0099080** | supramolecular complex | CC | 1 | -1.000 | -1.316 | 0.000 | 0.266 | 0.797 |
| **GO:0140657** | ATP-dependent activity | MF | 1 | 0.974 | 1.287 | 0.051 | 0.398 | 0.710 |
| **GO:0004386** | helicase activity | MF | 1 | 0.974 | 1.287 | 0.051 | 0.304 | 0.719 |
| **GO:0006725** | cellular aromatic compound metabolic process | BP | 5 | 0.971 | 1.143 | 0.130 | 0.488 | 1.000 |
| **GO:1901362** | organic cyclic compound biosynthetic process | BP | 5 | 0.971 | 1.142 | 0.115 | 0.476 | 1.000 |
| **GO:0010467** | gene expression | BP | 5 | 0.971 | 1.142 | 0.117 | 0.462 | 1.000 |
| **GO:0044271** | cellular nitrogen compound biosynthetic process | BP | 5 | 0.971 | 1.142 | 0.111 | 0.457 | 1.000 |
| **GO:0003676** | nucleic acid binding | MF | 5 | 0.971 | 1.141 | 0.110 | 0.446 | 1.000 |
| **GO:0031224** | intrinsic component of membrane | CC | 3 | 0.892 | 1.137 | 0.333 | 0.452 | 1.000 |
| **GO:0044249** | cellular biosynthetic process | BP | 5 | 0.971 | 1.137 | 0.092 | 0.440 | 1.000 |
| **GO:0034654** | nucleobase-containing compound biosynthetic process | BP | 5 | 0.971 | 1.136 | 0.127 | 0.431 | 1.000 |
| **GO:0046483** | heterocycle metabolic process | BP | 5 | 0.971 | 1.135 | 0.116 | 0.428 | 1.000 |
| **GO:0032774** | RNA biosynthetic process | BP | 5 | 0.971 | 1.132 | 0.106 | 0.424 | 1.000 |
| **GO:1901360** | organic cyclic compound metabolic process | BP | 5 | 0.971 | 1.130 | 0.104 | 0.420 | 1.000 |
| **GO:0016021** | integral component of membrane | CC | 3 | 0.892 | 1.122 | 0.366 | 0.442 | 1.000 |
| **GO:0016779** | nucleotidyltransferase activity | MF | 6 | 0.971 | 1.120 | 0.227 | 0.438 | 1.000 |
| **GO:0016772** | transferase activity, transferring phosphorus-containing groups | MF | 6 | 0.971 | 1.119 | 0.228 | 0.436 | 1.000 |
| **GO:0009987** | cellular process | BP | 6 | 0.971 | 1.118 | 0.098 | 0.437 | 1.000 |
| **GO:0097159** | organic cyclic compound binding | MF | 7 | 0.970 | 1.105 | 0.105 | 0.498 | 1.000 |

**Table S5.** Picorna-like viruses.

| Gene | Length | Domain hits | Accession match | Interval | E-value | Description | Max Score | Total Score | Query Cover | E value | Per. Ident | Acc. Len | Accession | No. ORFs |
|---|---|---|---|---|---|---|---|---|---|---|---|---|---|---|
| **g1894_i0** | 5679 | RNA_dep_RNAP | cd01699 | 313-1146 | 1.09E-57 | polyprotein [Soybean thrips picorna-like virus 8] | 4.01E+02 | 4.01E+02 | 99% | 3.00E-126 | 67.51% | 1.41E+03 | QQP18689.1 | 20 |
| | | RNA_helicase | pfam00910 | 3742-4083 | 3.70E-21 | polyprotein [Soybean thrips picorna-like virus 8] | 165 | 165 | 99% | 2.00E-44 | 59.29% | 1406 | QQP18689.1 | 6 |
| **g20928_i0** | 747 | CRPV_capsid superfamily | cl07393 | 68-304 | 2.78E-08 | structural polyprotein [Dicistroviridae sp.] | 78.6 | 78.6 | 98% | 1.00E-14 | 51.95% | 891 | QJI52023.1 | 4 |
| **g2275_i0** | 5222 | RNA_dep_RNAP | cd01699 | 327-1190 | 1.79E-59 | polyprotein [Soybean thrips iflavirus 9] | 356 | 356 | 99% | 5.00E-108 | 58.56% | 2773 | QQX28925.1 | 14 |
| | | RNA_helicase | pfam00910 | 4281-4607 | 1.05E-26 | polyprotein [Iflaviridae sp.] | 202 | 202 | 99% | 2.00E-57 | 84.26% | 2974 | ULF99903.1 | 10 |
| | | Peptidase_C3 superfamily | cl02893 | 1824-2357 | 3.28E-07 | polyprotein [Iflaviridae sp.] | 145 | 145 | 96% | 2.00E-36 | 41.62% | 2974 | ULF99903.1 | 9 |
| | | RING-HC | cd16449 | 3465-3587 | 2.19E-04 | polyprotein [Iflaviridae sp.] | 41.2 | 41.2 | 96% | 39 | 43.90% | 3018 | QKN89072.1 | 1 |
| **g2855_i0** | 4620 | RNA_helicase | pfam00910 | 247-591 | 1.99E-17 | hypothetical protein 1 [Dicistroviridae sp.] | 123 | 123 | 99% | 8.00E-30 | 55.26% | 1875 | ULF99784.1 | 10 |
| | | RNA_dep_RNAP | cd01699 | 3302-4171 | 1.83E-78 | RNA dependent RNA polymerase [Picornaviridae sp.] | 383 | 383 | 95% | 1.00E-125 | 62.95% | 663 | URG14911.1 | 18 |
| **g2855_i1** | 4065 | RNA_dep_RNAP | cd01699 | 1803-2672 | 1.47E-78 | RNA dependent RNA polymerase [Picornaviridae sp.] | 383 | 383 | 95% | 1.00E-125 | 62.95% | 663 | URG14911.1 | 18 |
| | | rhv_like | cd00205 | 792-1295 | 1.24E-20 | hypothetical protein [Dicistroviridae sp.] | 147 | 147 | 98% | 5.00E-37 | 43.45% | 2862 | ULF99798.1 | 16 |
| | | rhv_like | cd00205 | 6-350 | 6.00E-13 | hypothetical protein 2 [Beihai | 74.7 | 74.7 | 96% | 1.00E-12 | 32.74% | 1012 | YP_009333387.1 | 9 |

| | | | | | | | | | | | | | | |
|---|---|---|---|---|---|---|---|---|---|---|---|---|---|---|
| | | | | | | picorna-like virus 75] | | | | | | | | |
| | | Dicistro_VP4 super family | cl13011 | 546-683 | 9.48E-06 | hypothetical protein 2 [Wenling picorna-like virus 2] | 55.5 | 55.5 | 99% | 5.00E-07 | 53.33% | 945 | YP_009333181.1 | 5 |
| **g2855_i3** | 2739 | RNA_dep_RNAP | cd01699 | 1421-2290 | 3.68E-80 | RNA dependent RNA polymerase [Picornaviridae sp.] | 383 | 383 | 95% | 1.00E-125 | 62.95% | 663 | URG14911.1 | 18 |
| **g3684_i0** | 3984 | RNA_dep_RNAP | cd01699 | 3109-3930 | 1.19E-78 | RNA dependent RNA polymerase [Picornaviridae sp.] | 363 | 363 | 99% | 4.00E-118 | 60.81% | 663 | URG14911.1 | 18 |
| | | rhv_like | cd00205 | 1120-1593 | 4.55E-15 | polyprotein [Varroa dicistrovirus 1] | 123 | 123 | 78% | 6.00E-29 | 41.79% | 2979 | UCR92472.1 | 9 |
| | | rhv_like | cd00205 | 2080-2571 | 3.28E-14 | hypothetical protein [Dicistroviridae sp.] | 144 | 144 | 99% | 4.00E-36 | 46.06% | 2862 | ULF99798.1 | 11 |
| | | CRPV_capsid super family | cl07393 | 319-921 | 4.00E-09 | capsid protein [Guiyang dicistrovirus 4] | 79.3 | 79.3 | 98% | 5.00E-13 | 33.33% | 1010 | UHR49777.1 | 13 |
| | | Dicistro_VP4 super family | cl13011 | 1795-1929 | 2.34E-05 | putative structural polyprotein [Solenopsis invicta virus 13] | 43.5 | 43.5 | 81% | 6 | 41.86% | 835 | QBL75905.1 | 2 |
| **g3814_i0** | 3900 | RT_like super family | cl02808 | 3210-3899 | 1.87E-32 | RNA dependent RNA polymerase [Picornaviridae sp.] | 291 | 291 | 99% | 4.00E-87 | 58.44% | 1242 | URG15016.1 | 14 |
| | | Calici_coat super family | cl03015 | 2334-2921 | 1.20E-11 | calicivirus coat protein [PNG bee virus 12] | 186 | 186 | 86% | 3.00E-52 | 52.63% | 556 | QKW94216.1 | 12 |
| **g5994_i0** | 2787 | RNA_dep_RNAP | cd01699 | 451-1323 | 1.93E-56 | polyprotein [Bat iflavirus] | 494 | 494 | 99% | 4.00E-156 | 79.93% | 3179 | YP_009345906.1 | 14 |
| **g5994_i2** | 2334 | RNA_dep_RNAP | cd01699 | 452-1324 | 5.51E-57 | polyprotein [Bat iflavirus] | 494 | 494 | 99% | 4.00E-156 | 79.93% | 3179 | YP_009345906.1 | 14 |
| **g6982_i4** | 1572 | rhv_like super family | cl13999 | 1022-1273 | 8.66E-03 | Chain C, Structural polyprotein, VP2 [Israeli acute paralysis virus] | 92.8 | 92.8 | 99% | 5.00E-21 | 52.38% | 200 | 5CDD_C | 4 |

| | | | | | | | | | | | | | | |
|---|---|---|---|---|---|---|---|---|---|---|---|---|---|---|
| | | RT_like super family | cl02808 | 3-401 | 3.21E-31 | non-structural polyprotein [Linepithema humile virus 1] | 239 | 239 | 99% | 5.00E-70 | 82.58% | 1930 | AWK91473.1 | 6 |

**Table S6.** Narna-like virus.

| Gene | Length | Description | Max Score | Total Score | Query Cover | e-value | Per. Ident | Acc. Len | Accession |
|---|---|---|---|---|---|---|---|---|---|
| **g9141_i0** | 1862 | Guiyang Paspalum thunbergii narna-like virus 1 | 444 | 679 | 90% | 8.00E-178 | 65.59% | 1072 | UUW20993.1 |

**Table S7.** RdRp sequences used for phylogenetic analysis for picornavirales.

| Source | Family | Genus | Virus name | Abbreviation | Domain | GenBank accession number |
|---|---|---|---|---|---|---|
| This study | Unclassified Iflaviridae | Unclassified Iflaviridae | Putative A. octoarticulatus virus 1 (g2275) | AoV1 | RdRp | NA |
| This study | Unclassified Iflaviridae | Unclassified Iflaviridae | Putative A. octoarticulatus virus 2 (g5994) | AoV2 | RdRp | NA |
| NCBI | Iflaviridae | Unclassified Iflavirus | Soybean thrips iflavirus 9 | STV9 | RdRp | QQX28925.1 |
| NCBI | Iflaviridae | Unclassified Iflaviridae | Solenopsis invicta virus 11 | SINV-11 | RdRp | QBL75900.1 |
| NCBI | Iflaviridae | Unclassified Iflaviridae | Solenopsis invicta virus 16 | SINV-16 | RdRp | MT860233.1 |
| NCBI | Iflaviridae | Unclassified Iflaviridae | Solenopsis invicta virus 17 | SINV-17 | RdRp | MT860234.1 |
| NCBI | Iflaviridae | Unclassified Iflaviridae | King virus | KgV | RdRp | KX779454.1 |
| NCBI | Iflaviridae | Unclassified Iflavirus | Formica exsecta virus 2 | FeV2 | RdRp | KF500002 |
| NCBI | Iflaviridae | Unclassified Iflaviridae | Thaumetopoea pityocampa iflavirus 1 | TpIV1 | RdRp | KP217032 |
| NCBI | Iflaviridae | Unclassified Iflaviridae | Nilaparvata lugens honeydew virus-2 | NlHV2 | RdRp | AB826459.1 |
| NCBI | Iflaviridae | Unclassified Iflaviridae | Osiphanes invirae iflavirus 1 | OiIV1 | RdRp | KR534892 |
| NCBI | Iflaviridae | Unclassified Iflavirus | Bat iflavirus (Yinda, et al., 2016) | NA | RdRp | YP_009345906.1 |
| NCBI | Iflaviridae | Iflavirus | Antheraea pernyi iflavirus | LnApIV | RdRp | KF751885 |
| NCBI | Iflaviridae | Iflavirus | Brevicoryne brassicae virus | BBV | RdRp | EF517277 |
| NCBI | Iflaviridae | Iflavirus | Deformed wing virus | DWV | RdRp | AJ489744 |
| NCBI | Iflaviridae | Iflavirus | Dinocampus coccinellae paralysis virus | DcPV | RdRp | KF843822 |
| NCBI | Iflaviridae | Iflavirus | Ectropis obliqua virus | EoV | RdRp | AY365064 |
| NCBI | Iflaviridae | Iflavirus | Infectious flacherie virus | IFV | RdRp | AB000906 |
| NCBI | Iflaviridae | Iflavirus | Lymantria dispar iflavirus 1 | LdIV1 | RdRp | KJ629170 |
| NCBI | Iflaviridae | Iflavirus | Nilaparvata lugens honeydew virus 1 | NLHV1 | RdRp | AB766259 |
| NCBI | Iflaviridae | Iflavirus | Perina nuda virus | PnV | RdRp | AF323747 |
| NCBI | Iflaviridae | Iflavirus | Sacbrood virus | SBV | RdRp | AF092924 |
| NCBI | Iflaviridae | Iflavirus | Slow bee paralysis virus | SBPV | RdRp | EU035616 |
| NCBI | Iflaviridae | Iflavirus | Spodoptera exigua iflavirus 1 | SEIV1 | RdRp | JN091707 |

| | | | | | | |
|---|---|---|---|---|---|---|
| NCBI | Iflaviridae | Iflavirus | Spodoptera exigua iflavirus 2 | SEIV2 | RdRp | JN870848 |
| NCBI | Iflaviridae | Iflavirus | Varroa destructor virus 1 | VDV1 | RdRp | AY251269 |
| This study | Dicistroviridae | Unclassified Dicistroviridae | Putative A. octoarticulatus virus 3 (g6982) | AoV3 | RdRp | NA |
| NCBI | Dicistroviridae | Aparavirus | Kashmir bee virus | KBV | RdRp | AHL83499 |
| NCBI | Dicistroviridae | Aparavirus | Acute bee paralysis virus | ABPV | RdRp | AAN63803 |
| NCBI | Dicistroviridae | Aparavirus | Israeli acute paralysis virus | IAPV | RdRp | ACD01399 |
| NCBI | Dicistroviridae | Aparavirus | Solenopsis invicta virus 1 | SINV-1 | RdRp | AAU85375 |
| NCBI | Dicistroviridae | Aparavirus (Gruber et al., 2017) | Formica exsecta virus 1 | FEX-1 | RdRp | KF500001.1 |
| NCBI | Dicistroviridae | Aparavirus (Gruber et al., 2017) | Linepithema humile virus 1 | LHUV-1 | RdRp | MH213249.1 |
| NCBI | Dicistroviridae | Cripavirus | Aphid lethal paralysis virus | ALPV | RdRp | AF536531.1 |
| NCBI | Dicistroviridae | Triatovirus | Black queen cell virus | BQCV | RdRp | NP_620564 |
| This study | Unclassified Picornaviridae | Unclassified Picornaviridae | Putative A. octoarticulatus virus 4 (g1894) | AoV4 | RdRp | NA |
| This study | Unclassified Picornaviridae | Unclassified Picornaviridae | Putative A. octoarticulatus virus 5 (g2855) | AoV5 | RdRp | NA |
| This study | Unclassified Picornaviridae | Unclassified Picornaviridae | Putative A. octoarticulatus virus 6 (g3684) | AoV6 | RdRp | NA |
| This study | Unclassified Picornaviridae | Unclassified Picornaviridae | Putative A. octoarticulatus virus 7 (g3814) | AoV7 | RdRp | NA |
| NCBI | Picornaviridae | Enterovirus | Human poliovirus 1 Mahoney | POLIO1A | RdRp | V01149 |
| NCBI | Picornaviridae | Sapelovirus | Porcine enterovirus 8 | PEV8 | RdRp | AF406813 |
| NCBI | Picornaviridae | Unassigned: Shanbavirus | Bat picornavirus | | RdRp | KJ641694 |
| NCBI | Picornaviridae | Aphthovirus | Foot and mouth disease virus | FMDV-O1K | RdRp | X00871 |
| NCBI | Picornaviridae | Erbovirus | Equine rhinitis B virus 1 | | RdRp | X96871 |
| NCBI | Picornaviridae | Mosavirus | Mosavirus A1 | | RdRp | JF973687 |
| NCBI | Picornaviridae | Teschovirus | Teschovirus A1 | | RdRp | AF231769 |
| NCBI | Picornaviridae | Hunnivirus | Bovine hungarovirus 1 | BHuV-1 | RdRp | JQ941880 |

| | | | | | | |
|---|---|---|---|---|---|---|
| NCBI | Picornaviridae | Cardiovirus | Cardiovirus A1 | | RdRp | M81861 |
| NCBI | Picornaviridae | Senecavirus | Senecavirus A1 | | RdRp | DQ641257 |
| NCBI | Picornaviridae | Mischivirus | Miniopterus schreibersii picornavirus 1 | | RdRp | JQ814851 |
| NCBI | Picornaviridae | Cosavirus | Cosavirus A1 | | RdRp | FJ438902 |
| NCBI | Picornaviridae | Avisivirus | Turkey avisivirus | | RdRp | KC465954 |
| NCBI | Picornaviridae | Avihepatovirus | Duck hepatitis A virus 1 | DHAV-1 | RdRp | DQ249299 |
| NCBI | Picornaviridae | Pasivirus | Pasivirus A1 | | RdRp | JQ316470 |
| NCBI | Picornaviridae | Kunsagivirus | Kunsagivirus A1 | | RdRp | KC935379 |
| NCBI | Picornaviridae | Aquamavirus | Aquamavirus A1 | SePV1 | RdRp | EU142040 |
| NCBI | Picornaviridae | Parechovirus | Human parechovirus 1 | | RdRp | L02971 |
| NCBI | Picornaviridae | Potamipivirus | Eel picornavirus 1 | | RdRp | KC843627 |
| NCBI | Picornaviridae | Limnipivirus | Limnipivirus A1 | | RdRp | JX134222 |
| NCBI | Picornaviridae | Tremovirus | Tremovirus A1 | | RdRp | AJ225173 |
| NCBI | Picornaviridae | Hepatovirus | Hepatovirus A | | RdRp | M14707 |
| NCBI | Picornaviridae | Salivirus | Salivirus NG-J1 | | RdRp | GQ179640 |
| NCBI | Picornaviridae | Kobuvirus | Aichi virus 1 | | RdRp | AB010145 |
| NCBI | Picornaviridae | Sakobuvirus | Feline sakobuvirus A | | RdRp | KF387721 |
| NCBI | Picornaviridae | Passerivirus | Passerivirus A1 | | RdRp | GU182406 |
| NCBI | Picornaviridae | Gallivirus | Gallivirus A1 | | RdRp | JQ691613 |
| NCBI | Picornaviridae | Oscivirus | Oscivirus A1 | | RdRp | GU182408 |
| NCBI | Picornaviridae | Dicipivirus | Canine picodicistrovirus | | RdRp | JN819202 |
| NCBI | Picornaviridae | Rosavirus | Rosavirus M-7 | | RdRp | JF973686 |
| NCBI | Picornaviridae | Megrivirus | Turkey hepatitis virus 2993D | | RdRp | HM751199 |
| NCBI | Picornaviridae | Sicinivirus | Sicinivirus A | | RdRp | KF741227 |
| NCBI | Secoviridae | Comovirus | Cowpea mosaic virus | | RdRp | NC_003549.1 |
| NCBI | Secoviridae | Nepovirus | Arabis mosaic virus | | RdRp | NC_006057 |

| | | | | | | |
|---|---|---|---|---|---|---|
| NCBI | Secoviridae | Cheravirus | Cherry rasp leaf virus | | RdRp | NC_006271.1 |

**Table S8.** RdRp sequences used for phylogenetic analysis of Narna-like viruses.

| Source | Family | Genus | Virus name | Domain | GenBank accession number | Link-BLAST |
|---|---|---|---|---|---|---|
| This study | Narnaviridae | unclassified Narnaviridae | Putative A. octoarticulatus narna-like virus | RdRp | NA | NA |
| NCBI | Narnaviridae | unclassified Narnaviridae | Linepithema humile narna-like virus 1 | RdRp | MH213236 | https://www.ncbi.nlm.nih.gov/nuccore/MH213236 |
| NCBI | Narnaviridae | unclassified Narnaviridae | Insect narna-like virus 1 | RdRp | MW297845.1 | https://www.ncbi.nlm.nih.gov/nuccore/MW297845.1 |
| NCBI | Narnaviridae | unclassified Narnavirus | Drosophila-associated narnavirus 1 | RdRp | MZ852366.1 | https://www.ncbi.nlm.nih.gov/nuccore/2160075275 |
| NCBI | Narnaviridae | unclassified Narnaviridae | Culex narnavirus 1 | RdRp | OM817537.1 | https://www.ncbi.nlm.nih.gov/nuccore/OM817537.1 |
| NCBI | Narnaviridae | Narnavirus | Leptomonas Narna-like virus 1 | RdRp | KY628364.1 | https://www.ncbi.nlm.nih.gov/nuccore/KY628364.1 |
| NCBI | Narnaviridae | Narnavirus | Blechmonas luni narnavirus 1 | RdRp | NC_040829.1 | https://www.ncbi.nlm.nih.gov/nuccore/NC_040829.1 |
| NCBI | Narnaviridae | Narnavirus | Aedes angustivittatus narnavirus | RdRp | MK285331.1 | https://www.ncbi.nlm.nih.gov/nuccore/1567319223 |
| NCBI | Narnaviridae | unclassified Narnaviridae | Guiyang Paspalum thunbergii narna-like virus 1 | RdRp | OM514596.1 | https://www.ncbi.nlm.nih.gov/nuccore/2284931872 |
| NCBI | Narnaviridae | Narnavirus | Saccharomyces 23S RNA narnavirus | RdRp | MW476518.1 | https://www.ncbi.nlm.nih.gov/nuccore/MW476518.1 |
| NCBI | Narnaviridae | Narnavirus | Erysiphe necator associated narnavirus 27 | RdRp | MN605440.1 | https://www.ncbi.nlm.nih.gov/nuccore/1841985104 |
| NCBI | Botourmiaviridae | Ourmiavirus | Cassava virus | RdRp | FJ157981.1 | https://www.ncbi.nlm.nih.gov/nuccore/FJ157981.1 |

**Table S9.** Analysis and annotation of genes differentially expressed between high-quality ant bodyguards and broodcare workers.

| | baseMean | log2FoldChange | lfcSE | stat | pvalue | padj | Annotation (Blast and InterProScan) |
|---|---|---|---|---|---|---|---|
| **g5823** | 1006.862 | -1.940 | 0.164 | -11.817 | 3.18E-32 | 6.21E-28 | Gibberellin-regulated protein 14 |
| **g123** | 424.790 | -2.426 | 0.217 | -11.188 | 4.65E-29 | 4.53E-25 | CHIT1 protein |
| **g11825** | 245.035 | -3.127 | 0.288 | -10.842 | 2.19E-27 | 1.42E-23 | Histone deacetylase HDT2 |
| **g9792** | 2319.680 | -3.012 | 0.285 | -10.567 | 4.22E-26 | 2.06E-22 | Z512B protein |
| **g1707** | 175.503 | -2.530 | 0.247 | -10.263 | 1.03E-24 | 4.03E-21 | 65471CHS2_CAEEL Chitin synthase chs-2 {ECO:0000305} |
| **g5273** | 121.896 | -2.545 | 0.275 | -9.272 | 1.83E-20 | 5.95E-17 | PREDICTED: uncharacterized protein LOC105449918 |
| **g8064** | 445.782 | -2.983 | 0.344 | -8.669 | 4.34E-18 | 1.21E-14 | PREDICTED: uncharacterized protein LOC105456783 |
| **g3224** | 895.969 | -2.066 | 0.239 | -8.644 | 5.42E-18 | 1.32E-14 | Chitin deacetylase 1 isoform X1 |
| **g3043** | 725.979 | -1.768 | 0.219 | -8.067 | 7.19E-16 | 1.56E-12 | Zinc finger CCCH domain-containing protein 13 |
| **g5480** | 222.793 | -2.867 | 0.359 | -7.992 | 1.33E-15 | 2.59E-12 | Probable serine/threonine-protein kinase clkA |
| **g9996** | 125.271 | -3.282 | 0.417 | -7.877 | 3.36E-15 | 5.95E-12 | Early nodulin-75 |
| **g23784** | 215.102 | -2.211 | 0.287 | -7.707 | 1.29E-14 | 2.06E-11 | ---NA--- |
| **g1** | 2254.201 | -1.092 | 0.142 | -7.699 | 1.37E-14 | 2.06E-11 | FBN2 protein |
| **g6663** | 680.124 | -2.763 | 0.364 | -7.596 | 3.06E-14 | 4.27E-11 | Spore wall protein 1 isoform X1 |
| **g16911** | 87.699 | -2.266 | 0.301 | -7.540 | 4.71E-14 | 6.13E-11 | PREDICTED: uncharacterized protein LOC105453081 |
| **g4947** | 1169.374 | -1.800 | 0.240 | -7.514 | 5.73E-14 | 6.99E-11 | Chitin deacetylase |
| **g4305** | 614.913 | -1.662 | 0.224 | -7.411 | 1.26E-13 | 1.44E-10 | Chondroitin proteoglycan-2-like |
| **g4614** | 6277.228 | -1.612 | 0.226 | -7.146 | 8.95E-13 | 9.70E-10 | Lysosomal alpha-mannosidase |
| **g4967** | 198.421 | -2.889 | 0.405 | -7.127 | 1.02E-12 | 1.05E-09 | Neuronal pentraxin-2 |
| **g4715** | 476.850 | -1.728 | 0.246 | -7.017 | 2.26E-12 | 2.21E-09 | PREDICTED: uncharacterized protein LOC105449142 |
| **g25699** | 1127.747 | -0.858 | 0.124 | -6.933 | 4.13E-12 | 3.84E-09 | U6-MYRTX-Tb1a precursor |
| **g11369** | 129.691 | -3.042 | 0.451 | -6.751 | 1.47E-11 | 1.30E-08 | Mediator of RNA polymerase II transcription subunit 15-like |
| **g2926** | 138.442 | -2.161 | 0.327 | -6.617 | 3.67E-11 | 3.12E-08 | Nose resistant to fluoxetine protein 6 |

| | | | | | | | |
|---|---|---|---|---|---|---|---|
| **g4166** | 951.296 | -1.750 | 0.270 | -6.486 | 8.79E-11 | 6.98E-08 | Probable chitinase 3 |
| **g869** | 1161.714 | -1.411 | 0.218 | -6.484 | 8.94E-11 | 6.98E-08 | Mucin-5AC |
| **g13788** | 186.909 | -5.473 | 0.853 | -6.416 | 1.40E-10 | 1.05E-07 | Cuticle protein 64-like |
| **g2622** | 140.697 | -1.612 | 0.252 | -6.398 | 1.58E-10 | 1.14E-07 | 63 kDa sperm flagellar membrane protein |
| **g7906** | 321.605 | -1.575 | 0.247 | -6.386 | 1.70E-10 | 1.19E-07 | Uncharacterized protein DBV15_06052 |
| **g829** | 649.529 | -1.067 | 0.171 | -6.257 | 3.91E-10 | 2.63E-07 | Low-density lipoprotein receptor-related protein 1B |
| **g3958** | 89.814 | -2.880 | 0.462 | -6.229 | 4.69E-10 | 3.05E-07 | Peroxidase isoform X2 |
| **g20498** | 128.992 | -2.508 | 0.403 | -6.223 | 4.87E-10 | 3.07E-07 | ---NA--- |
| **g6591** | 36369.092 | -3.409 | 0.555 | -6.144 | 8.04E-10 | 4.90E-07 | Cuticle protein 64-like isoform X2 |
| **g12874** | 5170.721 | -0.980 | 0.160 | -6.122 | 9.25E-10 | 5.47E-07 | Probable salivary secreted peptide |
| **g13057** | 63.382 | -2.404 | 0.394 | -6.099 | 1.07E-09 | 6.12E-07 | Pupal cuticle protein-like |
| **g9809** | 4569.086 | -1.801 | 0.296 | -6.088 | 1.14E-09 | 6.38E-07 | Trypsin-1-like isoform X1 |
| **g26724** | 278.093 | -1.404 | 0.233 | -6.018 | 1.77E-09 | 9.48E-07 | Venom peptide precursor ECTX1-Rm64a |
| **g8173** | 152.164 | -2.381 | 0.396 | -6.015 | 1.80E-09 | 9.48E-07 | Putative mediator of RNA polymerase II transcription subunit 12 isoform X1 |
| **g7529** | 75.012 | -3.778 | 0.630 | -6.000 | 1.98E-09 | 9.89E-07 | Arylphorin subunit alpha-like |
| **g10119** | 177.206 | -2.930 | 0.488 | -5.998 | 2.00E-09 | 9.89E-07 | PE44 protein |
| **g6983** | 6733.981 | -1.280 | 0.214 | -5.996 | 2.03E-09 | 9.89E-07 | Arylphorin subunit alpha-like |
| **g5483** | 595.861 | -1.937 | 0.325 | -5.954 | 2.61E-09 | 1.24E-06 | ---NA--- |
| **g11760** | 658.109 | -3.666 | 0.617 | -5.945 | 2.77E-09 | 1.29E-06 | Hypothetical protein X777_15926 |
| **g12056** | 178.283 | -2.247 | 0.383 | -5.862 | 4.56E-09 | 2.07E-06 | Uncharacterized protein LOC120358463 |
| **g8459** | 77.171 | -2.149 | 0.367 | -5.853 | 4.82E-09 | 2.14E-06 | Pancreatic lipase-related protein 2-like |
| **g3892** | 6257.992 | -1.332 | 0.228 | -5.834 | 5.40E-09 | 2.34E-06 | PREDICTED: uncharacterized protein LOC108768620 |
| **g4981** | 425.633 | -2.257 | 0.387 | -5.825 | 5.70E-09 | 2.42E-06 | PREDICTED: uncharacterized protein LOC105449179 |
| **g3978** | 471.683 | -0.915 | 0.159 | -5.752 | 8.82E-09 | 3.66E-06 | Phosphoenolpyruvate carboxykinase [GTP]-like |
| **g1077** | 324.697 | -2.105 | 0.367 | -5.743 | 9.29E-09 | 3.78E-06 | Cadherin-89D isoform X2 |
| **g8463** | 6150.111 | 1.733 | 0.303 | 5.726 | 1.03E-08 | 4.08E-06 | Lipase member H-like |

| | | | | | | | |
|---|---|---|---|---|---|---|---|
| **g482** | 134.952 | -2.379 | 0.418 | -5.689 | 1.28E-08 | 5.00E-06 | SP63 protein |
| **g26067** | 59.560 | -2.817 | 0.496 | -5.681 | 1.34E-08 | 5.12E-06 | ---NA--- |
| **g7216** | 22658.250 | -3.060 | 0.542 | -5.642 | 1.68E-08 | 6.30E-06 | Cuticle protein 67-like |
| **g6490** | 922.306 | -1.190 | 0.212 | -5.621 | 1.90E-08 | 7.00E-06 | Venom acid phosphatase Acph-1-like |
| **g9196** | 20996.331 | 1.471 | 0.263 | 5.594 | 2.22E-08 | 8.02E-06 | Zinc metalloproteinase nas-4-like |
| **g66062** | 87.767 | -1.694 | 0.303 | -5.581 | 2.40E-08 | 8.51E-06 | PREDICTED: uncharacterized protein LOC105461227, partial |
| **g2226** | 125.839 | -6.029 | 1.082 | -5.574 | 2.48E-08 | 8.57E-06 | Fibril-forming collagen alpha chain-like |
| **g6602** | 11185.007 | -1.168 | 0.210 | -5.573 | 2.50E-08 | 8.57E-06 | Probable salivary secreted peptide |
| **g1048** | 11848.690 | -0.682 | 0.123 | -5.545 | 2.94E-08 | 9.89E-06 | Endocuticle structural glycoprotein SgAbd-1 |
| **g6897** | 80642.583 | 0.943 | 0.170 | 5.540 | 3.02E-08 | 9.95E-06 | ---NA--- |
| **g562** | 4775.242 | -1.277 | 0.231 | -5.538 | 3.06E-08 | 9.95E-06 | Collagen alpha-1(IV) chain |
| **g2759** | 589.643 | -0.943 | 0.171 | -5.507 | 3.64E-08 | 1.16E-05 | Sushi, von Willebrand factor type A, EGF and pentraxin domain-containing protein 1 |
| **g15993** | 45.725 | -2.404 | 0.437 | -5.499 | 3.82E-08 | 1.20E-05 | Ig-like V-type domain-containing protein ENSP00000329499-like protein |
| **g6874** | 16370.231 | 1.176 | 0.214 | 5.494 | 3.92E-08 | 1.22E-05 | Chymotrypsin-1-like |
| **g15782** | 522.084 | -1.387 | 0.253 | -5.481 | 4.23E-08 | 1.29E-05 | Uncharacterized protein LOC112454799 |
| **g13731** | 10552.550 | -4.354 | 0.795 | -5.477 | 4.33E-08 | 1.29E-05 | Tetra-peptide repeat homeobox protein 1 |
| **g1156** | 3987.571 | -1.320 | 0.241 | -5.475 | 4.37E-08 | 1.29E-05 | Collagen alpha-5(IV) chain |
| **g7620** | 497.297 | -1.776 | 0.325 | -5.471 | 4.48E-08 | 1.30E-05 | ---NA--- |
| **g17038** | 2102.037 | -4.223 | 0.774 | -5.455 | 4.89E-08 | 1.40E-05 | Protein G12 |
| **g2745** | 537.424 | -1.486 | 0.276 | -5.374 | 7.69E-08 | 2.17E-05 | Interference hedgehog-like isoform X4 |
| **g4234** | 240.803 | -1.212 | 0.226 | -5.365 | 8.10E-08 | 2.26E-05 | ATP-binding cassette sub-family G member 5 |
| **g26621** | 111.970 | -1.626 | 0.306 | -5.308 | 1.11E-07 | 3.00E-05 | Hypothetical protein ALC57_12870, partial |
| **g868** | 6028.188 | -1.079 | 0.203 | -5.308 | 1.11E-07 | 3.00E-05 | Alpha-2A adrenergic receptor |
| **g14873** | 17.415 | 20.705 | 3.909 | 5.297 | 1.18E-07 | 3.15E-05 | ---NA--- |

| | | | | | | | |
|---|---|---|---|---|---|---|---|
| **g164** | 120176.487 | 0.615 | 0.116 | 5.290 | 1.22E-07 | 3.23E-05 | Titin isoform X2 |
| **g9011** | 221.330 | 1.334 | 0.252 | 5.282 | 1.28E-07 | 3.32E-05 | ---NA--- |
| **g13235** | 55.915 | -2.683 | 0.515 | -5.207 | 1.92E-07 | 4.93E-05 | Putative mediator of RNA polymerase II transcription subunit 26 |
| **g21793** | 315.951 | -1.020 | 0.196 | -5.202 | 1.97E-07 | 4.99E-05 | Hypothetical protein ALC57_11718, partial |
| **g14080** | 990.751 | -1.600 | 0.308 | -5.198 | 2.02E-07 | 5.05E-05 | Venom allergen 3-like |
| **g33510** | 57.376 | -2.647 | 0.512 | -5.173 | 2.30E-07 | 5.64E-05 | ---NA--- |
| **g5301** | 74.984 | -2.427 | 0.469 | -5.172 | 2.31E-07 | 5.64E-05 | Putative mediator of RNA polymerase II transcription subunit 26 |
| **g658** | 599.182 | -1.426 | 0.278 | -5.128 | 2.93E-07 | 7.05E-05 | Protein Skeletor, isoforms B/C isoform X1 |
| **g24224** | 64084.039 | 0.846 | 0.165 | 5.112 | 3.19E-07 | 7.60E-05 | ---NA--- |
| **g6910** | 552.601 | -1.053 | 0.208 | -5.061 | 4.17E-07 | 9.80E-05 | Carbonic anhydrase-related protein 10 |
| **g3740** | 676.226 | -0.971 | 0.192 | -5.045 | 4.53E-07 | 0.000105196 | Zinc finger protein 260 isoform X1 |
| **g6509** | 1188.137 | -0.723 | 0.145 | -4.986 | 6.17E-07 | 0.000141761 | Superoxide dismutase [Cu-Zn]-like |
| **g13743** | 4593.527 | -5.127 | 1.030 | -4.979 | 6.41E-07 | 0.00014538 | A-kinase anchor protein 14-like |
| **g10981** | 167.686 | 1.253 | 0.252 | 4.973 | 6.59E-07 | 0.000147926 | PREDICTED: LOW QUALITY PROTEIN: uncharacterized protein LOC105462971 |
| **g13589** | 18936.586 | -3.948 | 0.799 | -4.942 | 7.73E-07 | 0.000171409 | Chondroitin sulfate glucuronyltransferase |
| **g8183** | 965.361 | -1.384 | 0.281 | -4.920 | 8.67E-07 | 0.000190083 | Beta-hexosaminidase subunit beta |
| **g5151** | 85545.981 | -3.484 | 0.709 | -4.916 | 8.82E-07 | 0.000191334 | Spidroin-1-like isoform X2 |
| **g8422** | 554.312 | -4.939 | 1.020 | -4.840 | 1.30E-06 | 0.000278198 | Cuticle protein 6 |
| **g6715** | 14136.603 | -0.741 | 0.154 | -4.825 | 1.40E-06 | 0.000296664 | Proteasome subunit beta type-5 |
| **g8438** | 318.235 | 1.014 | 0.210 | 4.822 | 1.42E-06 | 0.000298552 | Tyrosine-protein phosphatase non-receptor type 1 |
| **g15833** | 948.061 | -0.861 | 0.180 | -4.787 | 1.69E-06 | 0.000351349 | PREDICTED: uncharacterized protein LOC105452749 |
| **g413** | 26657.295 | 0.582 | 0.122 | 4.780 | 1.75E-06 | 0.000360219 | Ryanodine receptor 44F isoform X20 |
| **g1381** | 699.694 | -0.893 | 0.187 | -4.763 | 1.91E-06 | 0.000387274 | Homeobox protein 2 |
| **g508** | 1621.298 | -0.627 | 0.132 | -4.760 | 1.94E-06 | 0.000389972 | Probable serine/threonine-protein kinase dyrk2 isoform X1 |

| | | | | | | | |
|---|---|---|---|---|---|---|---|
| **g11844** | 60.700 | -4.099 | 0.867 | -4.726 | 2.29E-06 | 0.00045195 | probable chitinase 3 |
| **g41100** | 1753.542 | 0.790 | 0.168 | 4.704 | 2.55E-06 | 0.000497516 | ---NA--- |
| **g2273** | 1317.735 | -0.870 | 0.186 | -4.683 | 2.82E-06 | 0.000545315 | Alpha-N-acetylglucosaminidase |
| **g9305** | 5677.713 | -3.728 | 0.801 | -4.656 | 3.22E-06 | 0.000616566 | Cuticle protein 64-like |
| **g8242** | 145.898 | -1.419 | 0.305 | -4.650 | 3.33E-06 | 0.000630253 | Probable cytochrome P450 49a1 isoform X1 |
| **g29392** | 1691.234 | 1.047 | 0.227 | 4.613 | 3.97E-06 | 0.000744671 | Zinc metalloproteinase nas-4-like |
| **g329** | 438.973 | -1.019 | 0.222 | -4.594 | 4.34E-06 | 0.00080677 | Fibrillin-2-like |
| **g1838** | 424.018 | -0.683 | 0.149 | -4.591 | 4.41E-06 | 0.000812336 | Dual oxidase |
| **g26577** | 1076.587 | -1.138 | 0.250 | -4.552 | 5.32E-06 | 0.000970602 | Mariner Mos1 transposase |
| **g23139** | 9.268 | 6.370 | 1.406 | 4.530 | 5.90E-06 | 0.00106278 | Lymphoid-restricted membrane protein-like isoform X2 |
| **g10275** | 3083.960 | -1.217 | 0.269 | -4.529 | 5.94E-06 | 0.00106278 | ATP-dependent DNA helicase PIF1 |
| **g5675** | 845.813 | -0.711 | 0.157 | -4.518 | 6.24E-06 | 0.001106147 | PREDICTED: uncharacterized protein LOC105450202 |
| **g9315** | 84.150 | -2.749 | 0.609 | -4.515 | 6.34E-06 | 0.001115206 | Endochitinase isoform X2 |
| **g25995** | 4896.721 | -0.800 | 0.177 | -4.509 | 6.52E-06 | 0.00113687 | RecName: Full=U17-myrmicitoxin-Tb1f; Short=U17-MYRTX-Tb1f; Flags: Precursor |
| **g27808** | 101.574 | -2.720 | 0.606 | -4.492 | 7.06E-06 | 0.001219927 | Phospholipase A1-like |
| **g20799** | 553.618 | -0.636 | 0.143 | -4.446 | 8.74E-06 | 0.001496845 | Cytochrome P450 6k1-like |
| **g2732** | 513.649 | -0.972 | 0.219 | -4.439 | 9.06E-06 | 0.0015323 | Netrin receptor UNC5C isoform X1 |
| **g22293** | 82.251 | -2.502 | 0.564 | -4.437 | 9.11E-06 | 0.0015323 | Hypothetical protein G5I_05028 |
| **g10859** | 359.199 | -0.869 | 0.196 | -4.424 | 9.70E-06 | 0.001617125 | ---NA--- |
| **g8464** | 809.164 | -4.013 | 0.908 | -4.421 | 9.80E-06 | 0.001621219 | CRPI protein |
| **g3989** | 87.643 | -2.878 | 0.652 | -4.412 | 1.03E-05 | 0.001670814 | Chaoptin |
| **g38173** | 9.322 | -6.946 | 1.575 | -4.411 | 1.03E-05 | 0.001670814 | Acyl-CoA Delta(11) desaturase-like |
| **g15134** | 2546.349 | 0.666 | 0.151 | 4.406 | 1.05E-05 | 0.001695126 | Ejaculatory bulb-specific protein 3-like |
| **g1172** | 144.288 | -2.171 | 0.494 | -4.394 | 1.12E-05 | 0.001783545 | Y5729 protein |
| **g8245** | 1888.790 | -0.807 | 0.184 | -4.382 | 1.18E-05 | 0.001866855 | U6_MYRTX_Ta1a |

| | | | | | | | |
|---|---|---|---|---|---|---|---|
| **g5065** | 204.835 | -0.826 | 0.189 | -4.379 | 1.19E-05 | 0.001873505 | Cyclin-dependent kinase-like 4 |
| **g26785** | 1370.996 | 1.103 | 0.254 | 4.339 | 1.43E-05 | 0.00223799 | Zinc metalloproteinase nas-4-like |
| **g15701** | 69.184 | -1.642 | 0.382 | -4.294 | 1.75E-05 | 0.002692353 | Protein obstructor-E |
| **g20880** | 37688.225 | -1.065 | 0.249 | -4.270 | 1.95E-05 | 0.002951813 | ---NA--- |
| **g12996** | 125.460 | -1.675 | 0.393 | -4.263 | 2.02E-05 | 0.003030259 | Protein obstructor-E-like |
| **g6531** | 259.971 | 0.926 | 0.218 | 4.255 | 2.09E-05 | 0.003119098 | Protein takeout-like |
| **g14253** | 1136.959 | -0.725 | 0.171 | -4.245 | 2.19E-05 | 0.003236562 | Lipopolysaccharide-induced tumor necrosis factor-alpha factor homolog |
| **g8960** | 430.095 | -1.561 | 0.369 | -4.233 | 2.31E-05 | 0.003390801 | Major royal jelly protein 1-like |
| **g656** | 2488.164 | -0.610 | 0.145 | -4.204 | 2.62E-05 | 0.00378888 | Semaphorin-5B |
| **g14127** | 18.994 | 3.890 | 0.934 | 4.166 | 3.10E-05 | 0.004448777 | Copia protein |
| **g17637** | 30039.611 | -1.146 | 0.276 | -4.158 | 3.21E-05 | 0.004570894 | PH4H protein |
| **g17942** | 1834.078 | -0.944 | 0.227 | -4.151 | 3.31E-05 | 0.004682854 | PREDICTED: pilosulin-4-like |
| **g8148** | 90.975 | -2.613 | 0.632 | -4.134 | 3.56E-05 | 0.004999356 | PREDICTED: uncharacterized protein LOC105451758 |
| **g18584** | 74.627 | -2.958 | 0.717 | -4.128 | 3.66E-05 | 0.005070495 | Protein GVQW3-like |
| **g2940** | 541.111 | -0.820 | 0.199 | -4.128 | 3.66E-05 | 0.005070495 | Extended synaptotagmin-1 |
| **g683** | 208.751 | -0.932 | 0.227 | -4.115 | 3.87E-05 | 0.005320029 | Serine-rich adhesin for platelets-like |
| **g1423** | 5186.261 | 0.623 | 0.152 | 4.106 | 4.02E-05 | 0.005468547 | Interaptin isoform X2 |
| **g1495** | 1032.504 | -0.766 | 0.187 | -4.105 | 4.04E-05 | 0.005468547 | PREDICTED: uncharacterized protein LOC105459069 |
| **g48923** | 1142.174 | 0.763 | 0.186 | 4.095 | 4.22E-05 | 0.005685416 | Titin isoform X9 |
| **g34531** | 24.242 | -2.800 | 0.685 | -4.089 | 4.32E-05 | 0.005778525 | Phospholipase A1-like |
| **g32** | 5156.170 | -0.705 | 0.173 | -4.085 | 4.42E-05 | 0.005861525 | Fibrillin-2-like |
| **g27103** | 1454.749 | -2.809 | 0.689 | -4.077 | 4.57E-05 | 0.005981462 | U3_MYRTX_Ta1a |
| **g8108** | 25.096 | -4.307 | 1.057 | -4.074 | 4.63E-05 | 0.006018633 | PREDICTED: uncharacterized protein LOC105454030 isoform X1 |
| **g52588** | 4413.466 | -0.726 | 0.179 | -4.066 | 4.78E-05 | 0.00617846 | Coiled-coil domain-containing protein 8-like |
| **g10913** | 424.525 | -4.260 | 1.058 | -4.026 | 5.67E-05 | 0.007278351 | OR49B protein |

| | | | | | | | |
|---|---|---|---|---|---|---|---|
| **g1280** | 18443.979 | 0.609 | 0.152 | 3.998 | 6.38E-05 | 0.008066426 | ZC21C protein |
| **g5108** | 3275.188 | -2.128 | 0.532 | -3.997 | 6.41E-05 | 0.008066426 | Prisilkin-39-like isoform X1 |
| **g5073** | 2143.026 | -0.731 | 0.183 | -3.993 | 6.53E-05 | 0.008167157 | ---NA--- |
| **g17163** | 45472.023 | -0.929 | 0.233 | -3.984 | 6.78E-05 | 0.008372581 | ---NA--- |
| **g5191** | 1073.095 | 0.783 | 0.197 | 3.981 | 6.85E-05 | 0.008408978 | Lipase member H-like |
| **g1382** | 85.413 | -1.126 | 0.284 | -3.960 | 7.48E-05 | 0.009126163 | probable G-protein coupled receptor Mth-like 1 |
| **g6037** | 106.258 | -1.154 | 0.292 | -3.959 | 7.54E-05 | 0.009135647 | Zinc finger CCCH domain-containing protein 13 isoform X2 |
| **g31059** | 12.407 | -4.301 | 1.089 | -3.950 | 7.81E-05 | 0.009402155 | ---NA--- |

**Table S10.** Analysis and annotation of genes differentially expressed between low-quality ant bodyguards and broodcare workers.

| | baseMean | log2FoldChange | lfcSE | stat | pvalue | padj | Annotation (Blast and InterProScan) |
|---|---|---|---|---|---|---|---|
| **g10913** | 296.626 | -4.430 | 1.024 | -4.328 | 1.51E-05 | 0.001815064 | OR49B protein |
| **g10981** | 198.880 | 0.997 | 0.248 | 4.028 | 5.62E-05 | 0.005446093 | PREDICTED: LOW QUALITY PROTEIN: uncharacterized protein LOC105462971 |
| **g1172** | 129.009 | -3.968 | 0.486 | -8.161 | 3.32E-16 | 1.87E-13 | Y5729 protein |
| **g13589** | 9950.790 | -8.964 | 0.879 | -10.197 | 2.05E-24 | 3.46E-21 | Chondroitin sulfate glucuronyltransferase |
| **g13743** | 2528.935 | -8.101 | 0.925 | -8.754 | 2.05E-18 | 1.73E-15 | A-kinase anchor protein 14-like |
| **g1423** | 4429.463 | 0.977 | 0.169 | 5.795 | 6.85E-09 | 1.86E-06 | Interaptin isoform X2 |
| **g15134** | 2328.754 | 0.849 | 0.216 | 3.933 | 8.39E-05 | 0.00765447 | Ejaculatory bulb-specific protein 3-like |
| **g15701** | 47.340 | -1.618 | 0.412 | -3.929 | 8.53E-05 | 0.007738992 | Protein obstructor-E |
| **g22293** | 50.422 | -5.352 | 0.738 | -7.248 | 4.23E-13 | 1.70E-10 | Hypothetical protein G5I_05028 |
| **g32** | 3730.824 | -0.892 | 0.139 | -6.430 | 1.28E-10 | 4.39E-08 | Basement membrane-specific heparan sulfate proteoglycan core protein |
| **g329** | 406.100 | -1.061 | 0.249 | -4.270 | 1.96E-05 | 0.002244935 | Fibrillin-2-like |
| **g3740** | 477.579 | -1.133 | 0.206 | -5.505 | 3.68E-08 | 9.28E-06 | Zinc finger protein 260 isoform X1 |
| **g5108** | 1801.329 | -1.858 | 0.283 | -6.576 | 4.84E-11 | 1.70E-08 | Prisilkin-39-like isoform X1 |
| **g5151** | 63982.304 | -7.933 | 0.965 | -8.219 | 2.05E-16 | 1.19E-13 | Spidroin-1-like isoform X2 |
| **g5675** | 664.620 | -0.643 | 0.165 | -3.895 | 9.81E-05 | 0.00852802 | PREDICTED: uncharacterized protein LOC105450202 |
| **g8242** | 118.093 | -1.065 | 0.250 | -4.255 | 2.09E-05 | 0.002337073 | Probable cytochrome P450 49a1 isoform X1 |
| **g8960** | 264.769 | -2.488 | 0.284 | -8.747 | 2.19E-18 | 1.76E-15 | Major royal jelly protein 1-like |
| **g6509** | 1036.979 | -0.975 | 0.244 | -3.991 | 6.59E-05 | 0.006245458 | Superoxide dismutase [Cu-Zn]-like |
| **g9305** | 3443.106 | -6.906 | 0.354 | -19.495 | 1.21E-84 | 2.04E-80 | Cuticle protein 64-like |
| **g10119** | 125.992 | -4.796 | 0.530 | -9.058 | 1.33E-19 | 1.41E-16 | PE44 protein |
| **g1077** | 243.237 | -3.294 | 0.388 | -8.499 | 1.92E-17 | 1.29E-14 | Cadherin-89D isoform X2 |
| **g11369** | 106.519 | -4.115 | 0.458 | -8.983 | 2.65E-19 | 2.63E-16 | Mediator of RNA polymerase II transcription subunit 15-like |
| **g1156** | 2700.040 | -1.727 | 0.202 | -8.563 | 1.10E-17 | 8.42E-15 | Collagen alpha-5(IV) chain |
| **g11760** | 502.005 | -5.836 | 1.095 | -5.330 | 9.81E-08 | 2.30E-05 | Hypothetical protein X777_15926 |

| | | | | | | | |
|---|---|---|---|---|---|---|---|
| **g11825** | 269.661 | -4.706 | 0.514 | -9.160 | 5.19E-20 | 5.84E-17 | Histone deacetylase HDT2 |
| **g12056** | 145.318 | -3.545 | 0.400 | -8.872 | 7.19E-19 | 6.74E-16 | Uncharacterized protein LOC120358463 |
| **g123** | 251.955 | -3.486 | 0.373 | -9.351 | 8.69E-21 | 1.05E-17 | CHIT1 protein |
| **g13057** | 91.590 | -3.835 | 0.629 | -6.101 | 1.05E-09 | 3.23E-07 | Pupal cuticle protein-like |
| **g13235** | 38.841 | -3.011 | 0.589 | -5.111 | 3.21E-07 | 6.66E-05 | Putative mediator of RNA polymerase II transcription subunit 26 |
| **g13731** | 6613.007 | -8.767 | 0.872 | -10.050 | 9.17E-24 | 1.41E-20 | Tetra-peptide repeat homeobox protein 1 |
| **g13788** | 314.297 | -5.915 | 1.401 | -4.223 | 2.41E-05 | 0.002661437 | Cuticle protein 64-like |
| **g15993** | 32.488 | -2.189 | 0.550 | -3.976 | 6.99E-05 | 0.006585286 | Ig-like V-type domain-containing protein ENSP00000329499-like protein |
| **g16911** | 79.089 | -2.669 | 0.682 | -3.916 | 9.00E-05 | 0.007962579 | PREDICTED: uncharacterized protein LOC105453081 |
| **g1707** | 135.791 | -2.904 | 0.505 | -5.753 | 8.78E-09 | 2.35E-06 | 65471CHS2_CAEEL Chitin synthase chs-2 {ECO:0000305} |
| **g20498** | 96.839 | -4.043 | 0.484 | -8.350 | 6.80E-17 | 4.41E-14 | ---NA--- |
| **g23784** | 149.967 | -3.514 | 0.444 | -7.920 | 2.38E-15 | 1.22E-12 | ---NA--- |
| **g24224** | 62145.653 | 0.992 | 0.237 | 4.184 | 2.87E-05 | 0.003084681 | ---NA--- |
| **g26067** | 87.702 | -4.655 | 1.143 | -4.074 | 4.61E-05 | 0.004634344 | ---NA--- |
| **g2622** | 90.920 | -1.760 | 0.301 | -5.839 | 5.26E-09 | 1.45E-06 | 63 kDa sperm flagellar membrane protein |
| **g2745** | 316.980 | -1.214 | 0.310 | -3.920 | 8.85E-05 | 0.007906913 | Interference hedgehog-like isoform X4 |
| **g2759** | 433.042 | -1.362 | 0.254 | -5.364 | 8.12E-08 | 1.96E-05 | Sushi, von Willebrand factor type A, EGF and pentraxin domain-containing protein 1 |
| **g2926** | 90.283 | -2.998 | 0.435 | -6.884 | 5.82E-12 | 2.23E-09 | Nose resistant to fluoxetine protein 6 |
| **g3043** | 519.283 | -2.793 | 0.375 | -7.457 | 8.85E-14 | 3.93E-11 | Zinc finger CCCH domain-containing protein 13 |
| **g3224** | 642.492 | -3.146 | 0.377 | -8.346 | 7.07E-17 | 4.42E-14 | Chitin deacetylase 1 isoform X1 |
| **g3892** | 4596.575 | -1.376 | 0.250 | -5.495 | 3.90E-08 | 9.62E-06 | PREDICTED: uncharacterized protein LOC108768620 |
| **g3958** | 59.358 | -4.945 | 0.673 | -7.347 | 2.02E-13 | 8.33E-11 | Peroxidase isoform X2 |
| **g4166** | 727.183 | -2.610 | 0.295 | -8.858 | 8.13E-19 | 7.22E-16 | Probable chitinase 3 |
| **g4305** | 470.213 | -2.101 | 0.277 | -7.581 | 3.43E-14 | 1.65E-11 | Chondroitin proteoglycan-2-like |
| **g4614** | 4234.444 | -1.222 | 0.235 | -5.211 | 1.88E-07 | 4.06E-05 | Lysosomal alpha-mannosidase |
| **g4715** | 375.915 | -2.640 | 0.353 | -7.468 | 8.17E-14 | 3.72E-11 | PREDICTED: uncharacterized protein LOC105449142 |
| **g482** | 137.078 | -3.132 | 0.501 | -6.254 | 4.00E-10 | 1.32E-07 | SP63 protein |
| **g4947** | 919.707 | -3.571 | 0.572 | -6.240 | 4.36E-10 | 1.42E-07 | Chitin deacetylase |

| | | | | | | | |
|---|---|---|---|---|---|---|---|
| **g4967** | 159.644 | -4.101 | 0.493 | -8.310 | 9.55E-17 | 5.75E-14 | Neuronal pentraxin-2 |
| **g4981** | 239.973 | -3.303 | 0.306 | -10.810 | 3.07E-27 | 6.48E-24 | PREDICTED: uncharacterized protein LOC105449179 |
| **g5273** | 89.561 | -3.771 | 0.444 | -8.500 | 1.90E-17 | 1.29E-14 | PREDICTED: uncharacterized protein LOC105449918 |
| **g5301** | 64.427 | -3.703 | 0.625 | -5.923 | 3.16E-09 | 9.20E-07 | Putative mediator of RNA polymerase II transcription subunit 26 |
| **g5480** | 277.524 | -3.840 | 0.898 | -4.276 | 1.90E-05 | 0.002209539 | Probable serine/threonine-protein kinase clkA |
| **g562** | 3241.621 | -1.652 | 0.170 | -9.707 | 2.81E-22 | 3.95E-19 | Collagen alpha-1(IV) chain |
| **g5823** | 756.023 | -2.175 | 0.305 | -7.132 | 9.88E-13 | 3.87E-10 | Gibberellin-regulated protein 14 |
| **g658** | 535.395 | -1.326 | 0.339 | -3.914 | 9.06E-05 | 0.007965228 | Protein Skeletor, isoforms B/C isoform X1 |
| **g6591** | 29570.584 | -8.794 | 0.763 | -11.519 | 1.06E-30 | 2.56E-27 | Cuticle protein 64-like isoform X2 |
| **g6663** | 451.921 | -5.049 | 0.298 | -16.948 | 2.00E-64 | 1.12E-60 | Spore wall protein 1 isoform X1 |
| **g6897** | 77646.922 | 1.166 | 0.228 | 5.107 | 3.28E-07 | 6.66E-05 | ---NA--- |
| **g7216** | 17888.735 | -5.733 | 0.321 | -17.859 | 2.48E-71 | 2.09E-67 | Cuticle protein 67-like |
| **g7529** | 339.831 | -6.907 | 1.328 | -5.200 | 2.00E-07 | 4.26E-05 | Arylphorin subunit alpha-like |
| **g8064** | 299.190 | -3.607 | 0.379 | -9.512 | 1.87E-21 | 2.43E-18 | PREDICTED: uncharacterized protein LOC105456783 |
| **g8173** | 145.985 | -4.358 | 0.427 | -10.206 | 1.86E-24 | 3.46E-21 | Putative mediator of RNA polymerase II transcription subunit 12 isoform X1 |
| **g829** | 436.368 | -1.325 | 0.199 | -6.670 | 2.56E-11 | 9.20E-09 | Low-density lipoprotein receptor-related protein 1B |
| **g8459** | 71.756 | -2.038 | 0.410 | -4.968 | 6.77E-07 | 0.000117719 | Pancreatic lipase-related protein 2-like |
| **g8463** | 10033.948 | 1.850 | 0.314 | 5.885 | 3.98E-09 | 1.12E-06 | Lipase member H-like |
| **g868** | 5118.615 | -1.337 | 0.278 | -4.810 | 1.51E-06 | 0.000242558 | Alpha-2A adrenergic receptor |
| **g869** | 978.026 | -1.554 | 0.322 | -4.823 | 1.42E-06 | 0.000231811 | Mucin-5AC |
| **g9011** | 267.906 | 1.237 | 0.225 | 5.494 | 3.93E-08 | 9.62E-06 | ---NA--- |
| **g9196** | 25371.256 | 1.490 | 0.243 | 6.134 | 8.56E-10 | 2.67E-07 | Zinc metalloproteinase nas-4-like |
| **g9792** | 1526.529 | -4.695 | 0.357 | -13.154 | 1.62E-39 | 5.47E-36 | Z512B protein |
| **g9809** | 2759.608 | -1.512 | 0.318 | -4.751 | 2.03E-06 | 0.000307809 | Trypsin-1-like isoform X1 |
| **g9996** | 112.972 | -4.939 | 1.048 | -4.711 | 2.46E-06 | 0.000367644 | Early nodulin-75 |
| **g5389** | 794.403 | -4.084 | 0.292 | -13.995 | 1.68E-44 | 7.08E-41 | Mediator of RNA polymerase II transcription subunit 15-like |
| **g7219** | 301.872 | -7.339 | 0.616 | -11.916 | 9.74E-33 | 2.74E-29 | Alpha/beta-gliadin A-IV-like |
| **g1494** | 1579.863 | 1.148 | 0.135 | 8.521 | 1.59E-17 | 1.16E-14 | Sodium-dependent dopamine transporter isoform X1 |

| | | | | | | | |
|---|---|---|---|---|---|---|---|
| **g25125** | 76.242 | -2.951 | 0.363 | -8.134 | 4.16E-16 | 2.26E-13 | PREDICTED: uncharacterized protein LOC105457500 isoform X2 |
| **g17024** | 468.628 | -1.865 | 0.234 | -7.964 | 1.67E-15 | 8.81E-13 | ---NA--- |
| **g7743** | 11028.029 | 1.465 | 0.189 | 7.747 | 9.38E-15 | 4.65E-12 | Peptidylglycine alpha-hydroxylating monooxygenase |
| **g5511** | 1144.437 | -5.609 | 0.744 | -7.541 | 4.67E-14 | 2.19E-11 | Late embryogenesis abundant protein, group 3-like |
| **g10980** | 283.097 | 1.470 | 0.198 | 7.438 | 1.02E-13 | 4.43E-11 | Hypothetical protein ALC57_08594, partial |
| **g18527** | 145.045 | -3.422 | 0.464 | -7.378 | 1.61E-13 | 6.77E-11 | Keratin, type II cytoskeletal 1-like |
| **g643** | 3555.165 | -0.770 | 0.112 | -6.863 | 6.76E-12 | 2.54E-09 | Mitochondrial amidoxime-reducing component 1 |
| **g7372** | 998.250 | -0.844 | 0.126 | -6.698 | 2.12E-11 | 7.76E-09 | Cysteine-rich secretory protein 1-like |
| **g14156** | 141.290 | -2.510 | 0.391 | -6.424 | 1.33E-10 | 4.49E-08 | PREDICTED: uncharacterized protein LOC105457500 isoform X1 |
| **g68084** | 50.394 | 4.311 | 0.703 | 6.136 | 8.47E-10 | 2.67E-07 | Hymenoptaecin |
| **g4815** | 1067.689 | -0.904 | 0.148 | -6.093 | 1.11E-09 | 3.33E-07 | Endocuticle structural glycoprotein SgAbd-1 |
| **g8125** | 162.201 | -1.435 | 0.241 | -5.946 | 2.75E-09 | 8.15E-07 | DNA helicase |
| **g13531** | 719.926 | -3.760 | 0.636 | -5.911 | 3.41E-09 | 9.75E-07 | Endocuticle structural glycoprotein SgAbd-2-like |
| **g5767** | 416.159 | -6.049 | 1.068 | -5.663 | 1.49E-08 | 3.93E-06 | RNA polymerase II degradation factor 1-like |
| **g3245** | 156.871 | -1.351 | 0.245 | -5.507 | 3.66E-08 | 9.28E-06 | Sodium-dependent nutrient amino acid transporter 1-like isoform X1 |
| **g768** | 1884.357 | -0.778 | 0.141 | -5.506 | 3.67E-08 | 9.28E-06 | Integrin alpha-PS2 isoform X1 |
| **g19974** | 391.313 | 3.904 | 0.731 | 5.337 | 9.43E-08 | 2.24E-05 | HYTA protein |
| **g5161** | 157.208 | 1.386 | 0.263 | 5.262 | 1.43E-07 | 3.30E-05 | PREDICTED: uncharacterized protein LOC105451267 |
| **g2598** | 514.950 | 1.023 | 0.195 | 5.250 | 1.52E-07 | 3.48E-05 | Hypothetical protein ALC60_12428 |
| **g14166** | 1411.958 | 1.052 | 0.201 | 5.229 | 1.70E-07 | 3.83E-05 | Ejaculatory bulb-specific protein 3-like |
| **g8320** | 842.476 | 1.297 | 0.248 | 5.222 | 1.77E-07 | 3.93E-05 | Sodium-dependent dopamine transporter isoform X1 |
| **g6937** | 305.174 | 2.276 | 0.436 | 5.215 | 1.84E-07 | 4.03E-05 | PREDICTED: uncharacterized protein LOC105452884 |
| **g28489** | 468.730 | 1.252 | 0.241 | 5.184 | 2.17E-07 | 4.57E-05 | ---NA--- |
| **g523** | 1534.710 | 0.779 | 0.152 | 5.109 | 3.24E-07 | 6.66E-05 | Scavenger receptor class B member 1 |
| **g10104** | 65.403 | 2.086 | 0.409 | 5.102 | 3.36E-07 | 6.76E-05 | ---NA--- |
| **g35380** | 103.088 | 1.777 | 0.349 | 5.089 | 3.60E-07 | 7.15E-05 | ---NA--- |
| **g1078** | 158.820 | -1.221 | 0.241 | -5.078 | 3.81E-07 | 7.48E-05 | Protein jagged-1 |

| | | | | | | | |
|---|---|---|---|---|---|---|---|
| **g4107** | 17725.681 | 0.975 | 0.192 | 5.072 | 3.93E-07 | 7.62E-05 | PREDICTED: uncharacterized protein LOC105448412 isoform X2 |
| **g3884** | 64.878 | -2.157 | 0.426 | -5.067 | 4.04E-07 | 7.75E-05 | Hormone receptor 4 isoform X3 |
| **g4674** | 5482.494 | -0.771 | 0.153 | -5.028 | 4.96E-07 | 9.40E-05 | Circadian clock-controlled protein-like |
| **g18140** | 13.084 | -7.087 | 1.411 | -5.022 | 5.11E-07 | 9.58E-05 | ---NA--- |
| **g1039** | 2384.381 | 0.711 | 0.142 | 5.012 | 5.39E-07 | 9.99E-05 | Uricase |
| **g5791** | 2243.089 | -5.233 | 1.048 | -4.994 | 5.91E-07 | 0.000108367 | HEXA protein |
| **g13218** | 11.195 | -6.869 | 1.377 | -4.989 | 6.07E-07 | 0.000110106 | Fatty acid synthase-like |
| **g9189** | 55.234 | -2.725 | 0.547 | -4.983 | 6.26E-07 | 0.000112141 | Cytochrome P450 4C1-like |
| **g8104** | 24.785 | -6.174 | 1.239 | -4.981 | 6.31E-07 | 0.000112141 | PREDICTED: uncharacterized protein LOC105451516 |
| **g23888** | 139.799 | 1.478 | 0.297 | 4.971 | 6.67E-07 | 0.000117229 | Ejaculatory bulb-specific protein 3-like |
| **g19862** | 649.621 | 3.717 | 0.749 | 4.963 | 6.96E-07 | 0.000119776 | Hymenoptaecin |
| **g1628** | 1887.674 | 0.819 | 0.167 | 4.911 | 9.06E-07 | 0.000152917 | 28S ribosomal protein S10, mitochondrial |
| **g12963** | 90.352 | -2.071 | 0.426 | -4.865 | 1.15E-06 | 0.000191531 | Endocuticle structural protein SgAbd-6-like |
| **g12341** | 175.934 | -1.358 | 0.281 | -4.834 | 1.34E-06 | 0.000221519 | Cytochrome P450 4C1-like |
| **g4005** | 4410.407 | 0.995 | 0.207 | 4.816 | 1.47E-06 | 0.000238095 | Cytochrome P450 6A1-like |
| **g7255** | 1234.963 | -0.706 | 0.147 | -4.796 | 1.62E-06 | 0.000257823 | ---NA--- |
| **g30386** | 741.582 | 3.646 | 0.761 | 4.793 | 1.64E-06 | 0.000259222 | HYTA protein |
| **g8265** | 5808.445 | 0.909 | 0.190 | 4.789 | 1.67E-06 | 0.000261467 | Nuclear pore complex protein DDB_G0274915 isoform X1 |
| **g18224** | 37.990 | 2.053 | 0.429 | 4.780 | 1.75E-06 | 0.000271233 | Histone H2B |
| **g415** | 11471.095 | 0.883 | 0.186 | 4.754 | 2.00E-06 | 0.000306527 | PREDICTED: uncharacterized protein LOC105452570 isoform X3 |
| **g23003** | 16.940 | -4.988 | 1.058 | -4.713 | 2.44E-06 | 0.00036697 | ---NA--- |
| **g20826** | 282.780 | -1.173 | 0.250 | -4.695 | 2.67E-06 | 0.000395508 | DUF4817 domain-containing protein |
| **g18601** | 16.925 | 4.455 | 0.950 | 4.689 | 2.75E-06 | 0.000402205 | Putative nuclease HARBI1 isoform X2 |
| **g3171** | 3574.833 | 0.942 | 0.201 | 4.688 | 2.77E-06 | 0.000402205 | Serine protease easter-like isoform X1 |
| **g35603** | 348.107 | -2.124 | 0.455 | -4.671 | 3.00E-06 | 0.000432924 | ---NA--- |
| **g16054** | 1166.706 | 1.388 | 0.298 | 4.654 | 3.25E-06 | 0.000460987 | Zinc metalloproteinase nas-4-like |
| **g1142** | 183.950 | -1.066 | 0.230 | -4.642 | 3.46E-06 | 0.000485937 | Centromere-associated protein E-like |
| **g3023** | 6281.687 | 1.273 | 0.276 | 4.618 | 3.88E-06 | 0.000537581 | Ataxin-7-like protein 2 |

| | | | | | | | |
|---|---|---|---|---|---|---|---|
| **g3461** | 9000.369 | -1.018 | 0.221 | -4.617 | 3.89E-06 | 0.000537581 | Rhodopsin-like isoform X2 |
| **g9742** | 1269.228 | -1.221 | 0.265 | -4.610 | 4.03E-06 | 0.000552574 | Cartilage oligomeric matrix protein |
| **g13430** | 595.216 | -0.825 | 0.180 | -4.574 | 4.78E-06 | 0.000649908 | Cytochrome P450 4C1-like |
| **g37213** | 18.788 | -5.034 | 1.102 | -4.570 | 4.88E-06 | 0.000658171 | Transposable element Tc3 transposase |
| **g859** | 240.643 | -1.658 | 0.363 | -4.566 | 4.96E-06 | 0.000664248 | Putative chitinase 3 |
| **g1286** | 1004.993 | -0.854 | 0.188 | -4.534 | 5.78E-06 | 0.000768198 | SPON2 protein |
| **g9272** | 54.893 | -1.812 | 0.403 | -4.499 | 6.82E-06 | 0.000898743 | Retinol dehydrogenase 11 |
| **g6169** | 265.067 | -0.938 | 0.209 | -4.492 | 7.07E-06 | 0.000924841 | ---NA--- |
| **g1818** | 467.011 | -0.780 | 0.175 | -4.468 | 7.90E-06 | 0.001025812 | Brain-specific angiogenesis inhibitor 1-associated protein 2 |
| **g6443** | 68.134 | 1.400 | 0.315 | 4.448 | 8.67E-06 | 0.001116202 | 120.7 kDa protein in NOF-FB transposable element |
| **g43738** | 7481.732 | 1.122 | 0.253 | 4.429 | 9.47E-06 | 0.001210819 | PREDICTED: uncharacterized protein LOC105448412 isoform X2 |
| **g7018** | 43.712 | 1.650 | 0.373 | 4.422 | 9.77E-06 | 0.001239327 | Sodium/potassium-transporting ATPase subunit alpha-like |
| **g3242** | 5668.028 | 0.643 | 0.147 | 4.359 | 1.31E-05 | 0.001648655 | Esterase FE4 |
| **g9642** | 846.158 | -1.508 | 0.346 | -4.353 | 1.34E-05 | 0.001675468 | Putative defense protein 3 |
| **g991** | 594.132 | -0.581 | 0.134 | -4.349 | 1.36E-05 | 0.001693045 | Laminin subunit beta-1 isoform X1 |
| **g2861** | 216.980 | -1.185 | 0.273 | -4.345 | 1.39E-05 | 0.001701927 | Espin isoform X1 |
| **g14145** | 126.376 | -6.533 | 1.506 | -4.337 | 1.44E-05 | 0.001753762 | ---NA--- |
| **g4857** | 797.538 | -1.286 | 0.298 | -4.311 | 1.62E-05 | 0.001941085 | Cartilage oligomeric matrix protein |
| **g15365** | 222.823 | -7.454 | 1.737 | -4.291 | 1.78E-05 | 0.002114487 | Cuticle protein 21-like |
| **g11859** | 104.384 | -1.151 | 0.269 | -4.279 | 1.88E-05 | 0.002201793 | Urea transporter 1-like |
| **g14890** | 267.069 | 0.838 | 0.196 | 4.273 | 1.93E-05 | 0.00222693 | ---NA--- |
| **g12288** | 13.233 | -7.104 | 1.664 | -4.268 | 1.97E-05 | 0.002244935 | ---NA--- |
| **g822** | 211955.707 | -0.973 | 0.228 | -4.265 | 2.00E-05 | 0.00226164 | PREDICTED: uncharacterized protein LOC108756034 |
| **g147** | 4380.185 | -0.809 | 0.190 | -4.260 | 2.04E-05 | 0.00229839 | D-glucuronyl C5-epimerase |
| **g7845** | 244.170 | -0.935 | 0.221 | -4.224 | 2.40E-05 | 0.002661437 | General transcriptional corepressor trfA-like |
| **g5495** | 766.127 | 0.761 | 0.180 | 4.216 | 2.48E-05 | 0.002721502 | Venom acid phosphatase Acph-1-like |
| **g1871** | 443.223 | 0.653 | 0.155 | 4.211 | 2.54E-05 | 0.0027624 | Glucosylceramidase-like isoform X2 |
| **g872** | 1056.032 | -0.773 | 0.184 | -4.191 | 2.77E-05 | 0.002999106 | OTOP3 protein |

| | | | | | | | |
|---|---|---|---|---|---|---|---|
| **g17686** | 181.638 | -4.914 | 1.177 | -4.177 | 2.96E-05 | 0.003160046 | Cuticle protein 12.5 |
| **g2091** | 179.737 | -1.084 | 0.261 | -4.160 | 3.18E-05 | 0.003378115 | Chaoptin |
| **g4800** | 190.316 | -0.838 | 0.203 | -4.134 | 3.57E-05 | 0.003764653 | ---NA--- |
| **g7404** | 69.481 | -1.481 | 0.358 | -4.131 | 3.61E-05 | 0.003779254 | PREDICTED: uncharacterized protein LOC105452472 |
| **g11905** | 117.546 | 1.316 | 0.319 | 4.128 | 3.66E-05 | 0.003814119 | PREDICTED: uncharacterized protein LOC105459363 |
| **g8508** | 224.613 | -0.896 | 0.218 | -4.122 | 3.76E-05 | 0.003896216 | PREDICTED: uncharacterized protein LOC105462636 |
| **g41038** | 13.703 | -4.280 | 1.040 | -4.114 | 3.88E-05 | 0.003992044 | ---NA--- |
| **g15582** | 1575.261 | 1.171 | 0.285 | 4.110 | 3.96E-05 | 0.004052973 | PREDICTED: uncharacterized protein LOC105451837 |
| **g11957** | 573.067 | -1.098 | 0.268 | -4.104 | 4.05E-05 | 0.004119804 | Cytochrome P450 4C1-like |
| **g10896** | 14.394 | -3.066 | 0.752 | -4.075 | 4.60E-05 | 0.004634344 | Platelet glycoprotein Ib alpha chain |
| **g99** | 197.535 | -0.858 | 0.211 | -4.071 | 4.68E-05 | 0.00464588 | PREDICTED: uncharacterized protein LOC105450426 isoform X4 |
| **g13103** | 17.723 | -3.563 | 0.875 | -4.071 | 4.68E-05 | 0.00464588 | ---NA--- |
| **g28540** | 18.430 | -5.759 | 1.419 | -4.058 | 4.95E-05 | 0.004881622 | ---NA--- |
| **g8225** | 187.164 | 0.907 | 0.223 | 4.056 | 4.99E-05 | 0.004892035 | PREDICTED: uncharacterized protein LOC105561312 isoform X2 |
| **g9192** | 336.780 | 1.044 | 0.258 | 4.040 | 5.36E-05 | 0.005223689 | CCCP protein |
| **g8057** | 7907.965 | 1.147 | 0.285 | 4.025 | 5.69E-05 | 0.005488126 | Reverse transcriptase |
| **g11340** | 658.089 | -0.752 | 0.187 | -4.016 | 5.92E-05 | 0.005676733 | Mariner Mos1 transposase |
| **g12493** | 927.317 | 0.998 | 0.250 | 3.998 | 6.39E-05 | 0.00608807 | Ejaculatory bulb-specific protein 3-like |
| **g19019** | 135.412 | -4.025 | 1.013 | -3.975 | 7.03E-05 | 0.006585286 | ---NA--- |
| **g16627** | 454.161 | -1.100 | 0.278 | -3.959 | 7.52E-05 | 0.007008671 | RuvB-like 2 |
| **g10712** | 645.033 | -0.603 | 0.152 | -3.955 | 7.66E-05 | 0.00710121 | Chondroitin proteoglycan-2 |
| **g7369** | 664.919 | 0.713 | 0.181 | 3.952 | 7.76E-05 | 0.007152422 | Clavesin-2-like |
| **g18824** | 20.986 | 4.327 | 1.097 | 3.945 | 7.97E-05 | 0.007311719 | ---NA--- |
| **g5489** | 1623.031 | 0.721 | 0.184 | 3.928 | 8.58E-05 | 0.007738992 | Cytochrome P450 4g15 |
| **g618** | 294.550 | -0.710 | 0.181 | -3.920 | 8.86E-05 | 0.007906913 | B-cell lymphoma/leukemia 11A |
| **g2197** | 611.917 | -0.601 | 0.154 | -3.901 | 9.59E-05 | 0.008381552 | Regulating synaptic membrane exocytosis protein 2 isoform X1 |
| **g9745** | 45.024 | -1.895 | 0.488 | -3.882 | 0.000103464 | 0.008943284 | Fatty acyl-CoA reductase wat isoform X3 |

| | | | | | | | |
|---|---|---|---|---|---|---|---|
| **g295** | 446.954 | -0.680 | 0.175 | -3.873 | 0.000107 397 | 0.009186284 | A disintegrin and metalloproteinase with thrombospondin motifs 9 isoform X1 |
| **g1567** | 1025.787 | -0.905 | 0.234 | -3.872 | 0.000107 805 | 0.009186284 | Integrin beta-PS isoform X1 |
| **g157** | 6943.849 | 0.894 | 0.231 | 3.868 | 0.000109 65 | 0.009276752 | Dual oxidase 2-like |
| **g12236** | 15.735 | -3.845 | 0.994 | -3.867 | 0.000109 966 | 0.009276752 | Cuticle protein 8 |
| **g8467** | 939.008 | -0.966 | 0.250 | -3.860 | 0.000113 595 | 0.009535234 | Cytochrome P450 4C1-like isoform X1 |
| **g110** | 861.557 | -0.948 | 0.246 | -3.845 | 0.000120 593 | 0.009989911 | PREDICTED: uncharacterized protein LOC105449102 |
| **g7515** | 1687.168 | -0.792 | 0.206 | -3.843 | 0.000121 38 | 0.009989911 | D-beta-hydroxybutyrate dehydrogenase, mitochondrial |

**Table S11.** Differentially expressed genes and genes with differential transcript usage across comparisons. "HvsL" indicates the high- vs. low-activity bodyguard comparison. "BvsL" indicates the low-activity bodyguard vs. broodcare worker comparison. "BvsH" indicates the high-activity bodyguard vs. broodcare worker comparison.

| Gene | HvsL | BvsH | BvsL | Analysis | Comparison | Up-regulation |
|---|---|---|---|---|---|---|
| **g3245** | 1 | 0 | 1 | DGE (HvsL) | HvsL + BvsL | Broodcare workers and High-activity Bodyguards |
| **g7229** | 1 | 0 | 1 | DGE (HvsL) + DTU (HvsL) | HvsL + BvsL | Low-activity Bodyguards |
| **g15** | 1 | 0 | 1 | DTU (HvsL) | HvsL + BvsL | NA |
| **g3338** | 1 | 0 | 1 | DTU (HvsL) | HvsL + BvsL | NA |
| **g10981** | 0 | 1 | 1 | DGE | BvsH + BvsL | High-activity + Low-activity Bodyguards |
| **g1423** | 0 | 1 | 1 | DGE | BvsH + BvsL | High-activity + Low-activity Bodyguards |
| **g15134** | 0 | 1 | 1 | DGE | BvsH + BvsL | High-activity + Low-activity Bodyguards |
| **g24224** | 0 | 1 | 1 | DGE | BvsH + BvsL | High-activity + Low-activity Bodyguards |
| **g157** | 0 | 1 | 1 | DGE | BvsH + BvsL | High-activity + Low-activity Bodyguards |
| **g20** | 0 | 1 | 1 | DTU | BvsH + BvsL | NA |
| **g41100** | 0 | 1 | 1 | DGE | BvsH + BvsL | High-activity + Low-activity Bodyguards |
| **g48923** | 0 | 1 | 1 | DGE | BvsH + BvsL | High-activity + Low-activity Bodyguards |
| **g6874** | 0 | 1 | 1 | DGE | BvsH + BvsL | High-activity + Low-activity Bodyguards |
| **g6897** | 0 | 1 | 1 | DGE | BvsH + BvsL | High-activity + Low-activity Bodyguards |
| **g8463** | 0 | 1 | 1 | DGE | BvsH + BvsL | High-activity + Low-activity Bodyguards |
| **g9011** | 0 | 1 | 1 | DGE | BvsH + BvsL | High-activity + Low-activity Bodyguards |
| **g9196** | 0 | 1 | 1 | DGE | BvsH + BvsL | High-activity + Low-activity Bodyguards |
| **g12** | 0 | 1 | 1 | DTU | BvsH + BvsL | NA |
| **g1871** | 0 | 1 | 1 | DGE | BvsH + BvsL | High-activity + Low-activity Bodyguards |
| **g10980** | 0 | 1 | 1 | DGE | BvsH + BvsL | High-activity + Low-activity Bodyguards |
| **g16054** | 0 | 1 | 1 | DGE | BvsH + BvsL | High-activity + Low-activity Bodyguards |
| **g1** | 0 | 1 | 1 | DGE | BvsH + BvsL | Broodcare workers |
| **g10119** | 0 | 1 | 1 | DGE | BvsH + BvsL | Broodcare workers |

| | | | | | | |
|---|---|---|---|---|---|---|
| **g1077** | 0 | 1 | 1 | DGE | BvsH + BvsL | Broodcare workers |
| **g11369** | 0 | 1 | 1 | DGE | BvsH + BvsL | Broodcare workers |
| **g1156** | 0 | 1 | 1 | DGE | BvsH + BvsL | Broodcare workers |
| **g11760** | 0 | 1 | 1 | DGE | BvsH + BvsL | Broodcare workers |
| **g11825** | 0 | 1 | 1 | DGE | BvsH + BvsL | Broodcare workers |
| **g12056** | 0 | 1 | 1 | DGE | BvsH + BvsL | Broodcare workers |
| **g13057** | 0 | 1 | 1 | DGE | BvsH + BvsL | Broodcare workers |
| **g13235** | 0 | 1 | 1 | DGE | BvsH + BvsL | Broodcare workers |
| **g13788** | 0 | 1 | 1 | DGE | BvsH + BvsL | Broodcare workers |
| **g1382** | 0 | 1 | 1 | DGE | BvsH + BvsL | Broodcare workers |
| **g1495** | 0 | 1 | 1 | DGE | BvsH + BvsL | Broodcare workers |
| **g15701** | 0 | 1 | 1 | DGE | BvsH + BvsL | Broodcare workers |
| **g15993** | 0 | 1 | 1 | DGE | BvsH + BvsL | Broodcare workers |
| **g16911** | 0 | 1 | 1 | DGE | BvsH + BvsL | Broodcare workers |
| **g1707** | 0 | 1 | 1 | DGE | BvsH + BvsL | Broodcare workers |
| **g1110*** | 0 | 1 | 1 | DTU | BvsH + BvsL | NA |
| **g1150*** | 0 | 1 | 1 | DTU | BvsH + BvsL | NA |
| **g23784** | 0 | 1 | 1 | DGE | BvsH + BvsL | Broodcare workers |
| **g1253*** | 0 | 1 | 1 | DTU | BvsH + BvsL | NA |
| **g1435*** | 0 | 1 | 1 | DTU | BvsH + BvsL | NA |
| **g26067** | 0 | 1 | 1 | DGE | BvsH + BvsL | Broodcare workers |
| **g2622** | 0 | 1 | 1 | DGE | BvsH + BvsL | Broodcare workers |
| **g2759** | 0 | 1 | 1 | DGE | BvsH + BvsL | Broodcare workers |
| **g1672** | 0 | 1 | 1 | DTU | BvsH + BvsL | NA |
| **g2926** | 0 | 1 | 1 | DGE | BvsH + BvsL | Broodcare workers |
| **g3043** | 0 | 1 | 1 | DGE + DTU | BvsH + BvsL | Broodcare workers |
| **g19019** | 0 | 1 | 1 | DGE | BvsH + BvsL | Broodcare workers |

| | | | | | | |
|---|---|---|---|---|---|---|
| **g32** | 0 | 1 | 1 | DGE | BvsH + BvsL | Broodcare workers |
| **g3224** | 0 | 1 | 1 | DGE | BvsH + BvsL | Broodcare workers |
| **g329** | 0 | 1 | 1 | DGE | BvsH + BvsL | Broodcare workers |
| **g2039** | 0 | 1 | 1 | DTU | BvsH + BvsL | NA |
| **g2176*** | 0 | 1 | 1 | DTU | BvsH + BvsL | NA |
| **g3740** | 0 | 1 | 1 | DGE | BvsH + BvsL | Broodcare workers |
| **g2348*** | 0 | 1 | 1 | DTU | BvsH + BvsL | NA |
| **g3892** | 0 | 1 | 1 | DGE | BvsH + BvsL | Broodcare workers |
| **g3958** | 0 | 1 | 1 | DGE | BvsH + BvsL | Broodcare workers |
| **g3989** | 0 | 1 | 1 | DGE | BvsH + BvsL | Broodcare workers |
| **g4166** | 0 | 1 | 1 | DGE | BvsH + BvsL | Broodcare workers |
| **g4305** | 0 | 1 | 1 | DGE | BvsH + BvsL | Broodcare workers |
| **g4614** | 0 | 1 | 1 | DGE | BvsH + BvsL | Broodcare workers |
| **g4715** | 0 | 1 | 1 | DGE | BvsH + BvsL | Broodcare workers |
| **g482** | 0 | 1 | 1 | DGE | BvsH + BvsL | Broodcare workers |
| **g4947** | 0 | 1 | 1 | DGE | BvsH + BvsL | Broodcare workers |
| **g4967** | 0 | 1 | 1 | DGE | BvsH + BvsL | Broodcare workers |
| **g4981** | 0 | 1 | 1 | DGE | BvsH + BvsL | Broodcare workers |
| **g3623** | 0 | 1 | 1 | DTU | BvsH + BvsL | NA |
| **g5108** | 0 | 1 | 1 | DGE | BvsH + BvsL | Broodcare workers |
| **g5273** | 0 | 1 | 1 | DGE | BvsH + BvsL | Broodcare workers |
| **g5301** | 0 | 1 | 1 | DGE | BvsH + BvsL | Broodcare workers |
| **g5480** | 0 | 1 | 1 | DGE | BvsH + BvsL | Broodcare workers |
| **g5483** | 0 | 1 | 1 | DGE | BvsH + BvsL | Broodcare workers |
| **g562** | 0 | 1 | 1 | DGE | BvsH + BvsL | Broodcare workers |
| **g5675** | 0 | 1 | 1 | DGE | BvsH + BvsL | Broodcare workers |
| **g6509** | 0 | 1 | 1 | DGE | BvsH + BvsL | Broodcare workers |

| | | | | | | |
|---|---|---|---|---|---|---|
| **g658** | 0 | 1 | 1 | DGE | BvsH + BvsL | Broodcare workers |
| **g683** | 0 | 1 | 1 | DGE | BvsH + BvsL | Broodcare workers |
| **g7529** | 0 | 1 | 1 | DGE | BvsH + BvsL | Broodcare workers |
| **g13103** | 0 | 1 | 1 | DGE | BvsH + BvsL | Broodcare workers |
| **g829** | 0 | 1 | 1 | DGE | BvsH + BvsL | Broodcare workers |
| **g8459** | 0 | 1 | 1 | DGE | BvsH + BvsL | Broodcare workers |
| **g8464** | 0 | 1 | 1 | DGE + DTU | BvsH + BvsL | Broodcare workers |
| **g868** | 0 | 1 | 1 | DGE | BvsH + BvsL | Broodcare workers |
| **g869** | 0 | 1 | 1 | DGE + DTU | BvsH + BvsL | Broodcare workers |
| **g8467** | 0 | 1 | 1 | DGE | BvsH + BvsL | Broodcare workers |
| **g960*** | 0 | 1 | 1 | DTU | BvsH + BvsL | NA |
| **g9809** | 0 | 1 | 1 | DGE | BvsH + BvsL | Broodcare workers |
| **g9996** | 0 | 1 | 1 | DGE | BvsH + BvsL | Broodcare workers |
| **g295** | 0 | 1 | 1 | DGE | BvsH + BvsL | Broodcare workers |
| **g1818** | 0 | 1 | 1 | DGE | BvsH + BvsL | Broodcare workers |
| **g643** | 0 | 1 | 1 | DGE | BvsH + BvsL | Broodcare workers |
| **g5767** | 0 | 1 | 1 | DGE | BvsH + BvsL | Broodcare workers |
| **g6169** | 0 | 1 | 1 | DGE | BvsH + BvsL | Broodcare workers |
| **g2658*** | 0 | 1 | 1 | DTU | BvsH + BvsL | NA |
| **g13430** | 0 | 1 | 1 | DGE | BvsH + BvsL | Broodcare workers |
| **g17024** | 0 | 1 | 1 | DGE | BvsH + BvsL | Broodcare workers |
| **g60** | 0 | 1 | 1 | DTU | BvsH + BvsL | NA |
| **g10913** | 0 | 1 | 1 | DGE | BvsH + BvsL | Broodcare workers |
| **g1172** | 0 | 1 | 1 | DGE | BvsH + BvsL | Broodcare workers |
| **g2745** | 0 | 1 | 1 | DGE | BvsH + BvsL | Broodcare workers |
| **g2940** | 0 | 1 | 1 | DGE + DTU | BvsH + BvsL | Broodcare workers |
| **g508** | 0 | 1 | 1 | DGE + DTU | BvsH + BvsL | Broodcare workers |

| | | | | | | |
|---|---|---|---|---|---|---|
| **g8242** | 0 | 1 | 1 | DGE | BvsH + BvsL | Broodcare workers |
| **g8960** | 0 | 1 | 1 | DGE | BvsH + BvsL | Broodcare workers |
| **g13589** | 1 | 1 | 1 | DGE (HvsL) | HvsL + BvsH + BvsL | Broodcare workers and High-activity Bodyguards |
| **g13731** | 1 | 1 | 1 | DGE (HvsL) | HvsL + BvsH + BvsL | Broodcare workers and High-activity Bodyguards |
| **g13743** | 1 | 1 | 1 | DGE (HvsL) | HvsL + BvsH + BvsL | Broodcare workers and High-activity Bodyguards |
| **g20498** | 1 | 1 | 1 | DGE (HvsL) | HvsL + BvsH + BvsL | Broodcare workers and High-activity Bodyguards |
| **g22293** | 1 | 1 | 1 | DGE (HvsL) | HvsL + BvsH + BvsL | Broodcare workers and High-activity Bodyguards |
| **g5151** | 1 | 1 | 1 | DGE (HvsL) | HvsL + BvsH + BvsL | Broodcare workers and High-activity Bodyguards |
| **g5823** | 1 | 1 | 1 | DGE (HvsL) + DTU (HvsL) | HvsL + BvsH + BvsL | Broodcare workers and High-activity Bodyguards |
| **g6591** | 1 | 1 | 1 | DGE (HvsL) | HvsL + BvsH + BvsL | Broodcare workers and Low-activity Bodyguards |
| **g6663** | 1 | 1 | 1 | DGE (HvsL) + DTU | HvsL + BvsH + BvsL | Broodcare workers and High-activity Bodyguards |
| **g5024*** | 1 | 1 | 1 | DTU (HvsL) | HvsL + BvsH + BvsL | NA |
| **g7216** | 1 | 1 | 1 | DGE (HvsL) | HvsL + BvsH + BvsL | Broodcare workers and High-activity Bodyguards |
| **g8064** | 1 | 1 | 1 | DGE (HvsL) | HvsL + BvsH + BvsL | Broodcare workers and High-activity Bodyguards |
| **g8173** | 1 | 1 | 1 | DGE (HvsL) | HvsL + BvsH + BvsL | Broodcare workers and High-activity Bodyguards |
| **g9305** | 1 | 1 | 1 | DGE (HvsL) | HvsL + BvsH + BvsL | Broodcare workers and High-activity Bodyguards |
| **g9792** | 1 | 1 | 1 | DGE (HvsL) | HvsL + BvsH + BvsL | Broodcare workers and High-activity Bodyguards |
| **g5389** | 1 | 1 | 1 | DGE (HvsL) | HvsL + BvsH + BvsL | Broodcare workers and High-activity Bodyguards |
| **g7219** | 1 | 1 | 1 | DGE (HvsL) | HvsL + BvsH + BvsL | Broodcare workers and High-activity Bodyguards |
| **g5511** | 1 | 1 | 1 | DGE (HvsL) + DTU (HvsL) | HvsL + BvsH + BvsL | Broodcare workers and High-activity Bodyguards |
| **g123** | 1 | 1 | 1 | DGE (HvsL) | HvsL + BvsH + BvsL | Broodcare workers and High-activity Bodyguards |
| **g12874** | 1 | 1 | 0 | DGE (HvsL) | HvsL + BvsH | Broodcare workers and Low-activity Bodyguards |
| **g17038** | 1 | 1 | 0 | DGE (HvsL) | HvsL + BvsH | Broodcare workers and Low-activity Bodyguards |
| **g6602** | 1 | 1 | 0 | DGE (HvsL) | HvsL + BvsH | Broodcare workers and Low-activity Bodyguards |
| **g8308** | 1 | 1 | 0 | DGE (HvsL) | HvsL + BvsH | Broodcare workers and Low-activity Bodyguards |
| **g1894** | 1 | 0 | 0 | DGE (HvsL) | HvsL | High-activity Bodyguards |
| **g2855** | 1 | 0 | 0 | DGE (HvsL) | HvsL | High-activity Bodyguards |

| | | | | | | |
|---|---|---|---|---|---|---|
| **g5994** | 1 | 0 | 0 | DGE (HvsL) | HvsL | High-activity Bodyguards |
| **g3684** | 1 | 0 | 0 | DGE (HvsL) | HvsL | High-activity Bodyguards |
| **g2275** | 1 | 0 | 0 | DGE (HvsL) | HvsL | High-activity Bodyguards |
| **g3814** | 1 | 0 | 0 | DGE (HvsL) | HvsL | High-activity Bodyguards |
| **g20928** | 1 | 0 | 0 | DGE (HvsL) | HvsL | High-activity Bodyguards |
| **g6982** | 1 | 0 | 0 | DGE (HvsL) | HvsL | Low-activity Bodyguards |
| **g9141** | 1 | 0 | 0 | DGE (HvsL) | HvsL | Low-activity Bodyguards |
| **g24155** | 1 | 0 | 0 | DGE (HvsL) | HvsL | Low-activity Bodyguards |
| **g16177** | 1 | 0 | 0 | DGE (HvsL) | HvsL | Low-activity Bodyguards |
| **g6884** | 1 | 0 | 0 | DGE (HvsL) | HvsL | Low-activity Bodyguards |
| **g12782** | 1 | 0 | 0 | DGE (HvsL) | HvsL | Low-activity Bodyguards |
| **g6082** | 1 | 0 | 0 | DGE (HvsL) | HvsL | Low-activity Bodyguards |
| **g11332** | 1 | 0 | 0 | DGE (HvsL) | HvsL | Low-activity Bodyguards |
| **g6124** | 1 | 0 | 0 | DGE (HvsL) | HvsL | Low-activity Bodyguards |
| **g16223** | 1 | 0 | 0 | DGE (HvsL) | HvsL | Low-activity Bodyguards |
| **g4350*** | 1 | 0 | 0 | DTU (HvsL) | HvsL | NA |
| **g4830*** | 1 | 0 | 0 | DTU (HvsL) | HvsL | NA |
| **g4886*** | 1 | 0 | 0 | DTU (HvsL) | HvsL | NA |
| **g5260*** | 1 | 0 | 0 | DTU (HvsL) | HvsL | NA |
| **g5456*** | 1 | 0 | 0 | DTU (HvsL) | HvsL | NA |
| **g8013*** | 1 | 0 | 0 | DTU (HvsL) | HvsL | NA |
| **g8428*** | 1 | 0 | 0 | DTU (HvsL) | HvsL | NA |
| **g1280** | 0 | 1 | 0 | DGE | BvsH | High-activity Bodyguards |
| **g164** | 0 | 1 | 0 | DGE | BvsH | High-activity Bodyguards |
| **g26785** | 0 | 1 | 0 | DGE | BvsH | High-activity Bodyguards |
| **g29392** | 0 | 1 | 0 | DGE | BvsH | High-activity Bodyguards |
| **g1900** | 0 | 1 | 0 | DTU | BvsH | NA |

| | | | | | | |
|---|---|---|---|---|---|---|
| **g413** | 0 | 1 | 0 | DGE | BvsH | High-activity Bodyguards |
| **g3574** | 0 | 1 | 0 | DTU | BvsH | NA |
| **g5191** | 0 | 1 | 0 | DGE | BvsH | High-activity Bodyguards |
| **g6531** | 0 | 1 | 0 | DGE | BvsH | High-activity Bodyguards |
| **g6067** | 0 | 1 | 0 | DTU | BvsH | NA |
| **g8438** | 0 | 1 | 0 | DGE | BvsH | High-activity Bodyguards |
| **g10275** | 0 | 1 | 0 | DGE | BvsH | Broodcare workers |
| **g1048** | 0 | 1 | 0 | DGE | BvsH | Broodcare workers |
| **g10859** | 0 | 1 | 0 | DGE | BvsH | Broodcare workers |
| **g12996** | 0 | 1 | 0 | DGE | BvsH | Broodcare workers |
| **g1381** | 0 | 1 | 0 | DGE | BvsH | Broodcare workers |
| **g14080** | 0 | 1 | 0 | DGE | BvsH | Broodcare workers |
| **g14253** | 0 | 1 | 0 | DGE | BvsH | Broodcare workers |
| **g15782** | 0 | 1 | 0 | DGE | BvsH | Broodcare workers |
| **g15833** | 0 | 1 | 0 | DGE | BvsH | Broodcare workers |
| **g17163** | 0 | 1 | 0 | DGE | BvsH | Broodcare workers |
| **g17637** | 0 | 1 | 0 | DGE | BvsH | Broodcare workers |
| **g17942** | 0 | 1 | 0 | DGE | BvsH | Broodcare workers |
| **g20799** | 0 | 1 | 0 | DGE | BvsH | Broodcare workers |
| **g20880** | 0 | 1 | 0 | DGE | BvsH | Broodcare workers |
| **g21793** | 0 | 1 | 0 | DGE | BvsH | Broodcare workers |
| **g2226** | 0 | 1 | 0 | DGE | BvsH | Broodcare workers |
| **g2273** | 0 | 1 | 0 | DGE | BvsH | Broodcare workers |
| **g25699** | 0 | 1 | 0 | DGE | BvsH | Broodcare workers |
| **g25995** | 0 | 1 | 0 | DGE | BvsH | Broodcare workers |
| **g26577** | 0 | 1 | 0 | DGE | BvsH | Broodcare workers |
| **g26724** | 0 | 1 | 0 | DGE | BvsH | Broodcare workers |

| | | | | | | |
|---|---|---|---|---|---|---|
| **g27103** | 0 | 1 | 0 | DGE | BvsH | Broodcare workers |
| **g2732** | 0 | 1 | 0 | DGE | BvsH | Broodcare workers |
| **g33510** | 0 | 1 | 0 | DGE | BvsH | Broodcare workers |
| **g3978** | 0 | 1 | 0 | DGE | BvsH | Broodcare workers |
| **g4234** | 0 | 1 | 0 | DGE | BvsH | Broodcare workers |
| **g5065** | 0 | 1 | 0 | DGE | BvsH | Broodcare workers |
| **g5073** | 0 | 1 | 0 | DGE | BvsH | Broodcare workers |
| **g52588** | 0 | 1 | 0 | DGE | BvsH | Broodcare workers |
| **g6490** | 0 | 1 | 0 | DGE | BvsH | Broodcare workers |
| **g656** | 0 | 1 | 0 | DGE | BvsH | Broodcare workers |
| **g66062** | 0 | 1 | 0 | DGE | BvsH | Broodcare workers |
| **g6715** | 0 | 1 | 0 | DGE | BvsH | Broodcare workers |
| **g6910** | 0 | 1 | 0 | DGE | BvsH | Broodcare workers |
| **g6983** | 0 | 1 | 0 | DGE | BvsH | Broodcare workers |
| **g7620** | 0 | 1 | 0 | DGE | BvsH | Broodcare workers |
| **g7906** | 0 | 1 | 0 | DGE | BvsH | Broodcare workers |
| **g8183** | 0 | 1 | 0 | DGE | BvsH | Broodcare workers |
| **g8245** | 0 | 1 | 0 | DGE | BvsH | Broodcare workers |
| **g8422** | 0 | 1 | 0 | DGE | BvsH | Broodcare workers |
| **g8120** | 0 | 1 | 0 | DTU | BvsH | NA |
| **g9315** | 0 | 1 | 0 | DGE | BvsH | Broodcare workers |
| **g11844** | 0 | 1 | 0 | DGE | BvsH | Broodcare workers |
| **g14127** | 0 | 1 | 0 | DGE | BvsH | High-activity Bodyguards |
| **g14873** | 0 | 1 | 0 | DGE | BvsH | High-activity Bodyguards |
| **g1838** | 0 | 1 | 0 | DGE | BvsH | Broodcare workers |
| **g18584** | 0 | 1 | 0 | DGE | BvsH | Broodcare workers |
| **g23139** | 0 | 1 | 0 | DGE | BvsH | High-activity Bodyguards |

| | | | | | | |
|---|---|---|---|---|---|---|
| **g26621** | 0 | 1 | 0 | DGE | BvsH | Broodcare workers |
| **g27808** | 0 | 1 | 0 | DGE | BvsH | Broodcare workers |
| **g31059** | 0 | 1 | 0 | DGE | BvsH | Broodcare workers |
| **g34531** | 0 | 1 | 0 | DGE | BvsH | Broodcare workers |
| **g38173** | 0 | 1 | 0 | DGE | BvsH | Broodcare workers |
| **g6037** | 0 | 1 | 0 | DGE | BvsH | Broodcare workers |
| **g8108** | 0 | 1 | 0 | DGE | BvsH | Broodcare workers |
| **g8148** | 0 | 1 | 0 | DGE | BvsH | Broodcare workers |
| **g245** | 0 | 1 | 0 | DTU | BvsH | NA |
| **g1669** | 0 | 1 | 0 | DTU | BvsH | NA |
| **g1692** | 0 | 1 | 0 | DTU | BvsH | NA |
| **g2640** | 0 | 1 | 0 | DTU | BvsH | NA |
| **g3204** | 0 | 1 | 0 | DTU | BvsH | NA |
| **g8364** | 0 | 1 | 0 | DTU | BvsH | NA |
| **g8495** | 0 | 1 | 0 | DTU | BvsH | NA |
| **g8727** | 0 | 1 | 0 | DTU | BvsH | NA |
| **g12754** | 0 | 1 | 0 | DTU | BvsH | NA |
| **g20031** | 0 | 1 | 0 | DTU | BvsH | NA |
| **g35676** | 0 | 1 | 0 | DTU | BvsH | NA |
| **g110** | 0 | 0 | 1 | DGE | BvsL | Broodcare workers |
| **g18140** | 0 | 0 | 1 | DGE | BvsL | Broodcare workers |
| **g25125** | 0 | 0 | 1 | DGE | BvsL | Broodcare workers |
| **g35603** | 0 | 0 | 1 | DGE | BvsL | Broodcare workers |
| **g8104** | 0 | 0 | 1 | DGE | BvsL | Broodcare workers |
| **g11859** | 0 | 0 | 1 | DGE | BvsL | Broodcare workers |
| **g17686** | 0 | 0 | 1 | DGE | BvsL | Broodcare workers |
| **g872** | 0 | 0 | 1 | DGE | BvsL | Broodcare workers |

| | | | | | | |
|---|---|---|---|---|---|---|
| **g3461** | 0 | 0 | 1 | DGE | BvsL | Broodcare workers |
| **g4815** | 0 | 0 | 1 | DGE | BvsL | Broodcare workers |
| **g822** | 0 | 0 | 1 | DGE + DTU | BvsL | Broodcare workers |
| **g7845** | 0 | 0 | 1 | DGE | BvsL | Broodcare workers |
| **g8508** | 0 | 0 | 1 | DGE | BvsL | Broodcare workers |
| **g1388** | 0 | 0 | 1 | DTU | BvsL | NA |
| **g2002** | 0 | 0 | 1 | DTU | BvsL | NA |
| **g12493** | 0 | 0 | 1 | DGE | BvsL | Low-activity Bodyguards |
| **g11905** | 0 | 0 | 1 | DGE | BvsL | Low-activity Bodyguards |
| **g7743** | 0 | 0 | 1 | DGE | BvsL | Low-activity Bodyguards |
| **g15582** | 0 | 0 | 1 | DGE | BvsL | Low-activity Bodyguards |
| **g1494** | 0 | 0 | 1 | DGE | BvsL | Low-activity Bodyguards |
| **g2598** | 0 | 0 | 1 | DGE | BvsL | Low-activity Bodyguards |
| **g3023** | 0 | 0 | 1 | DGE | BvsL | Low-activity Bodyguards |
| **g6443** | 0 | 0 | 1 | DGE | BvsL | Low-activity Bodyguards |
| **g19862** | 0 | 0 | 1 | DGE | BvsL | Low-activity Bodyguards |
| **g30386** | 0 | 0 | 1 | DGE | BvsL | Low-activity Bodyguards |
| **g35380** | 0 | 0 | 1 | DGE | BvsL | Low-activity Bodyguards |
| **g5495** | 0 | 0 | 1 | DGE | BvsL | Low-activity Bodyguards |
| **g8320** | 0 | 0 | 1 | DGE | BvsL | Low-activity Bodyguards |
| **g8265** | 0 | 0 | 1 | DGE | BvsL | Low-activity Bodyguards |
| **g9192** | 0 | 0 | 1 | DGE | BvsL | Low-activity Bodyguards |
| **g11340** | 0 | 0 | 1 | DGE | BvsL | Broodcare workers |
| **g1567** | 0 | 0 | 1 | DGE | BvsL | Broodcare workers |
| **g9272** | 0 | 0 | 1 | DGE | BvsL | Broodcare workers |
| **g1211** | 0 | 0 | 1 | DTU | BvsL | NA |
| **g3049** | 0 | 0 | 1 | DTU | BvsL | NA |

| | | | | | | |
|---|---|---|---|---|---|---|
| **g12341** | 0 | 0 | 1 | DGE | BvsL | Broodcare workers |
| **g14145** | 0 | 0 | 1 | DGE | BvsL | Broodcare workers |
| **g20826** | 0 | 0 | 1 | DGE | BvsL | Broodcare workers |
| **g2197** | 0 | 0 | 1 | DGE | BvsL | Broodcare workers |
| **g4674** | 0 | 0 | 1 | DGE | BvsL | Broodcare workers |
| **g4857** | 0 | 0 | 1 | DGE | BvsL | Broodcare workers |
| **g1078** | 0 | 0 | 1 | DGE | BvsL | Broodcare workers |
| **g1142** | 0 | 0 | 1 | DGE | BvsL | Broodcare workers |
| **g1286** | 0 | 0 | 1 | DGE | BvsL | Broodcare workers |
| **g147** | 0 | 0 | 1 | DGE | BvsL | Broodcare workers |
| **g2091** | 0 | 0 | 1 | DGE | BvsL | Broodcare workers |
| **g2861** | 0 | 0 | 1 | DGE | BvsL | Broodcare workers |
| **g4800** | 0 | 0 | 1 | DGE | BvsL | Broodcare workers |
| **g5791** | 0 | 0 | 1 | DGE | BvsL | Broodcare workers |
| **g7255** | 0 | 0 | 1 | DGE | BvsL | Broodcare workers |
| **g768** | 0 | 0 | 1 | DGE | BvsL | Broodcare workers |
| **g10712** | 0 | 0 | 1 | DGE | BvsL | Broodcare workers |
| **g11957** | 0 | 0 | 1 | DGE | BvsL | Broodcare workers |
| **g12963** | 0 | 0 | 1 | DGE | BvsL | Broodcare workers |
| **g13531** | 0 | 0 | 1 | DGE | BvsL | Broodcare workers |
| **g14156** | 0 | 0 | 1 | DGE | BvsL | Broodcare workers |
| **g16627** | 0 | 0 | 1 | DGE | BvsL | Broodcare workers |
| **g18527** | 0 | 0 | 1 | DGE | BvsL | Broodcare workers |
| **g23003** | 0 | 0 | 1 | DGE | BvsL | Broodcare workers |
| **g37213** | 0 | 0 | 1 | DGE | BvsL | Broodcare workers |
| **g41038** | 0 | 0 | 1 | DGE | BvsL | Broodcare workers |
| **g7372** | 0 | 0 | 1 | DGE | BvsL | Broodcare workers |

| | | | | | | |
|---|---|---|---|---|---|---|
| **g7515** | 0 | 0 | 1 | DGE | BvsL | Broodcare workers |
| **g8125** | 0 | 0 | 1 | DGE | BvsL | Broodcare workers |
| **g9189** | 0 | 0 | 1 | DGE | BvsL | Broodcare workers |
| **g9742** | 0 | 0 | 1 | DGE | BvsL | Broodcare workers |
| **g9642** | 0 | 0 | 1 | DGE | BvsL | Broodcare workers |
| **g2274** | 0 | 0 | 1 | DTU | BvsL | NA |
| **g1628** | 0 | 0 | 1 | DGE | BvsL | Low-activity Bodyguards |
| **g5161** | 0 | 0 | 1 | DGE | BvsL | Low-activity Bodyguards |
| **g5489** | 0 | 0 | 1 | DGE | BvsL | Low-activity Bodyguards |
| **g14166** | 0 | 0 | 1 | DGE | BvsL | Low-activity Bodyguards |
| **g28489** | 0 | 0 | 1 | DGE | BvsL | Low-activity Bodyguards |
| **g68084** | 0 | 0 | 1 | DGE | BvsL | Low-activity Bodyguards |
| **g7369** | 0 | 0 | 1 | DGE | BvsL | Low-activity Bodyguards |
| **g14890** | 0 | 0 | 1 | DGE | BvsL | Low-activity Bodyguards |
| **g19974** | 0 | 0 | 1 | DGE | BvsL | Low-activity Bodyguards |
| **g4005** | 0 | 0 | 1 | DGE | BvsL | Low-activity Bodyguards |
| **g3242** | 0 | 0 | 1 | DGE | BvsL | Low-activity Bodyguards |
| **g4107** | 0 | 0 | 1 | DGE | BvsL | Low-activity Bodyguards |
| **g415** | 0 | 0 | 1 | DGE | BvsL | Low-activity Bodyguards |
| **g6937** | 0 | 0 | 1 | DGE | BvsL | Low-activity Bodyguards |
| **g7018** | 0 | 0 | 1 | DGE | BvsL | Low-activity Bodyguards |
| **g1039** | 0 | 0 | 1 | DGE | BvsL | Low-activity Bodyguards |
| **g3171** | 0 | 0 | 1 | DGE | BvsL | Low-activity Bodyguards |
| **g523** | 0 | 0 | 1 | DGE | BvsL | Low-activity Bodyguards |
| **g10104** | 0 | 0 | 1 | DGE | BvsL | Low-activity Bodyguards |
| **g18224** | 0 | 0 | 1 | DGE | BvsL | Low-activity Bodyguards |
| **g23888** | 0 | 0 | 1 | DGE | BvsL | Low-activity Bodyguards |

| | | | | | | |
|---|---|---|---|---|---|---|
| **g43738** | 0 | 0 | 1 | DGE | BvsL | Low-activity Bodyguards |
| **g8057** | 0 | 0 | 1 | DGE | BvsL | Low-activity Bodyguards |
| **g18824** | 0 | 0 | 1 | DGE | BvsL | Low-activity Bodyguards |
| **g8225** | 0 | 0 | 1 | DGE | BvsL | Low-activity Bodyguards |
| **g18601** | 0 | 0 | 1 | DGE | BvsL | Low-activity Bodyguards |
| **g3884** | 0 | 0 | 1 | DGE | BvsL | Broodcare workers |
| **g12288** | 0 | 0 | 1 | DGE | BvsL | Broodcare workers |
| **g10896** | 0 | 0 | 1 | DGE | BvsL | Broodcare workers |
| **g12236** | 0 | 0 | 1 | DGE | BvsL | Broodcare workers |
| **g13218** | 0 | 0 | 1 | DGE | BvsL | Broodcare workers |
| **g28540** | 0 | 0 | 1 | DGE | BvsL | Broodcare workers |
| **g618** | 0 | 0 | 1 | DGE | BvsL | Broodcare workers |
| **g859** | 0 | 0 | 1 | DGE | BvsL | Broodcare workers |
| **g9745** | 0 | 0 | 1 | DGE | BvsL | Broodcare workers |
| **g99** | 0 | 0 | 1 | DGE | BvsL | Broodcare workers |
| **g991** | 0 | 0 | 1 | DGE | BvsL | Broodcare workers |
| **g7404** | 0 | 0 | 1 | DGE | BvsL | Broodcare workers |
| **g15365** | 0 | 0 | 1 | DGE | BvsL | Broodcare workers |
| **g4043** | 0 | 0 | 1 | DTU | BvsL | NA |
| **g17778** | 0 | 0 | 1 | DTU | BvsL | NA |
| **g10445** | 0 | 0 | 1 | DTU | BvsL | NA |
| **g13462** | 0 | 0 | 1 | DTU | BvsL | NA |
| **g2236** | 0 | 0 | 1 | DTU | BvsL | NA |
| **g368** | 0 | 0 | 1 | DTU | BvsL | NA |
| **g9002** | 0 | 0 | 1 | DTU | BvsL | NA |
| **g10518** | 0 | 0 | 1 | DTU | BvsL | NA |
| **g1306** | 0 | 0 | 1 | DTU | BvsL | NA |

| | | | | | | |
|---|---|---|---|---|---|---|
| **g2157** | 0 | 0 | 1 | DTU | BvsL | NA |
| **g2793** | 0 | 0 | 1 | DTU | BvsL | NA |
| **g3589** | 0 | 0 | 1 | DTU | BvsL | NA |
| **g6180** | 0 | 0 | 1 | DTU | BvsL | NA |
| **g6497** | 0 | 0 | 1 | DTU | BvsL | NA |
| **g226** | 0 | 0 | 1 | DTU | BvsL | NA |
| **g2477** | 0 | 0 | 1 | DTU | BvsL | NA |

**Table S12.** Gene Set Enrichment Analysis (GSEA) comparing bodyguards and broodcare workers.

| GO ID | GO.Name | GO type | Size | ES | NES | Nominal p-val | FDR | FWER p-val |
|---|---|---|---|---|---|---|---|---|
| **GO:1901566** | organonitrogen compound biosynthetic process | BP | 23 | -0.451 | -2.245 | 0.000 | 0.0014 | 0.001 |
| **GO:0019637** | organophosphate metabolic process | BP | 18 | -0.470 | -2.080 | 0.000 | 0.0142 | 0.007 |
| **GO:0031090** | organelle membrane | CC | 27 | -0.371 | -2.043 | 0.000 | 0.0138 | 0.008 |
| **GO:0008061** | chitin binding | MF | 18 | 0.711 | 1.772 | 0.000 | 0.0163 | 0.02 |
| **GO:0005198** | structural molecule activity | MF | 25 | 0.665 | 1.688 | 0.001 | 0.0355 | 0.082 |

**Table S13.** RNA-Seq sequence number, sequence length, GC content, and quality scores before and after trimming.

| | Before trimming | | | | After trimming | | | |
|---|---|---|---|---|---|---|---|---|
| Sequence | Total Sequences | Sequence length | %GC | Quality score | Total Sequences | Sequence length | %GC | Quality score |
| 18be_S1_R1_forward | 25024461 | 10-51 | 42 | 36 | 24643733 | 30-51 | 42 | 36 |
| 18be_S1_R2_reverse | 25024461 | 10-51 | 42 | 36 | 24643733 | 30-51 | 41 | 36 |
| 28be_S2_R1_forward | 28584251 | 10-51 | 42 | 36 | 28341356 | 30-51 | 42 | 36 |
| 28be_S2_R2_reverse | 28584251 | 10-51 | 42 | 36 | 28341356 | 30-51 | 42 | 36 |
| 103be_S3_R1_forward | 32165646 | 10-51 | 41 | 36 | 31902391 | 30-51 | 41 | 36 |
| 103be_S3_R2_reverse | 32165646 | 10-51 | 42 | 36 | 31902391 | 30-51 | 41 | 36 |
| 107be_S4_R1_forward | 28290520 | 10-51 | 41 | 36 | 28094068 | 30-51 | 41 | 36 |
| 107be_S4_R2_reverse | 28290520 | 10-51 | 41 | 36 | 28094068 | 30-51 | 41 | 36 |
| 107id_S5_R1_forward | 21601826 | 10-51 | 41 | 36 | 21440112 | 30-51 | 41 | 36 |
| 107id_S5_R2_reverse | 21601826 | 10-51 | 41 | 36 | 21440112 | 30-51 | 41 | 36 |
| 109be_S6_R1_forward | 24984267 | 10-51 | 41 | 36 | 24788917 | 30-51 | 41 | 36 |
| 109be_S6_R2_reverse | 24984267 | 10-51 | 41 | 36 | 24788917 | 30-51 | 41 | 36 |
| 109id_S7_R1_forward | 28319007 | 10-51 | 43 | 36 | 28107286 | 30-51 | 43 | 36 |
| 109id_S7_R2_reverse | 28319007 | 10-51 | 43 | 36 | 28107286 | 30-51 | 43 | 36 |
| 125be_S8_R1_forward | 28988546 | 10-51 | 41 | 36 | 28690426 | 30-51 | 41 | 36 |
| 125be_S8_R2_reverse | 28988546 | 10-51 | 41 | 36 | 28690426 | 30-51 | 41 | 36 |
| 127be_S9_R1_forward | 30708594 | 10-51 | 42 | 36 | 30400429 | 30-51 | 42 | 36 |
| 127be_S9_R2_reverse | 30708594 | 10-51 | 42 | 36 | 30400429 | 30-51 | 41 | 36 |
| 136be_S10_R1_forward | 24639894 | 10-51 | 42 | 36 | 24272120 | 30-51 | 42 | 36 |
| 136be_S10_R2_reverse | 24639894 | 10-51 | 42 | 36 | 24272120 | 30-51 | 42 | 36 |
| 145be_S11_R1_forward | 21782608 | 10-51 | 41 | 36 | 21566912 | 30-51 | 41 | 36 |
| 145be_S11_R2_reverse | 21782608 | 10-51 | 41 | 36 | 21566912 | 30-51 | 41 | 36 |
| 187be_S12_R1_forward | 31128143 | 10-51 | 40 | 36 | 30710885 | 30-51 | 40 | 36 |

| | | | | | | | | |
|---|---|---|---|---|---|---|---|---|
| 187be_S12_R2_reverse | 31128143 | 10-51 | 40 | 36 | 30710885 | 30-51 | 40 | 36 |
| 193be_S13_R1_forward | 20358971 | 10-51 | 41 | 36 | 20127343 | 30-51 | 41 | 36 |
| 193be_S13_R2_reverse | 20358971 | 10-51 | 41 | 36 | 20127343 | 30-51 | 41 | 36 |
| 193id_S14_R1_forward | 34638976 | 10-51 | 41 | 36 | 34302241 | 30-51 | 41 | 36 |
| 193id_S14_R2_reverse | 34638976 | 10-51 | 41 | 36 | 34302241 | 30-51 | 41 | 36 |
| 199be_S15_R1_forward | 25565010 | 10-51 | 41 | 36 | 25189404 | 30-51 | 41 | 36 |
| 199be_S15_R2_reverse | 25565010 | 10-51 | 41 | 36 | 25189404 | 30-51 | 40 | 36 |
| 199id_S16_R1_forward | 20532203 | 10-51 | 41 | 36 | 20251855 | 30-51 | 41 | 36 |
| 199id_S16_R2_reverse | 20532203 | 10-51 | 41 | 36 | 20251855 | 30-51 | 40 | 36 |
| 201be_S17_R1_forward | 72617679 | 10-51 | 41 | 36 | 72504922 | 30-51 | 41 | 36 |
| 201be_S17_R2_reverse | 72617679 | 10-51 | 41 | 36 | 72504922 | 30-51 | 40 | 36 |
| 201id_S18_R1_forward | 18796937 | 10-51 | 41 | 36 | 18550927 | 30-51 | 41 | 36 |
| 201id_S18_R2_reverse | 18796937 | 10-51 | 41 | 36 | 18550927 | 30-51 | 41 | 36 |
| 207be_S19_R1_forward | 17953289 | 10-51 | 41 | 36 | 7517461 | 30-51 | 41 | 36 |
| 207be_S19_R2_reverse | 17953289 | 10-51 | 41 | 36 | 7517461 | 30-51 | 41 | 36 |
| 207id_S20_R1_forward | 27465485 | 10-51 | 41 | 36 | 27465485 | 30-51 | 41 | 36 |
| 207id_S20_R2_reverse | 27465485 | 10-51 | 41 | 36 | 27465485 | 30-51 | 41 | 36 |

**Table S14.** Summary statistics for *de novo A. octoarticulatus* transcriptome assembly.

| Metric | Before redundancy reduction | After redundancy reduction |
|---|---|---|
| Transcripts | 98129 | 93122 |
| Transcripts > 500 bp | 58958 | 54250 |
| Transcripts > 1000 bp | 41277 | 37405 |
| Average length of assembled transcripts | 1,546.496 | 1,478.065 |
| Longest transcript | 27414 | 27414 |
| Total length | 151756069 | 137640410 |
| Transcript N50 | 3333 | 3232 |

https://figshare.com/s/40abd7858a4cabebe994

**Movie S1.** *Allomerus octoarticulatus* bodyguards attacking a grasshopper during a field bioassay on the ant-plant *Cordia nodosa* in Peru.
